# Glycosyltransferase POMGNT1 deficiency affects N-cadherin-mediated cell-cell adhesion

**DOI:** 10.1101/2020.09.09.289306

**Authors:** Sina Ibne Noor, Marcus Hoffmann, Natalie Rinis, Markus F. Bartels, Patrick Winterhalter, Christina Hoelscher, René Hennig, Nastassja Himmelreich, Christian Thiel, Thomas Ruppert, Erdmann Rapp, Sabine Strahl

## Abstract

Defects in protein *O*-mannosylation lead to severe congenital muscular dystrophies known as α-dystroglycanopathy. A hallmark of these diseases is the loss of the *O*-mannose-bound matriglycan on α-dystroglycan, which leads to a reduction in cell adhesion to the extracellular matrix. Mutations in protein *O*-mannose β1,2-N-acetylglucosaminyltransferase 1 (POMGNT1), which is crucial for the elongation of *O*-mannosyl glycans, are mainly associated with muscle-eye-brain (MEB) disease. In addition to defects in cell-extracellular matrix adhesion, aberrant cell-cell adhesion has occasionally been observed in response to defects in POMGNT1. However, direct molecular mechanisms are largely unknown. We used POMGNT1 knock-out HEK293T cells and fibroblasts from a MEB patient to gain a deeper insight into the molecular changes in POMGNT1 deficiency. A combination of biochemical and molecular biological techniques with proteomics, glycoproteomics and glycomics revealed that a lack of POMGNT1 activity strengthens cell-cell adhesion. We demonstrate that the altered intrinsic adhesion properties are due to an increased abundance of N-cadherin (N-Cdh). In addition, site-specific changes in the *N*-glycan structures in the extracellular domain of N-Cdh were detected, which positively impact on homotypic interactions. We found that in POMGNT1 deficient cells ERK1/2 and p38 signaling pathways are activated and transcriptional changes that are comparable to the epithelial-mesenchymal transition (EMT) are triggered, defining a possible molecular mechanism underlying the observed phenotype. Our study indicates that changes in cadherin-mediated cell-cell adhesion and other EMT-related processes may contribute to the complex clinical symptoms of MEB or α-dystroglycanopathy in general, and suggests a previously underestimated impact of changes in *O*-mannosylation on *N*-glycosylation.

## Introduction

The modification of proteins by glycosylation is an ubiquitous feature of all living organisms (1). Protein-linked glycans are involved in a multitude of cellular processes ranging from monitoring the folding state of glycoproteins to cell adhesion and migration (2). Among the different types of glycosylation, *N*-glycosylation and *O*-mannosylation are evolutionary conserved from bacteria to mammals. In humans, changes in those essential protein modifications *inter alia* can modulate immune responses, promote cancer cell metastasis and underlie the pathophysiology of severe congenital disorders (3–5). Both modifications initiate at the endoplasmic reticulum (ER), where the target polypeptides and the donor saccharides are synthesized and eventually covalently linked (2). Only properly glycosylated and folded proteins can leave the ER and travel through the Golgi apparatus to reach their final cellular destinations. On their way, *N*-linked and *O*-mannosyl glycans can be further modified which leads to diverse species-or even cell type-specific glycans (2).

In the case of *N*-glycosylation, the dolichol-pyrophosphate-linked oligosaccharide Glc_3_Man_9_GlcNAc_2_ is assembled at the ER membrane and the glycan moiety is transferred *en bloc* to Asn residues of the consensus sequon Asn-Xaa-Ser/Thr/Cys (X: proline is excluded). This way, the vast majority of proteins that enter the secretory pathway are *N*-glycosylated including many cell surface receptors and cell adhesion molecules (6). Protein-linked carbohydrate moieties are then further processed, and finally extended in the Golgi through the concerted action of diverse specific glycosyltransferases resulting in three distinct types of *N*-glycans: high-mannose, complex- and hybrid-type, which contain the common core Man3GlcNAc2-Asn (2). The diverse glycan structures and glycosylation patterns on cell surface molecules are highly dynamic and can be differentially regulated both during development and in certain pathological conditions, often associated with the acquisition of altered cellular properties (4).

Classically, *O*-mannosylation is initiated by the conserved PMT family of protein *O*-mannosyltransferases (POMT1 and POMT2 in mammals), which catalyse the transfer of mannose from dolichol-phosphate-linked mannose to Ser and Thr residues of nascent proteins (7). Three different core structures can be built on the protein-linked mannose (8). Linear core m1 and branched core m2 glycans, which share the common inner core GlcNAc-β1,2-Man-Ser/Thr, are initiated in the cis Golgi by the addition of a GlcNAc residue by the protein *O*-mannose β1,2-N-acetylglucosaminyltransferase 1 (POMGNT1), and are further extended while proteins travel through the Golgi to the cell surface. In contrast, core m3 glycans are already elongated in the ER (GalNAc-β1,3-GlcNAc-β1,4-(phosphate-6)-Man-Ser/Thr) and then further modified in the Golgi by the sequential action of numerous glycosyltransferases including the ribulose-5-phosphate transferase fukutin (FKTN). The resulting complex polysaccharide structure, known as “matriglycan” is so far only found on α-dystroglycan (α-DG), a central member of the dystrophin glycoprotein complex family in peripheral membranes, and enables its interaction with extracellular matrix (ECM) components such as laminin (9). Defects in this complex biosynthetic pathway lead to the loss of the matriglycan on α-DG, and consequently impair interactions between α-DG and e.g. laminin, which interferes with the formation of basement membranes (9). This defect has been recognized as a major patho-mechanism of severe congenital muscular dystrophies with neuronal migration defects, known as α-dystroglycanopathy (OMIM 236670; 253280; 253800; 606612; 607155; 608840) (10).

The glycosyltransferase POMGNT1 has a key role in the elongation of *O*-mannosyl glycans (11). In its absence not only core m1 and m2 structures are missing, also formation of the matriglycan fails, since POMGNT1 recruits FKTN to maturing core m3 structures (12, 13). The great majority of mutations in *POMGNT1* have been linked to muscle-eye-brain disease (MEB; OMIM 253280), a congenital muscular dystrophy in humans, which is characterized by additional brain malformations and structural anomalies in the eye (11). In the murine model, knock-out of *POMGNT1* is viable with multiple developmental defects, similar to the clinical picture of human MEB patients (14, 15). The pathology of MEB suggests a functional role for POMGNT1 in control of cell adhesion and migration. For example, in the transgenic *POMGNT1*-based MEB mouse model impaired cell-ECM adhesion results in disruption of basement membranes and over migration of neurons during development of the cerebral cortex (15). However, also clusters of granule cells which failed to migrate have been frequently observed (15). In addition to its important role during mammalian development, POMGNT1 has recently been linked to the progression of glioblastoma, fatal primary brain tumors with survival time of 12-15 months, as well as the resistance of glioblastoma cells to the chemotherapeutic agent temozolomide (16, 17). Strikingly, in glioblastoma models increased cell-cell adhesion has been observed when POMGNT1 is missing (16). However, molecular reasons for the different consequences of POMGNT1 deficiency are just emerging.

Very recently, glyco-engineered human embryonic kidney (HEK) 293 cells turned out to be especially useful for the characterization of known, as well as the identification of new glycosylation pathways (18, 19). In the present work, we took advantage of a gene-targeted *POMGNT1* knock-out in HEK293T cells to study the consequences of POMGNT1 deficiency. The combination of glyco(proteo)mics with classic biochemistry, molecular and cell biology resulted in the discovery that cell-cell adhesion mediated by neuronal cadherin (N-Cdh) is affected and defined a possible molecular mechanism underlying the observed phenotype. Similar effects in MEB patient-derived fibroblasts confirmed the validity of the HEK293T model to study molecular effects of *O*-mannosylation deficiencies.

## Results

### POMGNT1 deficiency impairs cell-matrix and reinforces cell-cell interactions in a HEK293T cell model

To gain insight into functional implications of *POMGNT1* deficiency, we generated a gene-targeted knock-out in HEK293T cells (ΔPOMGnT1) as detailed in Experimental procedures and **Fig. S1**. The loss of POMGNT1 activity was confirmed by the *O*-mannosylation status of the endogenous substrate α-DG using the matriglycan-directed antibody IIH6. Whereas the *O*-linked matriglycan is absent in *POMGNT1*-depleted cells, reintroduction of human *POMGNT1* rescued *O*-mannosylation of α-DG verifying the specificity of the system (**Fig. 1A**). General characterization of the morphology of *POMGNT1* knock-out cells by confocal microscopy revealed that *POMGNT1*-deficient cells appear more rounded and stronger aggregated compared to wild type cells which show extensive spreading and even distribution. This phenotype is also reverted upon reintroduction of *POMGNT1* (**Fig. 1B**).

**Figure 1.**
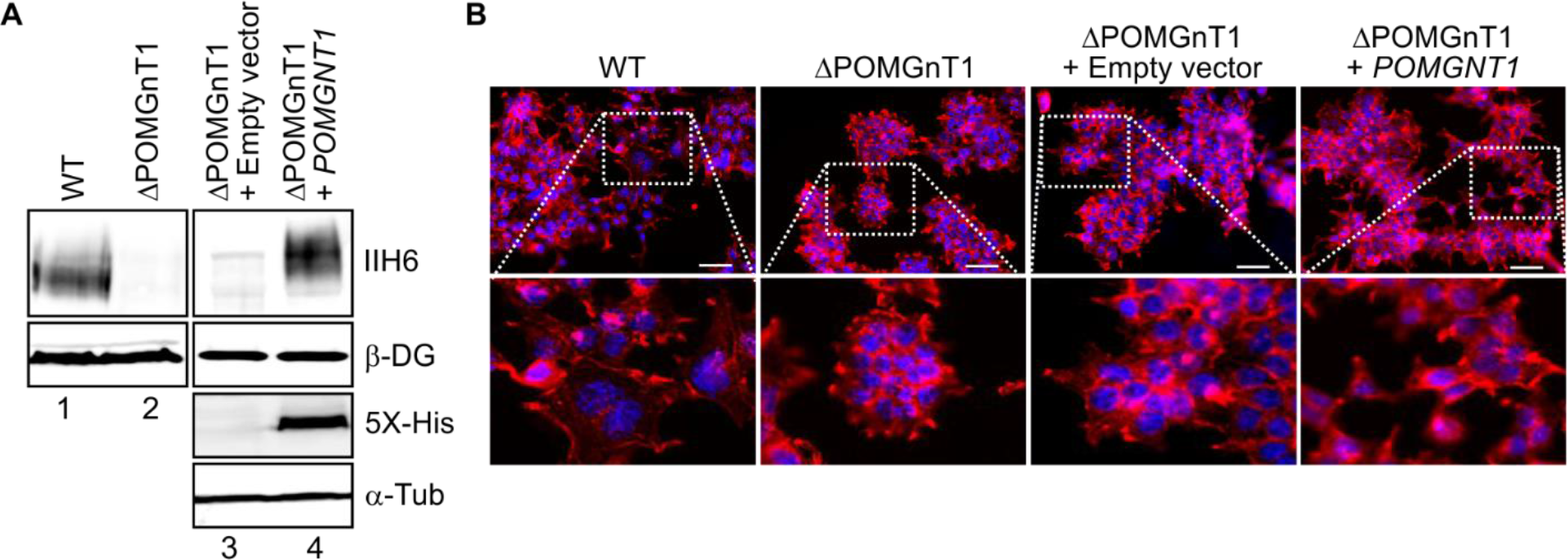
Characterization of *POMGNT1* knock-out HEK293T cells. (A, B) WT and ΔPOMGnT1 cells with or without *POMGNT1* complementation are shown. ΔPOMGnT1 cells were stably transfected with either empty vector (ΔPOMGnT1 + empty vector) or vector carrying *POMGNT1* (ΔPOMGnT1 + *POMGNT1*). (A) Western Blot analysis of α-DG from WGA-enriched fractions (first and second panel). Genome targeting of *POMGNT1* results in loss of its enzymatic activity as demonstrated by the lack of matriglycan on α-DG (as identified by IIH6 immuno-reactivity) (compare lanes 1 and 2). *O*-mannosylation of α-DG is restored upon reintroduction of *POMGNT1* (compare lanes 2 and 4). Analysis of whole cell lysates with anti-5x-His antibody shows His-tagged POMGNT1 in ΔPOMGnT1 cells (third and fourth panel). β-DG served as a loading control for the WGA-enriched fractions and α-Tubulin (α-Tub) for the whole cell lysates. (B) Confocal images displaying differences in cell-cell adhesion among WT, ΔPOMGnT1 and *POMGNT1*-transfected ΔPOMGnT1 cells. In the upper panels, cells fixed and stained with Rhodamine Phalloidin (red; actin filaments) and DAPI (blue; nucleus) after three days of seeding are shown. In the lower panels, the area of interest of each image is magnified. The scale bar represents 10 μm.

To further characterize molecular events responsible for the morphological differences, we analyzed cell-matrix and cell-cell adhesion. As expected, *POMGNT1*-deficient cells adhere to laminin, a major ECM component and interactor of the α-DG matriglycan, to a significantly lower extent when compared to wild type cells (**Fig. 2A**). Intriguingly, when confluent monolayers of wild type cells were incubated with wild type and knock-out cells, respectively, cell-cell adhesion of ΔPOMGnT1 cells turned out to be significantly increased. The same result is observed using a monolayer of ΔPOMGnT1 cells (**Fig. 2B**). Since cell-matrix and cell-cell interactions are major opposing forces balancing cellular migration, we further took advantage of xCELLigence real-time cell analysis that allows live monitoring of cell proliferation and cell migration. ΔPOMGnT1 cells proliferate slower than wild type cells with slopes of 0.07 and 0.09, respectively (**Fig. S2, A** and **B**). In agreement with increased cell-cell adhesion, the migration rate of ΔPOMGnT1 cells is reduced by a factor of three (**Fig. 2C** and **Fig. S2D**).

**Figure 2.**
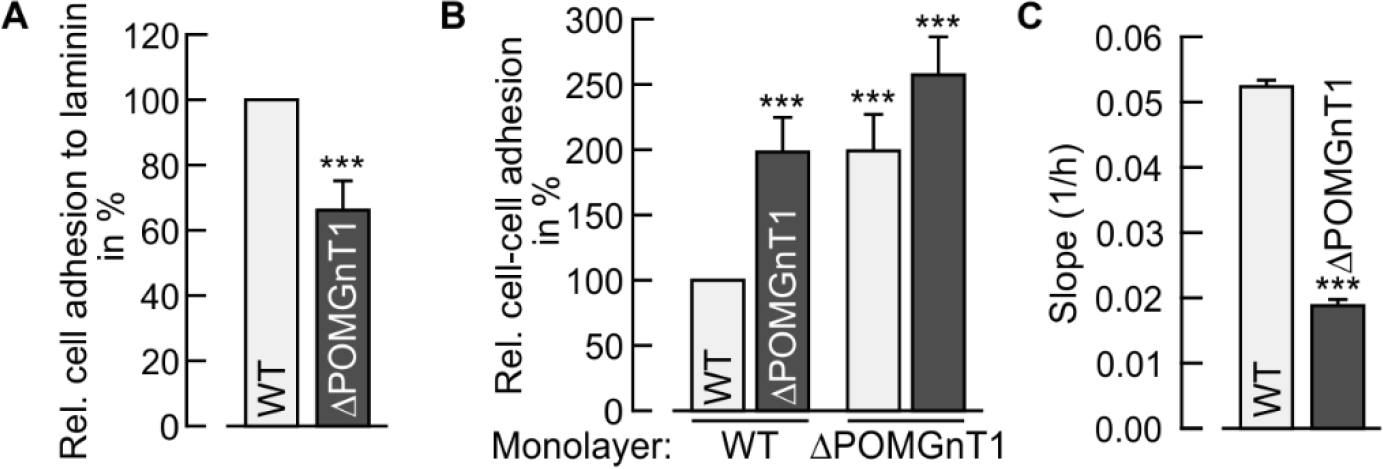
*POMGNT1* deficiency affects cell adhesion and migration in HEK293T cells. (A) Relative adhesion of WT and ΔPOMGnT1 cells to the ECM component laminin. WT or ΔPOMGnT1 cells deprived of FBS were seeded on laminin or BSA (as a control) pre-coated wells and allowed to adhere for 1 h at 37°C. Adherent cells were stained with crystal violet and quantified at OD_600_ after stain extraction. The absorbance for each laminin coated well was normalized against the mean absorbance of BSA-coated wells. Adhesion of ΔPOMGnT1 cells to laminin is represented as relative adhesion in %, considering adhesion of WT cells to laminin as 100%. (B) Relative cell-cell adhesion of WT and ΔPOMGnT1 cells to cell monolayers as indicated. WT or ΔPOMGnT1 cells deprived of FBS were seeded on confluent monolayers of cells with either the same or different genotype and allowed to adhere for 20 min at 37°C. Adherent cells were stained with crystal violet and quantified at OD_600_ after stain extraction. The absorbance of each well to which cells were added was normalized against the mean absorbance of wells, where no cells were added. Respective cell-cell adhesion is represented as relative adhesion in %, considering adhesion of WT to WT cells as 100%. (C) Comparison of the migration rate between WT and ΔPOMGnT1 cells. WT and ΔPOMGnT1 cells deprived of FBS were seeded on xCELLigence microelectrode-equipped trans-well chambers and allowed to migrate for 90 h at 37°C. Migration was monitored in real-time as increase of impedance and was expressed as Cell Index, an arbitrary unit. The migration rate was calculated from the slope of the curves between 20 h and 60 h (Fig. S2D). (A-C) Assays were performed at least in triplicate from two independent experiments. Data are represented as means ± SD (Standard Deviation). Asterisks denote statistical significance in comparison to WT cells: *** *p* ≤ 0.001.

Taken together, the *POMGNT1* knock-out HEK293T cell model revealed that cell-cell adhesion increases, whereas cell-matrix interactions and cell migration are negatively affected when *O*-mannosyl glycans are not further elongated.

### Increased cell-cell adhesion of POMGNT1-deficient cells is caused by changes in N-Cdh abundance

In order to identify determinants which underlie the observed phenotype in ΔPOMGnT1 cells, we performed label-free quantitative proteomics of whole cell lysates from wild type and ΔPOMGnT1 HEK293T cells. Five independent replicates were analyzed and homoscedasticity and normal distribution were confirmed (**Fig. S3, A** and **B**). Altogether, 86 out of 437 proteins with differential abundance in *POMGNT1-deficient* cells could be identified (**Fig. 3A** and **Fig. S3D, Table S4**). Interestingly, gene ontology term functional annotation of proteins with a significant regulation revealed enrichment for proteins under the molecular function term of “cadherin binding involved in cell-cell adhesion” (**Fig. 3A** and **S3D, Table S4**), pointing to an impact of POMGNT1 deficiency on cadherin-mediated cell-cell adhesion. In addition, the protein N-Cdh was found to be increased by a factor of three in ΔPOMGnT1 cells (**Fig. 3A**). This result was also confirmed by Western blot (**Fig. 3, B** and **C**) and correlated well with increased mRNA levels of N-Cdh (**Fig. 3D**). To investigate the general validity of our findings, we took advantage of skin fibroblasts derived from an MEB patient who presented characteristic symptoms such as mental retardation and blindness due to variant c.535_751del (p.Asp179Argfs* 11) in the *POMGNT1* gene (NM_017739.4). In accordance with our HEK293T model, protein and mRNA abundance of N-Cdh in MEB patient-derived fibroblasts showed increased values compared to fibroblasts from two healthy donors (**Fig. 3, E-G**).

**Figure 3.**
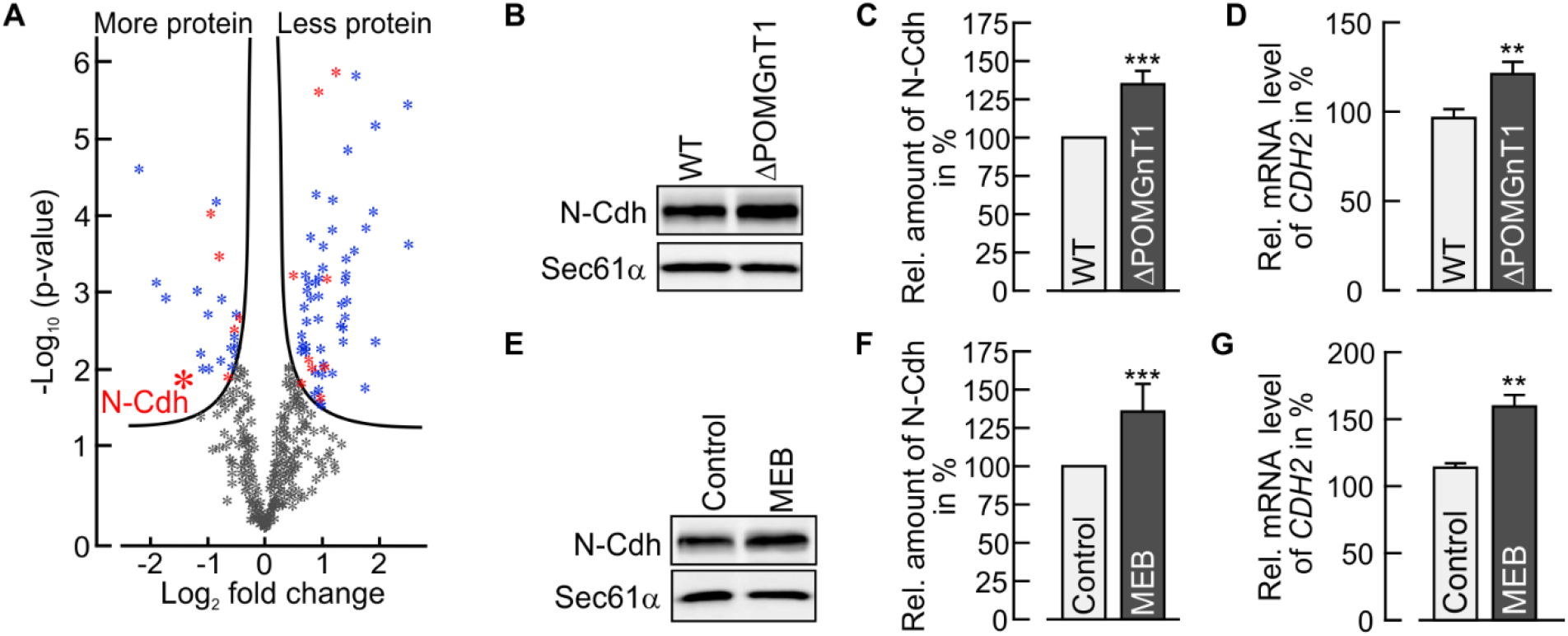
Defects in *POMGNT1* induce expression of N-Cadherin. (A) Volcano plot of differentially expressed proteins of WT and ΔPOMGnT1 cells identified by label-free LC-MS/MS of five biological replicates each. The solid lines represent the threshold of statistical significance with false discovery rate (FDR) < 0.05 and an s0 of 0.1 (controlling the relative importance of t-test p-value and the difference between means. The complete list of proteins is included in Table S4. Proteins with a significant fold change are depicted in blue dots, with a subset of Cadherin-associated proteins in red dots. Gray dots indicate proteins that failed to meet the criteria of FDR or fold change. (B, E) Representative Western Blot used for N-Cdh protein quantification in 20 μg of membrane fractions from WT and ΔPOMGnT1 cells (B) and from control and MEB patient-derived fibroblasts (E). Sec61α was used as loading control. (C, F) Densitometry-based quantification of N-Cdh signal detected by Western Blot in WT and ΔPOMGnT1 cells (C) and in control and MEB patient derived fibroblasts (F). N-Cdh signals for ΔPOMGnT1 cells and MEB patient-derived fibroblasts were normalized to the respective Sec61α signal and subsequently normalized to the N-Cdh/Sec61α ratio calculated for WT cells and control fibroblasts, respectively. Protein levels of N-Cdh are represented in %, considering pixel density of N-Cdh in WT cells and MEB patient-derived fibroblasts as 100%, respectively. Assays were performed with three biological replicates. (D, G) qRT-PCR analysis of *CDH2* mRNA levels in WT and ΔPOMGnT1 cells (D) and in control and MEB patient-derived fibroblasts (G). *CDH2* mRNA levels in ΔPOMGnT1 cells and MEB patient-derived fibroblasts are calculated as %, considering *CDH2* mRNA levels in WT cells and MEB patient derived fibroblasts as 100%, respectively. For normalization the housekeeping gene hypoxanthine-guanine phosphoribosyltransferase (HPRT) was used. Assays were performed in duplicate with three dilutions from three biological replicates. All data are presented as means ± SD. Asterisks denote statistical significance in comparison to WT cells: ** *p* ≤ 0.01, *** *p* ≤ 0.001.

Cadherins are major players in the formation of cellular junctions (20). These membrane-anchored cell surface glycoproteins mediate cell-cell adhesion through homotypic interactions of their conserved extracellular domains. Thus, to determine whether elevated cell-cell adhesion observed in *POMGNT1*-deficient cells is directly linked to the increased abundance of N-Cdh, we performed cell-cell adhesion assays in presence of either N-Cdh blocking or IgG-directed antibodies. As shown in **Fig. 4**, ΔPOMGnT1 cells show increased adhesion to wild type and other ΔPOMGnT1 cells in the IgG-treated controls. This increase, however, is diminished upon incubation with an N-Cdh targeting antibody, demonstrating N-Cdh as a key molecular driver for increased cell-cell adhesion in the established ΔPOMGnT1 HEK293T cell model.

**Figure 4.**
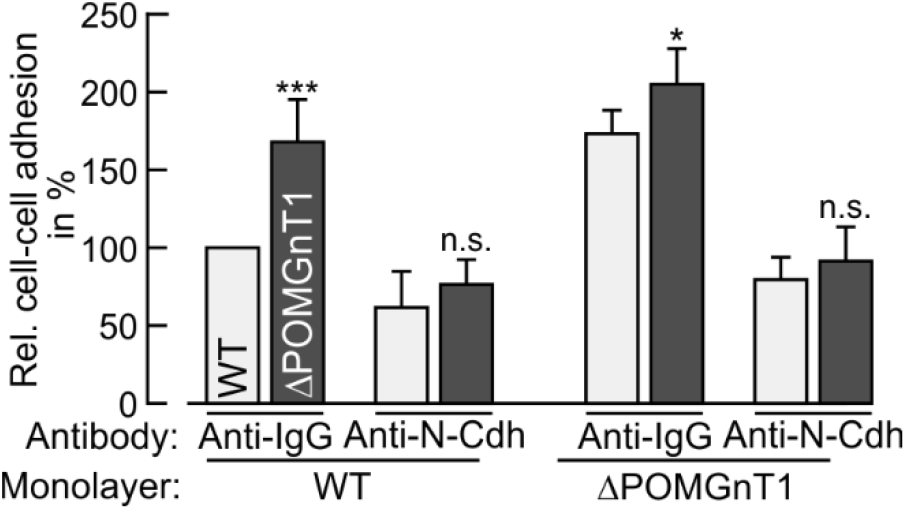
Increased cell-cell adhesion of *POMGNT1*-deficient cells is mediated by increased abundance of N-cadherin. Relative cell-cell adhesion of WT (bright bars) and ΔPOMGnT1 (dark bars) cells to the indicated cell monolayers in presence of either N-Cdh targeting or anti-human IgG antibody. WT or ΔPOMGnT1 cells that were deprived of FBS, were incubated with an anti-N-Cdh antibody and subsequently seeded on an anti-N-Cdh antibody-treated confluent monolayer of cells. Anti-human IgG antibody served as a control. Cells were allowed to adhere for 20 min at 37 °C. Adherent cells were stained with crystal violet and quantified at OD600 after stain extraction. The absorbance of each well to which cells were added was normalized against the mean absorbance of wells, where no cells were added. Respective cell-cell adhesion is represented as relative adhesion in %, considering relative adhesion of WT to WT cells pretreated with anti-human IgG antibody as 100%. Assays were performed in triplicate from two independent experiments. All data are presented as means ± SD. Asterisks denote statistical significance in comparison to WT cells: * *p* ≤ 0.05, *** *p* ≤ 0.001, *n.s*. not significant.

### POMGNT1 deficiency affects N-Cdh N-glycosylation

N-Cdh is highly *N*-glycosylated and *N*-linked glycans affect its adhesive properties by modulating its homomeric interactions in cis and in trans (21, 22). We therefore asked whether an altered *N*-glycosylation, as an indirect effect of POMGNT1 deficiency, could contribute to increased N-Cdh-mediated cell-cell adhesion. We performed comprehensive *N*-glycomics and *N*-glycoproteomics on N-Cdh. For that purpose, the extracellular domain (EC) of N-Cdh was recombinantly expressed and purified from wild type and ΔPOMGnT1 cells (detailed in Experimental procedures; **Fig. S4**). *N*-glycomics analysis by multiplexed capillary gel electrophoresis with laser induced fluorescence detection (xCGE-LIF) revealed that total abundance of fully galactosylated and sialylated *N*-glycan structures is remarkably decreased on N-Cdh derived from ΔPOMGnT1 cells when compared to wild type (**Fig. 5A**, normalized intensity of peaks 2, 4, 9 and 10). In accordance, the abundance of non-galactosylated *N*-glycan structures is increased (**Fig. 5A**, normalized intensity of peaks 6 and 7). Moreover, fully galactosylated multiantennary *N*-glycans (**Fig. 5A**, A3+F) are reduced on ΔPOMGnT1-derived N-Cdh and a non-galactosylated complex-type *N*-glycan with a bisecting GlcNAc (GlcNAc(5)Man(3)+Fuc(1)) represents the dominating *N*-glycan structure (detailed in Supporting information **SI5**).

**Figure 5.**
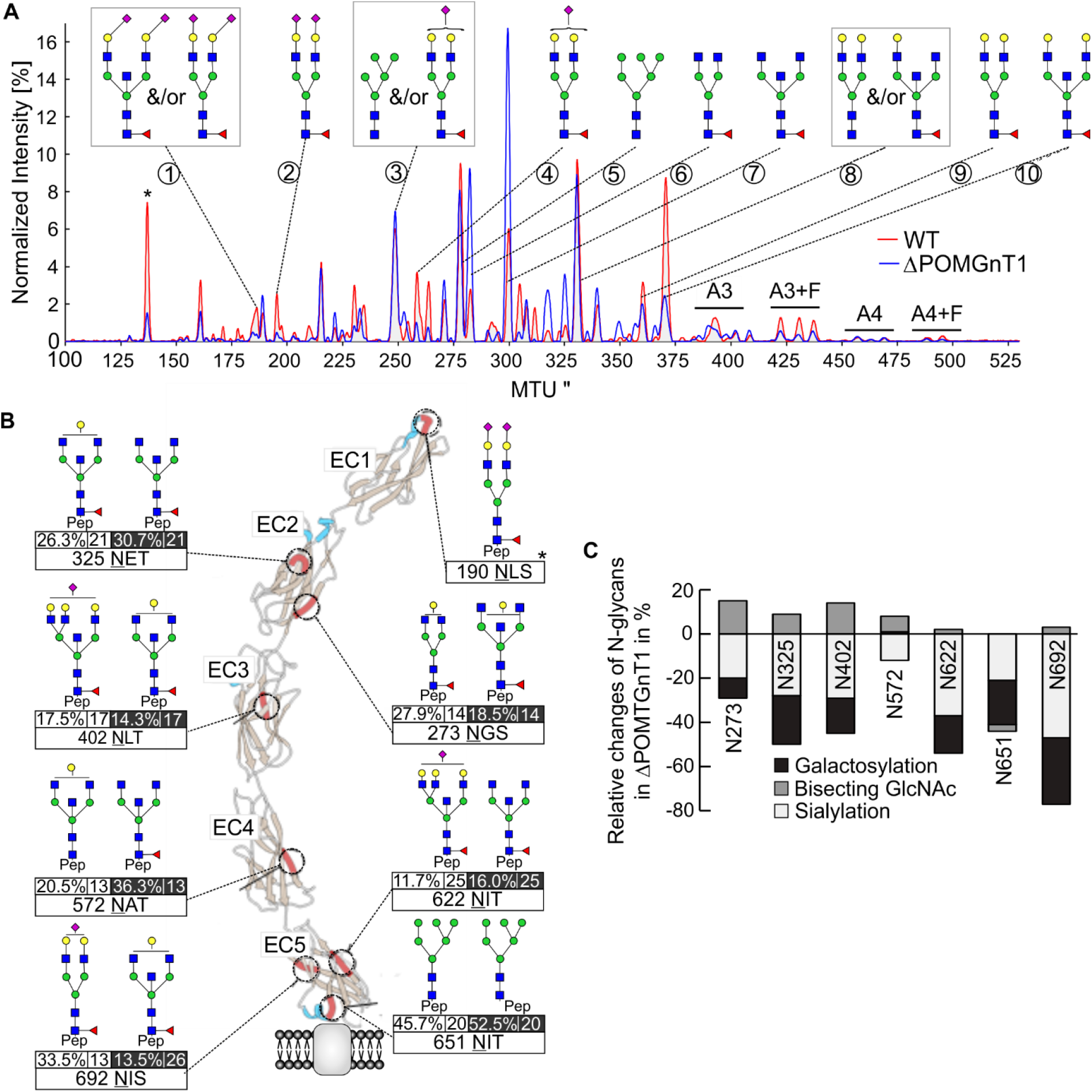
Deficiency in POMGNT1 affects *N*-glycosylation of N-cadherin. (A) Comparative overlay of the electropherograms as obtained by xCGE-LIF after APTS labeling to evaluate *N*-glycan profile of N-Cdh derived from WT (red line) and ΔPOMGnT1 (blue line) HEK293T cells. The normalized signal intensity of APTS-labeled *N*-glycans (relative signal intensity (%), i.e. peak signal divided by the summed peak height of all quantifiable *N*-glycan peaks with signal-to-noise-ratio >9) is plotted over their normalized migration time units (MTU”). Relevant peaks are numbered and annotated with corresponding glycan structures. A3 and A4 indicate three- and four-antennary *N*-glycan structures, respectively; while F indicates the presences of a fucose among the *N*-glycan structures. The asterisk denotes a peak of an internal migration time standard. (B) Overview of site-specific *N*-glycan microheterogeneity on extracellular domains (EC1-5) of N-Cdh, purified from WT and ΔPOMGnT1 HEK293T cells. Cartoon represents the molecular structure of five EC domains (EC1-EC5) of N-Cdh, which form a β-barrel structure (brown and blue ribbons represent beta sheet and alpha helix, respectively). The position of identified *N*-glycosylation site are shown in red ribbons. For each *N*-glycosylation site, the major *N*-glycoform detected in WT (left) and ΔPOMGnT1 (right) is depicted. Values beneath the each glycan structures represent the relative abundance of this *N*-glycoform (quantitative values for each glycoform as proportions relative to the sum of all glycoforms detected) and the total of *N*-glycoforms detected on this site, respectively. Values for ΔPOMGnT1 are highlighted in black. Asterisk at position Asn190 indicates the scarcely *N*-glycosylated site (only one *N*-glycopeptide derived from WT N-Cdh could be identified). The molecular structure of N-Cdh was modeled by using open source software UCSF Chimera Version 1.10.2. (A, B) *N*-glycan structures were drawn with GlycoWorkbench Version 1.1, by following the guideline of the Consortium for Functional Glycomics. Pep = peptide moiety; Green circle = mannose; yellow circle = galactose; blue square = N-acetylglucosamine; red triangle = fucose; purple diamond = N-acetylneuraminic acid. (C) Site-specific relative changes in the *N*-glycan traits (bisecting GlcNAc, sialylation and galactosylation) in ΔPOMGnT1-derived N-Cdh. For each *N*-glycosylation site, quantitative changes in the *N*-glycan microheterogeneity are depicted as increase or decrease in ΔPOMGnT1-derived N-Cdh relative to the level in the WT. The quantities represent *N*-glycopeptide AUC values (EIC MS1) that were summed based on common *N*-glycan traits (*N*-glycopeptides carrying a bisecting GlcNAc, sialylation and/or galactosylation). Since site N190 is only scarcely *N*-glycosylated, its microheterogeneity could not be quantified.

In order to gain even deeper insights, an exploratory site-specific approach was taken to comprehensively map glycosylation sites and the corresponding *N*-glycoforms of N-Cdh. Hydrophilic interaction liquid chromatography (HILIC)-enriched tryptic and proteinase K-generated N-Cdh *N*-glycopeptides were analyzed by nano-reverse-phase liquid chromatography coupled online to an electrospray ionization orbitrap tandem mass spectrometer (nano-RP-LC-ESI-OT MS/MS) (for details see Experimental procedures). All eight potential *N*-glycosylation sites (Asn190, 273, 325, 402, 572, 622, 651, 692) of N-Cdh were identified (**Fig. 5B**, indicated as red ribbons). With the exception of Asn651 which predominantly carries a high-mannose-type *N*-glycan, all sites feature complex-type *N*-glycans. Characterization of the *N*-glycan microheterogeneity based on intact *N*-glycopeptides revealed differences in relative abundance of the major *N*-glycoform at each site on N-Cdh derived from ΔPOMGnT1 compared to wild type cells (**Fig. 5C**). All *N*-glycosylation sites (except for Asn651 and Asn692 which are both located at the stem region of the molecule) feature non-galactosylated complex-type *N*-glycans with a bisecting GlcNAc (GlcNAc(5)Man(3)±Fuc(1)) as the dominating composition on N-Cdh from ΔPOMGnT1 cells. Comparative and site-specific *N*-glycoproteomics of quantitative changes in the *N*-glycan microheterogeneity revealed a significant decrease in galactosylation (−27% on average) and sialylation (−16% on average) throughout the vast majority of N-glycosylation sites (only exception are Asn190 and Asn651) (**Fig. 5C**). This decrease in galactosylation and sialylation is associated with a significant increase of non-galactosylated *N*-glycan compositions that feature a bisecting GlcNAc (**Fig. 5B**). The site-specific relative changes of *N*-glycan traits on N-Cdh are shown in detail in Supporting information **SI1-SI3**. Overall, *N*-glycosylation of N-Cdh in ΔPOMGnT1 cells exhibits a reduction in the degree of galactosylation and sialylation with non-galactosylated complex-type *N*-glycans with a bisecting GlcNAc dominating.

We next measured the transcript levels of genes encoding for major glycosyltransferases of *N*-glycan processing that could be responsible for the observed changes in the *N*-glycan profile of N-Cdh using the nCounter technology (23). A decrease by 27% of β-1,4-galactosyltransferase 1 *B4GALT1* mRNA as well as a reduction by 83 % in the transcript level of sialyltransferase *ST6GAL1* was detected, explaining the limited occurrence of respective *N*-glycans on N-Cdh in ΔPOMGnT1 cells (**Table S6**). Further, mild reduction in the mRNA levels of the Golgi GlcNAc-transferases *MGAT1* (by 17%), *MGAT2* (by 13%) and *MGAT5* (by 19%) were observed.

In summary, our data demonstrate that in *POMGNT1*-deficient cells the *N*-glycan profile of N-Cdh is changed and that the observed alterations correlate to a great extent with transcriptional changes of respective *N*-glycan modifying enzymes.

In addition to *N*-glycans, N-Cdh carries non-elongated *O*-linked mannose residues (24, 25) which depend on a recently identified class of *O*-mannosyltransferases (tetratricopeptide repeats (TPR)-containing proteins (TMTC 1-3), rather than classic *O*-mannosylation (18). The *O*-mannose glycoproteome of N-Cdh revealed 15 *O*-mannose glycosylation sites – four of which have not been reported before (Supporting information **SI4**). However, no difference in the *O*-mannosylation pattern of N-Cdh between ΔPOMGnT1 and wild type HEK293T cells was detected, excluding indirect effects on the activity of TMTCs.

### Changes in N-Cdh N-glycosylation impact on its homotypic interactions in vitro

Hypo-*N*-glycosylation of N-Cdh increases the prevalence of cis N-Cdh dimers on the cell membrane, thereby stabilization of cell-cell contacts (21, 22). To determine whether the observed changes in *N*-glycosylation of N-Cdh also favor homotypic interactions, the adhesive properties of the recombinant extracellular domain of N-Cdh (see above) were analyzed using an *in vitro* bead aggregation assay (26). Dynabeads coated with the protein isolated from wild type or ΔPOMGnT1 HEK293T cells were incubated with either CaCl_2_ or EDTA as a control. Aggregation of beads, reflecting Ca^2+^-dependent homotypic interactions of the extracellular domain of N-Cdh, was recorded and quantified at specific time intervals over the course of 1 hour. As shown in **Fig. 6A**, ΔPOMGnT1-derived N-Cdh triggers the formation of larger aggregates in comparison to wild type-derived N-Cdh, demonstrating that the adhesive properties of N-Cdh from ΔPOMGnT1 cells are enhanced. Taken together, changes in cell-cell adhesion of ΔPOMGnT1 cells are due to a higher frequency of N-Cdh with less complex *N*-linked glycan structures that facilitate stronger homotypic interactions.

**Figure 6.**
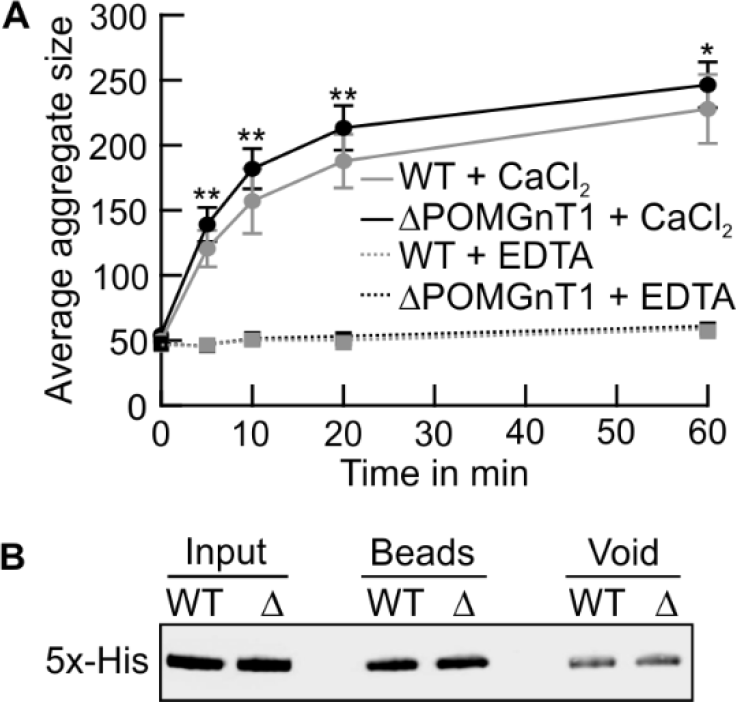
N-Cadherin from *POMGNT1-deficient* cells shows enhanced homotypic interactions *in vitro*. (A) Semi-quantitative analysis of the mean size of dynabead aggregates coated with the recombinant extracellular domain of recombinant N-Cdh purified from WT (gray line) and ΔPOMGnT1 cells (black line). Protein-coated dynabeads were imaged following incubation in the presence of either CaCl_2_ (solid lines) or EDTA (broken lines) at the indicated time intervals. Mean aggregate size is plotted against time. Assays were performed at quadruplicate from two independent experiments. Data are represented as means ± SD. Asterisks denote statistical significance in comparison to WT cells: * *p* ≤ 0.05, ** *p* ≤ 0.01. (B) Western Blot analysis evaluating levels of protein from WT and ΔPOMGnT1 (Δ) bound to dynabeads. Dynabead-bound recombinant N-Cdh was eluted in SDS sample buffer (beads), resolved on a 10% SDS PAA gel. For detection on Western blot an anti-5xHis antibody was used. In addition, protein before binding to beads (input) and unbound fractions (void) are shown.

### Loss of function of POMGNT1 activates ERK and p38 signaling

To further investigate how POMGNT1 deficiency modulates N-Cdh-mediated cell-cell adhesion, we explored possible changes in cellular signaling using a human phosphokinase array with 46 different kinases. Of all kinases examined, the extracellular signal-regulated kinase ERK1/2 and the mitogen-activated protein kinase p38 showed significantly enhanced phosphorylation in ΔPOMGnT1 HEK293T cells when compared to wild type cells (**Fig. 7, A** and **B**). Since dysregulation of DG and matriglycan biosynthesis has been associated with modulation of the ERK-MAPK pathway (27, 28), the identified changes were further validated by Western Blot. Consistent with the phospho-kinase array data, ΔPOMGnT1 cells showed a significant increase in the phosphorylation levels of ERK1/2. Additionally, phosphorylation of the upstream MAP kinase kinase MEK1/2 (29) was found to be increased (**Fig. 7, C** and **D**) further corroborating that the ERK-MAPK pathway is activated in our ΔPOMGnT1 HEK293T cell model.

**Figure 7.**
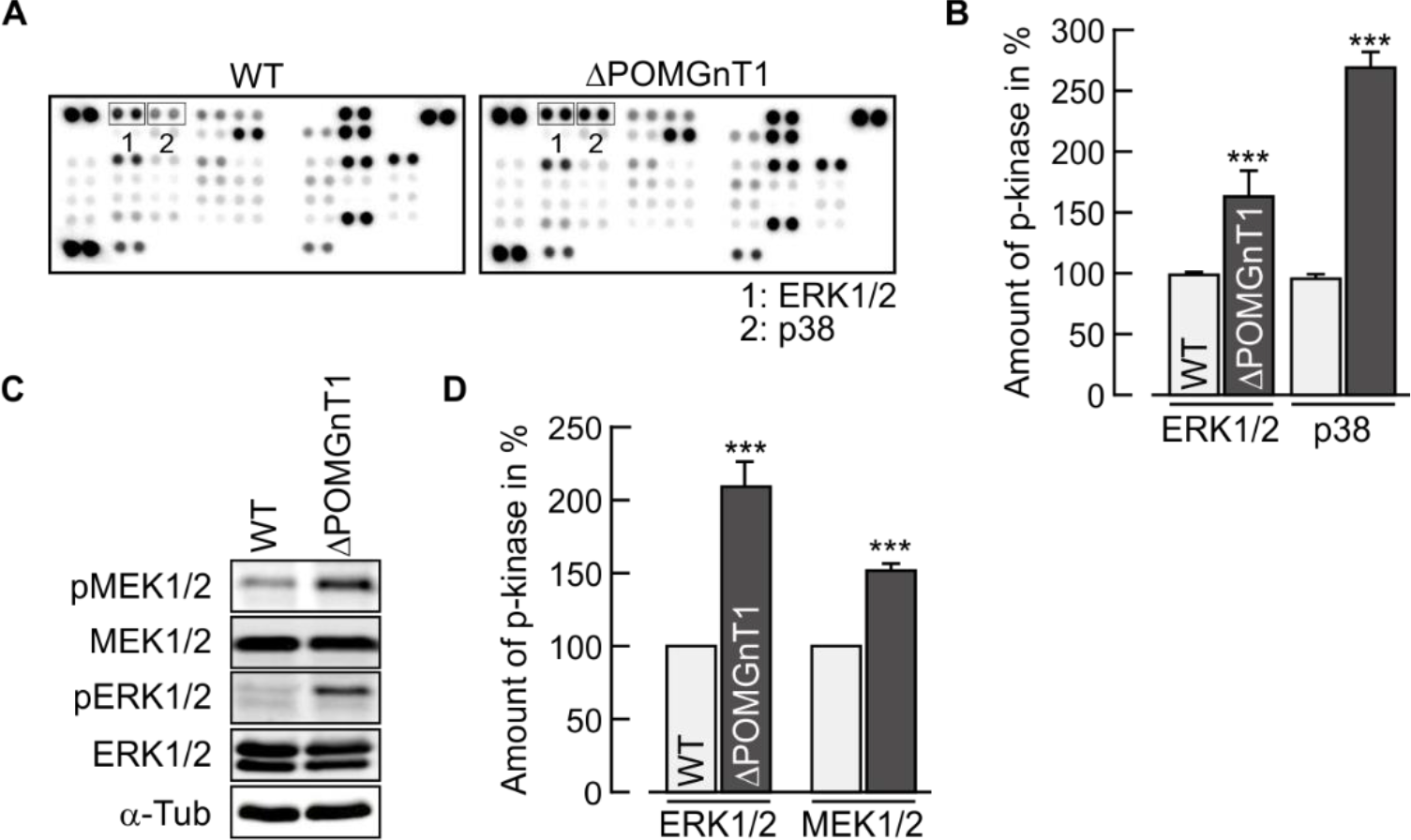
*POMGNT1* deficiency activates ERK1/2 and p38 signaling pathways. (A) Representative phospho-kinase array performed with 500 μg of whole cell extract from WT (left panel) and ΔPOMGnT1 cells (right panel). Antibodies against phosphorylated protein kinases are spotted in duplicate. Signals for phosphorylated ERK1/2 and p38 are highlighted in boxes and denoted as 1 and 2, respectively. (B) Densitometry-based quantification of phosphorylated ERK1/2 and p38 signals detected in phospho-kinase array for WT and ΔPOMGnT1 cells. Protein signals detected in ΔPOMGnT1 cells were normalized to the reference protein signal and subsequently normalized to the protein/ reference signal ratio in WT cells. Analysis is based on two independent experiments. (C) Representative Western Blot analysis of p (phosphorylated)-ERK1/2 and MEK1/2 levels in 20 μg of whole cell extract from WT and ΔPOMGnT1 cells. Total levels of ERK1/2 and MEK1/2 were analyzed using protein-directed antibodies. α-Tubulin (α-Tub) was used as a loading control. (D) Densitometry-based quantification of phosphorylated ERK1/2 and MEK1/2 signals detected in Western Blot analysis for WT and ΔPOMGnT1 cells. Phosphorylated protein signals detected in ΔPOMGnT1 cells were normalized to the respective total protein signals and subsequently normalized to the phosphorylated/ total protein signal ratio in WT cells. Data of three technical replicates are shown. (B, D) Protein levels of phosphorylated kinases in ΔPOMGnT1 cells is calculated as %, considering pixel density of the respective protein in WT cells as 100%. All data are presented as means ± SD. Asterisks denote statistical significance in comparison to WT cells: *** *p* ≤ 0.001.

### Lack of POMGNT1 activity induces an EMT-related transcriptional response

Increased expression of N-Cdh (30), transcriptional modulation of *N*-glycan modifying enzymes (31) and activation of the ERK-MAPK pathway (32) have been linked to epithelial-mesenchymal transition (EMT), a process which is of major importance during development, wound healing as well as tumor progression and metastasis (33). Therefore, the impact of POMGNT1 deficiency on the expression of EMT marker genes was investigated. As shown in **Fig. 8A**, the mRNA levels of the epithelial markers *LAMA2* (encoding laminin subunit α2) and *CLDN6* (encoding claudin 6) are markedly decreased, whereas expression levels of the mesenchymal markers *ACTA2* (encoding actin α2) and *CDH2* (encoding N-Cdh) are significantly elevated in ΔPOMGnT1 compared to wild type cells. Reintroduction of the *POMGNT1* gene reversed the observed transcriptional changes thereby confirming their specificity. Moreover, mRNA levels of the EMT-inducing transcription factors (*Slug, Snail, TCF3, LEF1*) showed significant upregulation in ΔPOMGnT1 cells (**Fig. S5**).

**Figure 8.**
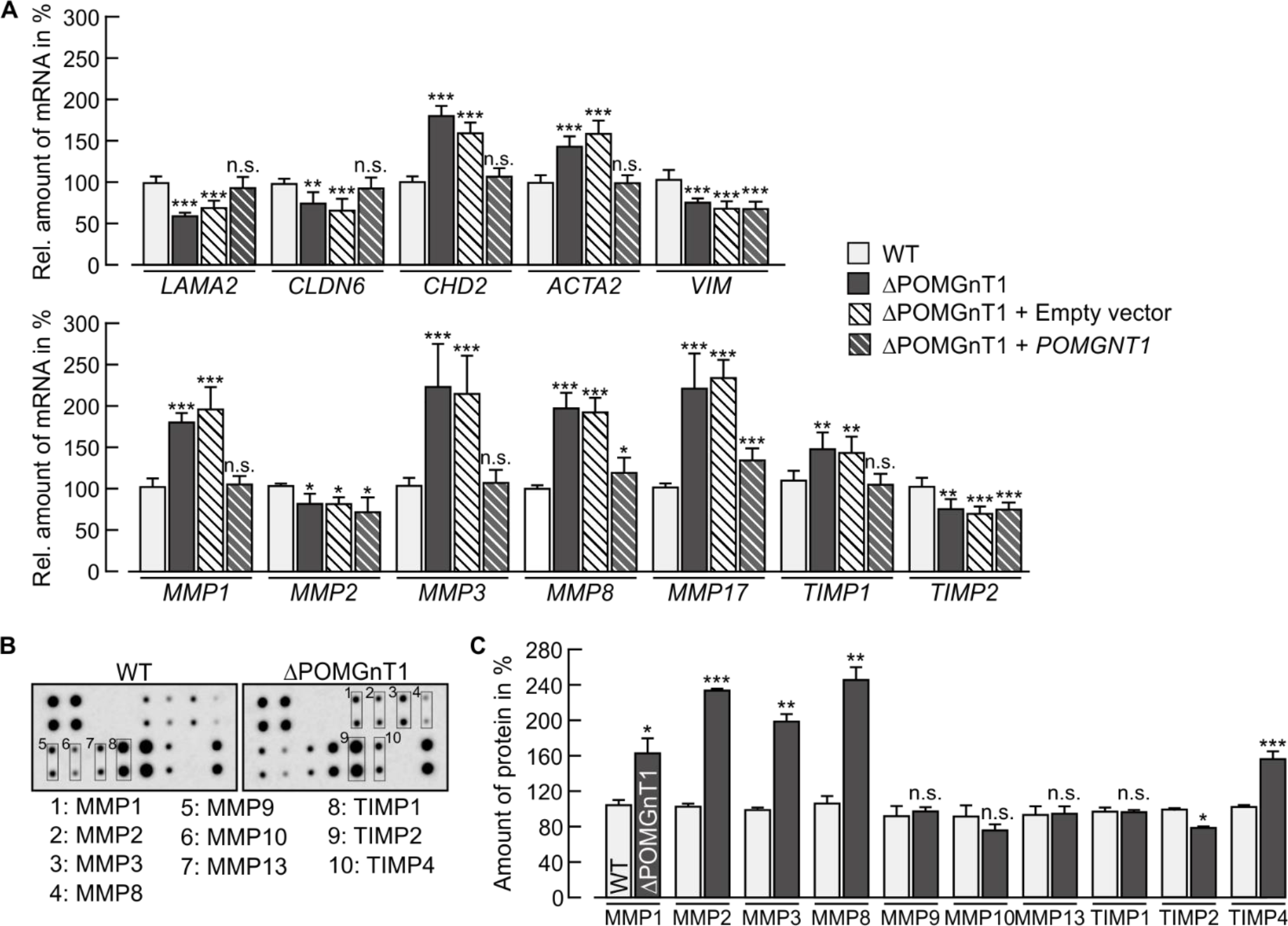
*POMGNT1* deficiency induces changes in the expression of EMT-related genes in HEK293T cells. (A) Gene expression analysis by qRT-PCR of selected EMT-related genes. Epithelial markers: *LAMA2* and *CLDN6;* mesenchymal markers: *ACTA2,CDH2* and *VIM;* matrix metallopeptidases (MMPs): *MMP1, MMP2, MMP3, MMP8* and *MMP17;* tissue inhibitors of metalloproteinases (TIMPs): *TIMP1* and *TIMP2*. Relative mRNA levels in WT cells (white bars) were compared with levels detected in ΔPOMGnT1 cells (gray bars), *POMGNT1*-transfected ΔPOMGnT1 cells (ΔPOMGnT1 + *POMGNT1;* strip gray bars) and empty vector-transfected ΔPOMGnT1 cells (ΔPOMGnT1 + empty vector; strip white bars). Transcript levels are calculated as %, considering transcript levels of respective genes in WT cells as 100%. HPRT was used for normalization. Assays were performed in duplicate with three dilutions from three biological replicates. (B) Representative MMP array performed with 500 μg of whole cell extract from WT cells (left panel) and ΔPOMGnT1 cells (right panel). Antibodies against MMPs and TIMPs are spotted in duplicate. Signals for MMPs and TIMPs on the representative membranes are highlighted in boxes and denoted with numbers. (C) Densitometry-based quantification of MMP and TIMP signals detected in MMP array for WT and ΔPOMGnT1 cells. Protein signals detected in ΔPOMGnT1 cells were normalized to the reference protein signal and subsequently normalized to the protein/reference signal ratio in WT cells. Protein levels of MMPs and TIMPs in ΔPOMGnT1 cells are calculated in %, considering pixel density of the respective protein in WT cells as 100%. Analysis is based on two independent experiments. (A, C) Data are presented as means ± SD. Asterisks denote statistical significance in comparison to WT cells: * *p* ≤ 0.05, ** *p* ≤ 0.01, *** *p* ≤ 0.001, *n.s*. not significant.

EMT processes are associated with the upregulation of matrix metallopeptidases (MMPs) which represent a family of zinc-dependent endoproteases involved in degrading ECM components (34). A strong and specific increase of *MMP1, MMP3, MMP8* and *MMP17* mRNA is observed in ΔPOMGnT1 cells (**Fig. 8A**). Also expression of metallopeptidase inhibitors (TIMPs; (35)) is affected. Likewise, analysis of the abundance of MMP proteins using a human MMP antibody array showed that MMP1, MMP3 and MMP8 levels are increased (**Fig. 8, B** and **C**).

To determine whether decreased POMGNT1 activity also triggers an EMT-like transcriptional response in MEB patient-derived fibroblasts, mRNA levels of the above genes were assessed. Indeed, in comparison to control fibroblasts, epithelial markers (*LAMA2* and *CLDN6*) were found to be significantly reduced, whereas mesenchymal markers (*ACTA2*, *VIM* encoding vimentin and *FN* encoding fibronectin) were significantly elevated in MEB patients (**Fig. 9A**). In addition, the transcriptional modulation of *MMPs* and *TIMPs* was also comparable to the changes observed in ΔPOMGnT1 HEK293T cells with particularly large increase in transcript levels of *MMP1* and *MMP17* (**Fig. 9B**).

**Figure 9.**
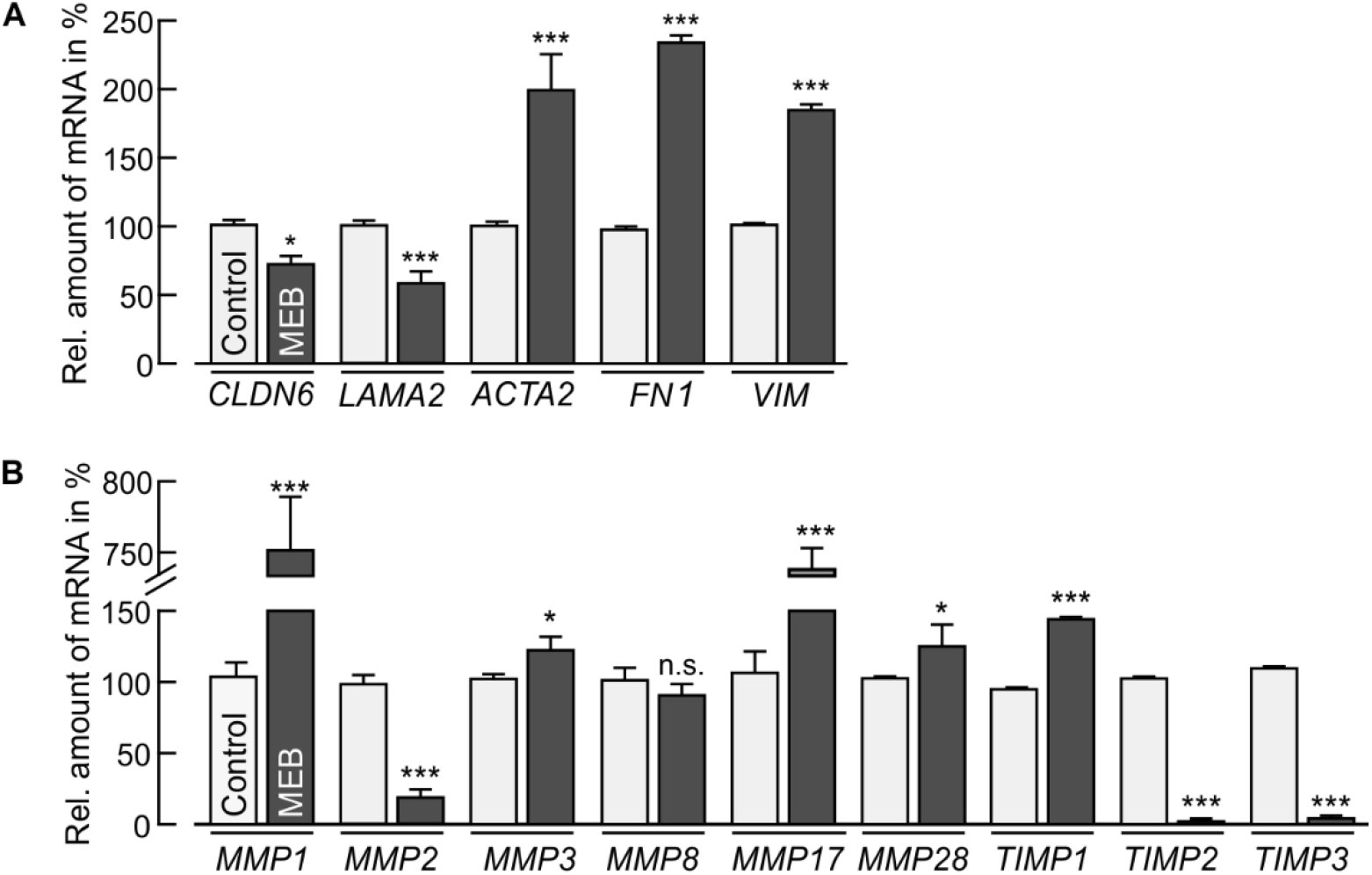
MEB patient-derived fibroblasts show changes in the expression of EMT-related genes. Gene expression analysis by qRT-PCR of (A) epithelial (*LAMA2* and *CLDN6*), and mesenchymal (*ACTA2*, *FN1* and *VIM*) marker genes as well as (B) the indicated *MMPs* and *TIMPs*. Data were calculated as described in Fig. 8A.

Altogether, our data demonstrate that lack of POMGNT1 activity induces an EMT-like transcriptional response resulting, *inter alia*, in elevated levels of ECM degrading enzymes.

## Discussion

α-Dystroglycanopathy is associated with the development of a variety of medical conditions, including muscular dystrophy and mild to severe changes in the central nervous system and the eyes. So far, 18 causative genes for α-dystroglycanopathy have been identified that are associated with the *O*-mannose glycosylation pathway of α-DG (10). Among those are *POMT1/POMT2* and *POMGNT1*, the major disease causing factors of the Walker Warburg syndrome (WWS; lacking classic *O*-mannose glycans), and MEB (no elongation of *O*-linked mannoses), respectively. Current knowledge regarding the predominant disease mechanism suggests that incomplete matriglycan biosynthesis impairs the binding of α-DG to extracellular matrix proteins such as laminin, resulting in damage to cell membrane integration and defective basement membranes (9). In recent years, it turned out that changes in classic *O*-mannosylation due to altered expression of POMT2 impact also on epithelial cadherin (E-Cdh)-mediated cell-cell adhesion during murine embryonic development and in human gastric carcinoma (36, 37). However, whether the observed changes only apply to E-Cdh and POMT2 is unclear, as are the underlying molecular mechanisms. Taking advantage of POMGNT1-deficient HEK293T cells and MEB patient-derived fibroblasts, we now show that, in general, aberrant classic *O*-mannosylation impacts on cadherin-mediated cell-cell adhesion. We demonstrate that loss of POMGNT1 function results in (1) reinforced N-Cdh-mediated cell-cell adhesion and impaired cell migration potential; (2) increased levels of N-Cdh and changed *N*-glycan structures on its extracellular domain, which in turn enhance its intrinsic adhesive properties; (3) activation of ERK1/2 and p38 signaling pathways and induction of transcriptional modulations which are comparable to EMT-like events.

Changing cell migration behavior is a (patho)physiological process determined by the opposing forces that define cell-ECM and cell-cell interactions. In invasive cancers, decreased levels of POMT1, POMT2 and other enzymes of the classic *O*-mannosylation pathway correlate with high cell migration and invasion (28). But, low POMGNT1 levels can also hinder cell migration. A *POMGNT1*-based MEB mouse model revealed clusters of granule cells within the cerebellum, which have failed to migrate during development (14). Further, Abbott and coworkers reported that knock-down of *POMGNT1* and *MGAT5B* impairs neuronal cell migration (38), and Lan and coworkers found that silencing of *POMGNT1* decreases cell proliferation and invasion in glioblastoma (16). Therefore, *POMGNT1* deletion may also act as a break on invasion and migration, however, the direct molecular mechanism for the observed phenotypes are not fully understood. Our HEK293T cell model recapitulates the observed impact of *POMGNT1* on cell adhesion and migration observed in neuronal and glioblastoma cells, and identifies increased N-Cdh-mediated cell-cell adhesion as one of the reasons that influences the cellular migration potential when POMGNT1 activity is reduced. In addition to the increased amount of N-Cdh, its *N*-glycosylation pattern is changed to further promote N-Cdh homotypic interactions. In general, on N-Cdh from ΔPOMGnT1 *N*-glycans with a lower degree of galactosylation and sialylation, along with a slight increase in the abundance of bisecting GlcNAc are observed. These changes are driven by the transcriptional regulation of specific glycosyltransferases, such as B4GALT1 and ST6GAL1, which act as heteromeric complex in the successive addition of terminal β1,4-linked galactose and α-2,6-linked sialic acid to *N*-glycans (39). Most interestingly, the changes in *N*-glycosylation are site specific and especially site N402 in the EC3 domain contains less branched complex glycans, which have been reported to favor cell-cell adhesion. N402, along with N273 and N325, has previously been identified as one of the most relevant *N*-glycosylation sites that strongly increase cis-dimerization of N-Cdh when complexity of *N*-glycans is decreased (21). In line with previous findings of the group of S. Pinho (in collaboration with our group) that in human gastric carcinomas expression levels of POMT2 correlate with altered structures of *N*-glycans on E-cadherin (37), we now demonstrate a link between classic *O*-mannosylation and *N*-glycosylation of cadherins which was for the first time analyzed in such depth.

Very recently it turned out that cadherins are also major targets of TMTC1-3 which add single non-extended *O*-linked mannoses to EC domains (18) that impact on cellular adherence (40). Indirect effects on TMTC-based *O*-mannosylation of N-Cdh due to lack of *POMGNT1* could be excluded, as its *O*-mannosylation is not altered in our HEK293T cell model.

Increased expression of N-Cdh is part of a transformation process that mostly epithelial cells undergo known as epithelial-to-mesenchymal transition (30). EMT is inherent to physiological processes such as embryonic stem cell differentiation and development (41) and has further been associated with pathological conditions such as wound healing, fibrosis and cancer stemness and progression (42–45). Key events of EMT are dissolution of epithelial cell-cell junctions, loss of the apical-basal polarity and reorganization of the cytoskeleton as well as downregulation of epithelial gene expression in favor of genes that establish the more motile mesenchymal phenotype (33). Long regarded as an all-or-nothing event, EMT is now considered a transition suggesting a gradual and reversible process that does not exclusively concern epithelial cells (46). EMT does not necessarily include increased cell motility but can even lead to the opposite migration behavior (47, 48). Our results on *POMGNT1*-deficient HEK293T and fibroblast cells point to a partial EMT event that is accompanied by reduced expression of epithelial and increased expression for most of the mesenchymal markers tested. Very recently differential expression of POMGNT1 in human glioma cell lines has been reported to impact on the expression of some EMT marker proteins (17) further corroborating the general relevance and validity of our HEK293T cell model. We also observe a predominant induction of MMPs that is indicative of an EMT-like transition. Increased level of MMPs have been shown to be involved in degradation of the basement membrane in developmental processes (49, 50), wound healing (51) and cancer progression (34). Since breakdown of basement membranes is also one of the hallmark events in MEB disease (11) or α-dystroglycanopathy in general, elevated levels of MMPs could increase ECM protein degradation and impede cell-ECM interactions further deteriorating the clinical outcome of these patients. A similar role has been suggested for MMP2 and MMP9 in physiological and pathological conditions involving members of the dystrophin glycoprotein complex (52).

In ΔPOMGnT1 cells partial EMT correlates with sustained activation of ERK1/2 and p38 MAPK signaling which have been associated with EMT induction (53). Whereas these pathways seem to be decisive for the EMT response observed, the source of its activation remains unclear. Although other factors cannot be excluded completely, the most likely candidate that induces ERK signaling is the dystrophin glycoprotein complex and DG in particular, that was suggested as a multifunctional adaptor, capable of interacting with components of the MAPK cascade including MEK and ERK (27). The role of DG on developmental and pathological EMT processes is well documented (50, 54). In line with our data, α-DG glycosylation and its interaction with laminin has been suggested to enable β-DG to sequester ERK preventing it from translocating into the nucleus and promoting differentiation of a mesenchymal phenotype (55). To our best knowledge, so far no link between p38 and DG has been reported. How p38 MAPK pathway is linked to POMGNT1 deficiency will be an interesting future question. In many cancers, heterogeneity in protein structures due to aberrant glycosylation results in induction of EMT (31). Thus, the observed modulation of *N*-glycan modifying enzymes in ΔPOMGnT1 cells also might contribute to some extend to the induction of EMT-linked signaling pathways.

Our study proved ΔPOMGnT1 HEK293T cells as an excellent system to study molecular details underlying α-dystroglycanopathy. Our data suggest a model (**Fig. 10**) in which due to the loss of *O*-linked matriglycan on α-DG, ERK and p38 signaling cascades are activated in POMGNT1-deficient cells. As a consequence, EMT-like transcriptional events result *inter alia* in the induction of N-Cdh (*CDH2*) and MMPs. In addition, transcriptional modulation of *N*-glycan modifying enzymes contributes to increased N-Cdh homotypic interactions. In combination, these events result in enhanced cell-cell adhesion and disordered basement membranes thereby contributing to the molecular pathogenesis of MEB disease.

**Figure 10.**
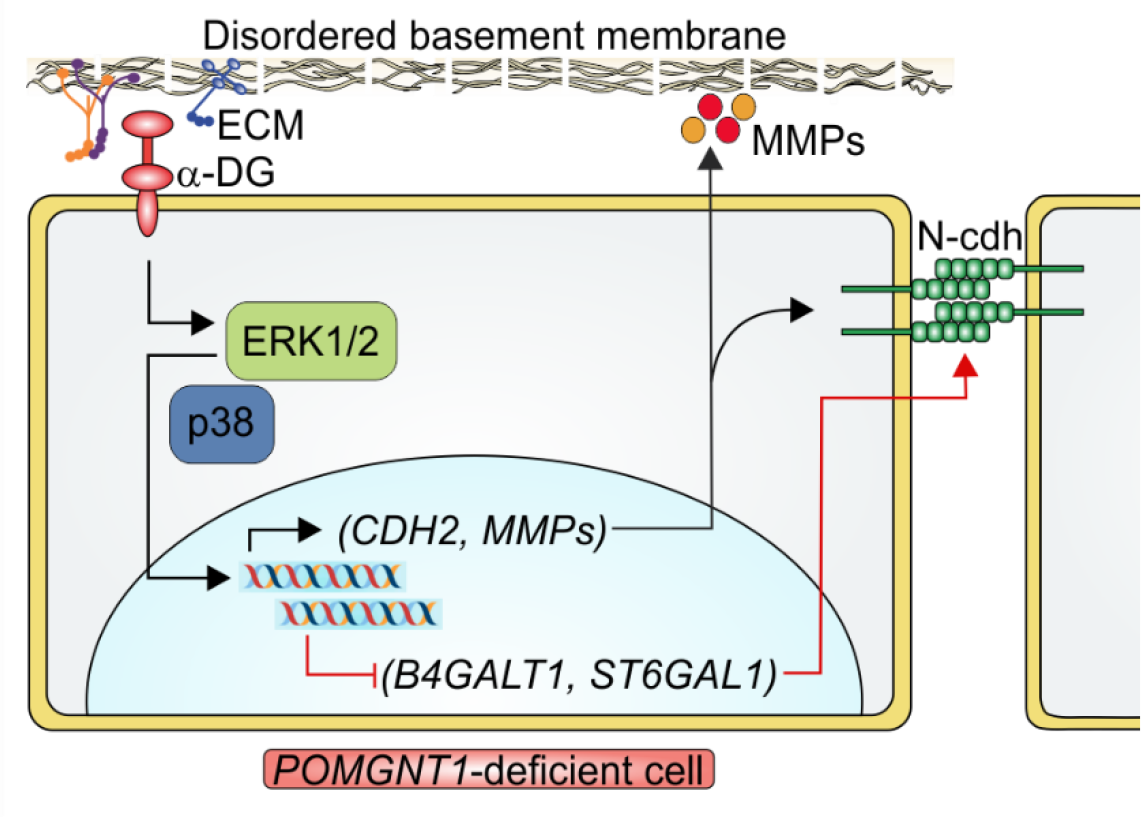
Hypothetical model integrating the observed changes in POMGNT1-deficient cells. For details see discussion

## Experimental procedures

### Cell lines and culture conditions

HEK293T cells were maintained in high glucose DMEM (ThermoFisher Scientific) supplemented with 10% FBS (ThermoFisher Scientific), 1% non-essential amino acids (ThermoFisher Scientific) and 1% Pen/Strep (ThermoFisher Scientific) (referred to as complete media) at 37 °C in a humidified incubator with 5% of CO_2_.

### Generation of TALEN-mediated POMGNT1 knock-out cells

Transcription Activator-Like Effector Nuclease (TALEN)-based constructs were designed utilizing the TAL Effector Nucleotide Targeter 2.0 online tool (56). Three TALEN constructs were selected which target the ATG start codon of *POMGNT1*, the coding sequence of the D-X-D motif and an intron-exon boundary upstream of the exon containing the D-X-D motif. All constructs present a restriction site close to the predicted TALEN cleavage site to allow screening of induced mutations by restriction fragment length polymorphism (RFLP) analysis. TALENs were assembled using the Golden Gate TALEN and TAL Effector Kit 2.0 (57) and employing pC-GoldyTALEN as the final expression vector (#1000000024 and #38143 respectively; both from Addgene). Correct assembly of TALEN plasmid DNA was verified by restriction site analysis and sequencing and TALEN plasmids were transfected into wild type HEK293T cells. Next, genomic DNA was isolated from transfected cells using the DNeasy Blood and Tissue Kit (Qiagen) according to manufacturer’s instructions and *POMGNT1* regions were amplified by PCR and digested using restriction sites close to the predicted TALEN cutting site. Detection of PCR products resistant to restriction, indicated efficient cleavage by TALENs and was followed by dilutional cloning (1 cell/100 μl/96-well) and further genomic DNA isolation, PCR and RFLP analysis to obtain single cell knock-outs. From the selected TALEN construct combinations, the one targeting the ATG start codon of *POMGNT1* (close to a *HpaII* restriction site; forward TALEN construct: TGGTGACCCGCCAAT, reverse TALEN construct: GGAAGCCCAGCCCCCTC) was the most efficient in inducing mutations and was therefore used for subsequent dilutional cloning, PCR and RFLP. PCR analysis and sequencing of genomic DNA confirmed the knock-out of *POMGNT1* (**Fig. S1**). *POMGNT1* knock-out clones were furthermore evaluated by Wheat Germ Agglutinin (WGA) enrichment for absence of reactivity towards matriglycan on α-DG recognized by IIH6 antibodies (58) (see below). For *POMGNT1* complementation, pMBA40 was stably transfected into *POMGNT1* knock-out HEK293T cells.

### Generation of stable cell lines

Transfections were performed with the calcium phosphate method (59). In short, 5 μg of plasmid DNA in 250 mM CaCl_2_ were added dropwise to double strength HBSS (280 mM NaCl, 2.8 mM Na_2_HPO_4_, 50 mM HEPES, pH 7.2) while mildly vortexing and the final transfection solution was applied drop wise to cells with 40-50% confluency in 15 cm dishes. After 6 h the medium was replaced, and selection was performed with 500 μg/ml Zeocin (Invitrogen) for 4 weeks.

### Patient material

The study was performed in accordance with the declaration of Helsinki and approved by the Ethics Committee of the Medical Faculty Heidelberg. MEB skin fibroblasts were derived from a patient who presented with characteristic stigmata as mental retardation, blindness, spastic tetraplegia, contracture of limbs and severely reduced proprioceptive reflex due to variant c.535_751del (p.Asp179Argfs*11) in the *POMGNT1* gene (NM_017739.4). Control skin fibroblasts were obtained from two healthy anonymous donors. Fibroblasts were maintained in high glucose DMEM (ThermoFisher Scientific) supplemented with 10% FBS (PAN Biotech) and 1% Pen/Strep at 37 °C in a humidified incubator with 5% of CO_2_. Growth medium was changed every 72 h.

### Plasmid construction

To generate pMLHD7 for expression and subsequent purification of N-Cdh extracellular domain (EC) (EC-N-Cdh) C-terminally fused to a His_8_/Strep-tag, PCR was performed on mouse cDNA using primers Forward: ac*GGCGCGCC***GACTGGGTCATCCCGCCAA** and Reverse: *ccgCTCGAG*<u>TTATTTTTCGAACTGCGGGTGGCTCCA</u>AGCGCTGTGATGATG<u>GTGGTGAGGTGATG</u>**GGCGCCCGTGCCAAGCC**. The encoded protein shows 96.8% identity with human N-Cdh. *AscI* and *XhoI* restriction sites are shown in italics, His_8_- and Strep-tag coding sequences are underlined and the overlapping sequences to the extracellular domain of N-Cdh are depicted in bold. Purified PCR products were digested with *AscI* and *XhoI* and inserted into the pSecTag2 expression vector (Invitrogen).

Human full-length *POMGNT1* was amplified by PCR from cDNA extracted from HEK293T cells using primers Forward: cga*CTCGAG***TGTCTGTTCTGGGGCTCCTGG** and Reverse: cttga*GCTAGC***CCCGCCAATCCGGTATGGA**. *XhoI* and *NheI* restriction sites are shown in italics and the overlapping sequences to human *POMGNT1* are depicted in bold. Purified PCR products were digested with *XhoI* and *NheI* and inserted into the pcDNA6/V5-His expression vector (Invitrogen) generating plasmid pMBA40 for *POMGNT1* complementation. All plasmids created in this study were verified by sequencing.

### Whole cell lysate and crude membrane preparation

Cells at 90% confluency (~5.0 x 10^6^ cells) were washed with ice-cold PBS and lysed with 500 μl of lysis buffer (10 mM Tris-HCl pH 7.6, 1% (w/v) SDS, 1 mM EDTA, supplemented with Halt™ protease and phosphatase inhibitor cocktail without EDTA (ThermoFisher Scientific)). Lysis was performed for 30 min at 4 °C on a rotator with occasional vortexing and whole cell lysates were collected upon centrifugation (16,000 x g, 10 min, 4 °C).

For crude membrane extraction, cell pellets of ~5.0 x 10^6^ cells were resuspended in 750 μl of ice-cold hypotonic buffer (20 mM Tris-HCl pH 7.4, 10 mM NaCl, 1.5 mM MgCl_2_, supplemented with protease inhibitor cocktail without EDTA (ThermoFisher Scientific) and 0.05 U/μl benzonase (Sigma Aldrich) and homogenized by passing through a 27-gauge needle for 10 times. Homogenized samples were subjected to centrifugation (16,000 x g, 15 min, 4 °C) and resulting pellets were resuspended in 500 μl of hypotonic buffer. Protein concentration was measured using the BCA protein assay (Pierce).

### WGA pull-down

Around 1.0 x 10^7^ cells were incubated with 500 μl of ice-cold homogenization buffer (50 mM Tris-HCl pH 7.6, 150 mM NaCl, 1% Triton X-100, 1 mM PMSF and 1 mM benzamidine) for 20 min at 4 °C. Lysates were subjected to centrifugation (20,000 x g, 5 min, 4 °C), equivalent amounts of solubilized proteins from the supernatant (input) were added to 75 μl of WGA-bound agarose beads (Vector Laboratories) and incubated o/n at 4 °C. After centrifugation (2,000 rpm, 3 min, 4 °C) the supernatants (void) were removed, and beads washed three times with 1 ml of ice-cold wash buffer (50 mM Tris-HCl pH 7.6, 150 mM NaCl, 0.1% Triton X-100). Bead-bound proteins were eluted by boiling in 100 μl of 2.5x SDS sample buffer (5x sample buffer: 250 mM Tris-HCl pH 6.8, 500 mM DTT, 50% glycerol (v/v), 0.25% Bromophenol blue (w/v), 10% SDS (w/v)) (3 min, 95 °C) followed by centrifugation at 14,000 rpm for 1 min. 30 μl of eluted proteins were analyzed by SDS-PAGE and Western Blot.

### Heterologous protein expression and purification

For expression of EC-N-Cdh, HEK293T cells stably transfected with pMLHD7 were cultured for 72 h in serum-free medium. Upon initial centrifugation for removal of dead cells (1,000 x g, 10 min, 4 °C) culture medium from two 90% confluent 15 cm dishes was filtered through a 1.2 μm filter (Sartorius Stedim Biotech) and applied to a HisTrap column (1 ml bed volume, GE Healthcare) at a drop rate of 500 μl/min at 4 °C for o/n. The column was washed with 10 ml of wash buffer (20 mM Na3PO4, 500 mM NaCl, 5 mM imidazole, pH 7.4) and elution of bound proteins was performed in 10 fractions of 500 μl elution buffer (20 mM Na3PO4, 500 mM NaCl, 500 mM imidazole, pH 7.4). Protein concentration of eluates was determined by BCA protein assay (Pierce) and analyzed by SDS-PAGE followed by Coomassie blue staining and Western Blot.

### Western Blot

Protein samples were resolved on 10% SDS PAA gels and transferred on nitrocellulose membranes according to standard protocols. After blocking and incubation with respective antibodies, proteins were detected using ECL Prime Western Blotting Detection Reagent (GE Healthcare) and imager ImageQuant LAS 500 (GE Healthcare). Protein levels were normalized to the loading control using ImageJ software (version 1.52i, NIH). Antibodies used: Rabbit anti-N-cadherin (#22018-1-AP, Proteintech), 1:1000; Mouse anti-α-dystroglycan, clone IIH6C4 (#05-298, Millipore), 1:2500; Rabbit anti-β-dystroglycan (AP83, kind gift from K. Campbell), 1:1000; Mouse anti-Sec61α (#sc-393182, Santa Cruz), 1:100; Mouse anti-Penta-His-tag (#P-21315, ThermoFisher Scientific), 1:2000; Mouse anti-β-tubulin (#T6199, Sigma Aldrich), 1:500; Rabbit anti-ERK1/2 (#9102, Cell Signaling), 1:1000; Rabbit anti-Phospho-ERK1/2(Thr202/Tyr204) (#9101, Cell Signaling), 1:1000; Rabbit anti-MEK1/2 (#9122, Cell Signaling), 1:1000; Rabbit anti-Phospho-MEK1/2(Ser217/221) (#9121, Cell Signaling), 1:1000; Goat anti-rabbit HRP-conjugated IgG (#12-348, Sigma Aldrich), 1:5000; Rabbit anti-mouse HRP-conjugated IgG (#AP160P, Sigma Aldrich), 1:5000.

### Cell proliferation and migration assay

Cell proliferation and migration were monitored by the xCELLigence system (ACEA Biosciences) according to manufacturer’s instructions. The xCELLigence system is a real-time, non-labeled, impedance-based cell analysis system that allows cell proliferation and migration to be monitored in a continuous and quantitative manner.

To monitor proliferation, cells from exponential phase cultures were detached using 0.05% Trypsin-EDTA (Invitrogen), counted with a Scepter 2.0 device (Merck-Millipore) and 2.0 x 10^4^ cells were seeded in each well of a 96-well E-Plate in a final volume of 200 μl complete growth medium. The xCELLigence station was placed in a standard cell culture incubator and the cell index was monitored every 15 min over a period of 90 h. The background impedance of the E-Plate was determined with complete growth medium alone.

Cell migration was monitored using a 16-well cell invasion/migration plate with each well consisting of an upper and a lower chamber separated by a microporous membrane containing randomly distributed 8 μm pores. Cells deprived of FBS for 24 h were seeded in the upper chambers at a density of 7.5 x 10^3^ cells/well in a total volume of 180 μl of FBS-free media. The lower chamber was filled with 160 μl of complete medium. After mounting both chambers to each other, the plate was loaded into the xCELLigence station and placed into a standard cell culture incubator. Impedance read out, expressed as cell index, was executed every 15 min over a period of 90 h. Cell proliferative and migratory responses were determined by calculating the mean cell index value and slope of the line between two given time points. Each sub-population was measured in quadruplicate.

### Cell-ECM and cell-cell adhesion assay

ECM adhesion was monitored using the Millicoat™ ECM Screening Kit (Millipore). Prior to each experiment, cells were deprived of FBS for 24 h. To initiate the experiment, sub-confluent cell monolayers were treated with 10 mM EDTA in PBS to obtain a single cell suspension. Cells were washed twice with FBS-free medium, added at a density of 8.0 x 10^4^ cells per 96-well in a total volume of 180 μl and allowed to adhere for 1 h at 37 °C. Unattached cells were removed by washing three times with ice-cold PBS containing 1 mM CaCl_2_ and 1 mM MgCl_2_. Attached cells were fixed with freezer-cold 100% methanol. Adherent cells were stained with 0.2% (v/v) crystal violet, washed and the staining was released according to manufacturer’s instructions. To determine the relative number of bound cells, OD600 recorded for BSA-coated wells was subtracted from OD600 recorded for ECM-coated wells. Percentage of adhesion of ΔPOMGnT1 cells to ECM was calculated taking adhesion of wild type as 100%.

To analyze cell-cell adhesion, 1.0 x 10^4^ cells/well were plated in flat-bottom 96-well plates and allowed to grow to a confluent monolayer for 48 h. Single cell suspensions (5.0 x 10^4^ cells/well) were added on top of each cell monolayer and allowed to adhere for 20 min at 37 °C. Subsequent steps including washing, fixing, staining and OD600 measurement were performed as mentioned above.

To analyze N-Cdh-mediated cell-cell adhesion, wild type or ΔPOMGnT1 cells deprived of FBS o/n were incubated for 30 min with 15 μg/ml human anti-N-Cdh antibody (#22018-1-AP, Proteintech) and seeded on anti-N-Cdh antibody pre-treated confluent cell monolayers. As a negative control, both single cell suspensions and cell monolayers were pre-treated with anti-human IgG antibody. The adhesion assay was performed as described above. The absorbance measured was normalized against the mean absorbance of wells, where no cells were added. The quantification of respective cell-cell adhesion was represented as relative adhesion in %, considering relative adhesion of wild type to wild type cells pre-treated with anti-human IgG antibody as 100%.

### Bead aggregation assay

To detect the homophilic interactions of the recombinant extracellular domain (EC) of N-Cdh, bead aggregation assays were conducted as previously described with slight modifications (26). Briefly, EC-N-Cdh protein purified from wild type and ΔPOMGnT1 HEK293T cells (see above) was incubated with pre-washed Dynabeads™ (Invitrogen) at a ratio of 40 μg of protein per 40 μl of bead suspension o/n at 4 °C. Next, beads were washed twice in binding buffer (50 mM Tris-HCl pH 7.2, 100 mM NaCl, 10 mM KCl and 0.2% BSA) and resuspended in 200 μl of the same buffer. Bead suspensions were briefly sonicated and split into two tubes, and either 2 mM CaCl_2_ or 2 mM EDTA was added to each tube for the “calcium” and “no-calcium” conditions, respectively. The samples were incubated at room temperature. Bead aggregation was assessed by spotting 10 μl of each condition on glass depression slides at various time points and images were captured on an Axiostar microscope (Zeiss) at 10 fold magnification. Bead aggregates were quantified using ImageJ software (version 1.52i, NIH). Briefly, images were thresholded, the area of the detected aggregated particles was measured in units of pixels and the average size was calculated. Assays were repeated three times using two independent protein purifications and their mean aggregate size (± SDM) at each time point was plotted. To quantify protein coupled to the beads, 100 μl of bead suspension were heated with SDS sample buffer (10 min, 95 °C) and the supernatant was analyzed by SDS-PAGE and Western Blot.

### Proteomics

Proteome analysis of wild type and ΔPOMGnT1 cells was performed in five replicates. Cells were grown to 90% confluency in 15 cm dishes and lysed in 1 ml lysis buffer (PBS with 1% Triton X-100 (v/v), 1 mM PMSF, 1 mM benzamidine, 0.25 mM TLCK, 50 μg/ml TPCK, 20 μg/ml antipain, 1 μg/ml leupeptin, 1 μg/ml pepstatin) for 30 min on ice. Supernatants of whole cell lysates were collected after centrifugation (20,000 x g, 10 min, 4 °C), sonicated (Branson Sonifier 450 at 60% Duty Cycle, Level 6 Output Control) for 15 min and resolved on 10% SDS PAA gels. Gels were washed four times in ddH2O (200 ml, 30 min), stained in a colloidal Coomassie solution (5% aluminium sulphate hydrate (w/v), 10% ethanol (v/v), 0.02% Coomassie Brilliant Blue G-250 (w/v), 2% phosphoric acid (v/v)) for 1-2 h and destained o/n in 500 ml ddH2O. Proteins were isolated from respective gel pieces, carbamidomethylated, subjected to tryptic digestion (1:50) and acidified with 1% trifluoroacetic acid. Peptide analysis was performed on a liquid chromatography-mass spectrometer (LC-MS), therefore an UltiMate 3000 nanoRSLC (ThermoFisher Scientific) was used coupled to an LTQ Orbitrap Elite MS (ThermoFisher Scientific). Peptides were trapped and desalted on a C18-column (5 μm Acclaim PepMap100 300 μm x 5 mm, ThermoFisher Scientific) at a flow rate of 30 μl/min with solution A (1% acetonitrile (ACN), 1% formic acid (FA)). After 3 min of loading, peptides were separated on an analytical column (2 μm Acclaim PepMap RSLC 75 μm x 25 cm, ThermoFisher Scientific) using a 150 min method and an effective gradient of solution A (1% ACN, 5% DMSO, 0.1% FA) and B (90% ACN, 5% DMSO, 0.1% FA): 3% to 40% B in 110 min; 40% to 90% B in 10 min; flow rate: 300 nl/min. Full-scan mass spectra were acquired in the Orbitrap analyzer in the positive ion mode at 60,000 resolution at 200 m/z. A lock-mass of 445.120020 m/z was used for internal recalibration, followed by collision-induced dissociation fragmentation of the thirty most intense ions (top 30). Spectra were acquired in the centroid mode using an ion trap detector. MS data were analyzed by MaxQuant (version 1.6.5.0) (60) and Andromeda search engine (61) with standard settings except noted otherwise. The UniProt human database was used (downloaded on July 08/2019, reviewed database entries: 26468). Carbamidomethylation of cysteine was set as a fixed modification. Methionine oxidation, asparagine/glutamine deamidation, protein N-terminus acetylation and serine/threonine mono hexosylation were set as variable modifications. A minimal peptide length of seven amino acids, and at maximum two miscleavages (trypsin/p) were allowed. The maximum protein, peptide and site false discovery rates were set to 0.05. “Re-quantify” and “Match between runs (2 min)” was activated. LFQ-intensities of the proteins were used and analyzed in the program Perseus (label-free quantification). Only identified by site, reverse and potential contaminants were filtered out. LFQ-intensities were logarithmized to the base of 2. Multi scatter plot and histograms were performed with that data. For the analysis of the principle component analysis (PCA) and the heat map the candidates were filtered, only proteins with valid values in all ten samples were allowed. The MS proteomics data have been deposited to the ProteomeXchange Consortium via the PRIDE (62) partner repository with the dataset identifier PXD021321.

### N-and O-mannose glycoproteomics of N-cadherin

Sample preparation, measurement and glycoproteomic analysis was conducted as previously described in Hoffmann et al. (63). Briefly, EC-N-Cdh purified from wild type and ΔPOMGnT1 HEK293T cells was proteolytically digested for 8 h at 37 °C using trypsin (enzyme:substrate ratio 1:30) as well as proteinase K (enzyme:substrate ratio 1:10). Specifically, for *O*-mannose glycoproteomics, proteins were treated with PNGase F (Sigma Aldrich) (0.5 U, 37 °C, 8 h) prior to the proteolytic digest. Enrichment of *N*-linked and *O*-mannosyl glycopeptides was performed using spin-cotton HILIC SPE. Enriched glycopeptides were analyzed by reverse-phase liquid chromatography coupled online to RP-LC-ESI-OT MS/MS (ThermoFisher Scientific) using two higher-energy collision dissociation fragmentation regimes (HCD.low and HCD.step). Glycopeptide mass spectra were analyzed using glyXtoolMS (in-house developed software) (64), as well as Byonic and Byologic (both from Protein Metrics). All MS glycoproteomics raw data have been deposited to the MassIVE repository under the dataset identifier MSV000085243.

### N-glycome analysis of N-cadherin

*N*-glycan structures were analyzed by xCGE-LIF, as previously described with slight modifications (65–67). Briefly, proteins were linearized and disulfide bonds were reduced using SDS and DTT for 10 min at 60 °C. *N*-glycans were released from protein backbone by PNGase F incubation (Sigma Aldrich) for 12 h at 37 °C. Released *N*-glycans were labeled with 8-aminopyrene-1,3,6-trisulfonic acid (APTS, Sigma Aldrich), followed by a sample clean-up step (HILIC SPE) using glyXbeads™ (glyXera) to remove excess label, salt and other impurities. The purified labeled *N*-glycans were analyzed by xCGE-LIF. Using the glycoanalysis system glyXbox™ (incl. kits, software (glyXtool™ v5.3.1) and database (glyXbase™); glyXera), the generated *N*-glycan electropherogram data (migration times) were normalized to an internal standard, resulting in a so-called “*N*-glycan fingerprint” with highly reproducible standardized migration time units. Subsequently, glyXtool™ was utilized for automated peak picking, integration and relative quantification, as well as for the structural assignment of the *N*-glycan peaks by migration time matching to the *N*-glycan database glyXbase™. To confirm *N*-glycan compositions and linkages, a comprehensive exoglycosidase sequencing was performed, using the enzymes α(2–3) sialidase (Prozyme), α(2–3,6,8) sialidase (Prozyme), α(1–2,3,4,6) fucosidase (Prozyme), β(1–4,6) galactosidase (Prozyme) and α(1–2,3,6) mannosidase (Prozyme). All exoglycosidase digests were carried out at 37 °C and at the reaction conditions recommended by the respective enzyme supplier. Each exoglycosidase enzyme was carefully tested for activity, specificity and possible side activities. Please note, differing from the original protocol, HEK293T cell-derived EC-N-Cdh samples were desalted prior to *N*-glycan release. To this end, each N-Cdh sample was desalted against PBS using Amicon centrifugal filters with a 3 kDa molecular weight cutoff according to the manufacturer’s protocol (Merck-Millipore). Unprocessed xCGE-LIG *N*-glycocomics raw data are available upon request.

### Quantitative real-time PCR

Total RNA from cells was extracted using the Universal RNA Purification Kit (Roboklon) according to the manufacturer’s recommendations. 2 μg of total RNA were reverse transcribed into cDNA using the FastGeneScriptase Basic cDNA Kit (Tiangen Biotech) with random Oligo(dT) primers following manufacturer’s instructions. Quantitative real-time PCR was performed on a Rotor-Gene Q (Qiagen) using the qPCRBIO SyGreen Mix Lo-ROX (PCR Biosystems). PCR reactions were performed as follows: 95 °C for 30 s followed by 40 cycles of 95 °C for 30 s, 60 °C for 30 s and 72 °C for 30 s in a final volume of 15 μl containing 3 μl of 1:10 cDNA dilution and 0.5 mM of respective oligonucleotides. As technical replicates and for determination of qRT-PCR efficiency, 1:100 and 1:1000 cDNA dilutions were included. Only qRT-PCR reactions with efficiencies ranging from 0.9 to 1.1 were further analyzed. Gene expression was normalized to expression of the housekeeping gene hypoxanthine-guanine phosphoribosyltransferase (HPRT). For calculation of relative gene expression, the standard curve-based method was used. The primers are provided in **Table S1.**

### nCounter gene expression profiling

Total RNA was isolated from 1.0 x 10^6^ HEK293T cells by using the RNeasy Mini Kit (Qiagen) in combination with the QIAshredder system (Qiagen) according to the manufacturer’s protocol. 50 ng of total RNA were used per hybridization reaction to determine the transcript levels of Golgi glycosyltransferases.

Further processing was conducted at the nCounter Core Facility Heidelberg using the nCounter SPRINT system and nCounter Elements chemistry as described (68). Probe design for the glycosyltransferase genes and reference genes are given on request. Data normalization was performed with the nSolver Analysis Software 3.0 (nanoString Technologies).

### Human phospho-kinase and MMP array

Phosphorylation of several kinases and expression levels of matrix metallopeptidases (MMPs) were analyzed using Proteome Profiler Human Phospho-Kinase Array Kit (R&D Systems) and Human MMP Antibody Array Kit (Abcam), respectively, according to the manufacturer’s instructions. Briefly, ~2.0 x 10^6^ cells were grown in complete media for 48 h, lysed and protein concentration was measured using the BCA protein assay (Pierce). For both arrays, pre-blocked nitrocellulose membranes were incubated with 500 μg of cellular extract o/n at 4 °C and subsequently washed and incubated with biotin-conjugated antibodies and HRP-conjugated streptavidin. Membranes were then incubated with Chemi Reagent Mix and developed in ImageQuant LAS 500 imaging system (GE Healthcare). For protein level quantification, the integrated optical density (IOD) of each array spot was quantified using ImageJ software (version 1.52i, NIH). IOD values were normalized to those of the positive controls on each membrane and corrected for background signal. Relative expression levels of proteins in two groups of cells were calculated as %. The relative amount of the respective protein in wild type was considered as 100%.

### Immunofluorescence

For actin staining, cells were seeded on poly-L-lysine-coated coverslips and fixed after 72 h with pre-warmed paraformaldehyde (3% in PBS) for 10 min. Subsequently, cells were washed with PBS and permeabilized for 5 min with 0.1% Triton X-100 in PBS and blocked with 5% ChemiBlocker (Merck-Millipore) in PBS for 30 min. Coverslips were incubated with Rhodamine Phalloidin (ThermoFisher Scientific) and DAPI (Sigma Aldrich) both at 1:1000 dilution in blocking solution for 1 h at 37 °C in a wet chamber, washed three times with PBS and mounted in Mowiol (Polysciences). Images were acquired using an Olympus IX81 inverted microscope (Olympus) equipped with a Spectra Xlight system (Lumencor), Dualband GFP/mCherry sbx ET and QuadDAPI/FITC/Cy3/Cy5 sbx HC filter sets, an UPLSAPO 20x/0.75 Air objective lens (Olympus), an ORCA-R2 camera (Hamamatsu) and CellSens dimension software (Olympus). Image brightness and contrast were linearly adjusted for publication using ImageJ software (version 1.52i, NIH).

### Statistical analysis

For statistical testing, two-sided paired Student’s t-tests were performed using Excel. Error bars represent standard deviations of at least three independent experiments unless indicated otherwise.

## Data availability

MS proteomics data have been deposited to the ProteomeXchange Consortium via the PRIDE (62) partner repository with the dataset identifier PXD021321. MS glycoproteomics raw data have been deposited to the MassIVE repository under the dataset identifier MSV000085243. Unprocessed xCGE-LIG *N*-glycocomics raw data are available upon request. All other data described are contained within the article.

## Acknowledgements

We are very grateful to Mark Lommel for cloning plasmid pMLHD7 and his help in the generation and characterization of TALEN-mediated knock-out cells. We thank the group of Joachim Wittbrodt for advice and support in TALEN technology; Kevin P. Campbell for providing the anti-β dystroglycan antibody; and Ricardo Carvalho and the ZMBH Imaging Facility (Heidelberg University; Germany) for excellent technical support in microscopy. We thank Udo Reichl for generously providing research infrastructure and support to ER.

## Author contributions

SIN, MFB, PW, TR, ER, CT and SS conceived and designed the experiments. SIN, MH, RH, PW, MFB, CH and NH performed the experiments. In particular: SIN performed adhesion and bead aggregation assays, qRT-PCR as well as kinase and MMP arrays. MH and RH performed glycoproteome and glycome analysis, PW mass spectrometry, MFB proliferation and migration assays, CH immunofluorescence and WGA pull-down assay, and NH nCounter gene expression profiling. SIN, MH, NR, PW, MFB, CH, RH, NH, ER and SS analyzed and evaluated data. SIN, NR and SS wrote and UR edited the manuscript.

## Funding

This work was funded by the Deutsche Forschungsgemeinschaft (DFG, German Research Foundation) – Project-ID STR 443/6-1 (to S. S.), RA2992/1-1 (to E. R.), RU 747/1-1 (to T. R.), TH1461/7-1 (to C. T.) – Forschungsgruppe FOR 2509.

## Conflicts of Interest

The authors declare no conflict of interest.

## Abbreviations

The abbreviations used are:

α/β-DG: α/β-Dystroglycan
E/N-Cdh: E(epithelial)/N(neural)-Cadherin
ECM: Extracellular Matrix
EMT: Epithelial-Mesenchymal Transition
ER: Endoplasmic Reticulum
HILIC SPE: Hydrophilic Interaction Liquid Chromatography Solid Phase Extraction
HPRT: Hypoxanthine-guanine Phosphoribosyltransferase
MEB disease: Muscle-Eye-Brain disease
MMPs: Matrix Metallopeptidases
qRT-PCR: Quantitative Real-Time PCR
RFLP: Restriction Fragment Length Polymorphism
RP-LC-ESI-OT MS/MS: Reverse-Phase Liquid Chromatography Coupled Online to Electrospray Ionization Orbitrap Tandem Mass Spectrometer
PCA: Principle Component Analysis
TALENs: Transcription Activator-Like Effector Nucleases
WGA: Wheat Germ Agglutinin
xCGE-LIF: Multiplexed Capillary Gel Electrophoresis with Laser Induced Fluorescence Detection

## Notes

### Competing Interest Statement

The authors have declared no competing interest.

## References

1. Corfield, A. P., and Berry, M. (2015) Glycan variation and evolution in the eukaryotes. Trends in Biochemical Sciences 40, 351–359

2. Moremen, K. W., Tiemeyer, M., and Nairn, A. V. (2012) Vertebrate protein glycosylation: diversity, synthesis and function. Nature Reviews Molecular Cell Biology 13, 448–462

3. Pereira, M. S., Alves, I., Vicente, M., Campar, A., Silva, M. C., Padrão, N. A., Pinto, V., Fernandes, Â., Dias, A. M., and Pinho, S. S. (2018) Glycans as key checkpoints of T cell activity and function. Frontiers in Immunology 9, 2754

4. Pinho, S. S., and Reis, C. A. (2015) Glycosylation in cancer: mechanisms and clinical implications. Nature Reviews Cancer 15, 540–555

5. Theodore, M., and Morava, E. (2011) Congenital disorders of glycosylation: sweet news. Curr Opin Pediatr 23, 581–587

6. Zielinska, D. F., Gnad, F., Wisniewski, J., and Mann, M. (2010) Precision mapping of an in vivo *N*-glycoproteome reveals rigid topological and sequence constraints. Cell 141, 897–907

7. Neubert, P., and Strahl, S. (2016) Protein *O*-mannosylation in the early secretory pathway. Curr Opin Cell Biol 41, 100–108

8. Endo, T. (2019) Mammalian *O*-mannosyl glycans: Biochemistry and glycopathology. Proc Jpn Acad Ser B Phys Biol Sci 95, 39–51

9. Yoshida-Moriguchi, T., and Campbell, K. P. (2015) Matriglycan: a novel polysaccharide that links dystroglycan to the basement membrane. Glycobiology 25, 702–713

10. Endo, T. (2014) Glycobiology of α-dystroglycan and muscular dystrophy. The Journal of Biochemistry 157, 1–12

11. Yoshida, A., Kobayashi, K., Manya, H., Taniguchi, K., Kano, H., Mizuno, M., Inazu, T., Mitsuhashi, H., Takahashi, S., Takeuchi, M., Herrmann, R., Straub, V., Talim, B., Voit, T., Topaloglu, H., Toda, T., and Endo, T. (2001) Muscular dystrophy and neuronal migration disorder caused by mutations in a glycosyltransferase, POMGnT1. Developmental Cell 1, 717–724

12. Xiong, H., Kobayashi, K., Tachikawa, M., Manya, H., Takeda, S., Chiyonobu, T., Fujikake, N., Wang, F., Nishimoto, A., Morris, G. E., Nagai, Y., Kanagawa, M., Endo, T., and Toda, T. (2006) Molecular interaction between fukutin and POMGnT1 in the glycosylation pathway of α-dystroglycan. Biochemical and Biophysical Research Communications 350, 935–941

13. Kuwabara, N., Manya, H., Yamada, T., Tateno, H., Kanagawa, M., Kobayashi, K., Akasaka-Manya, K., Hirose, Y., Mizuno, M., Ikeguchi, M., Toda, T., Hirabayashi, J., Senda, T., Endo, T., and Kato, R. (2016) Carbohydrate-binding domain of the POMGnT1 stem region modulates *O*-mannosylation sites of α-dystroglycan. Proc Natl Acad Sci U S A 113, 9280–9285

14. Liu, J., Ball, S. L., Yang, Y., Mei, P., Zhang, L., Shi, H., Kaminski, H. J., Lemmon, V. P., and Hu, H. (2006) A genetic model for muscle-eye-brain disease in mice lacking protein *O*-mannose 1,2-N-acetylglucosaminyltransferase (POMGnT1). Mech Dev 123, 228–240

15. Hu, H., Yang, Y., Eade, A., Xiong, Y., and Qi, Y. (2007) Breaches of the pial basement membrane and disappearance of the glia limitans during development underlie the cortical lamination defect in the mouse model of muscle-eye-brain disease. J Comp Neurol 501, 168–183

16. Lan, J., Guo, P., Lin, Y., Mao, Q., Guo, L., Ge, J., Li, X., Jiang, J., Lin, X., and Qiu, Y. (2015) Role of glycosyltransferase PomGnT1 in glioblastoma progression. Neuro Oncol 17, 211–222

17. Liu, Q., Xue, Y., Chen, Q., Chen, H., Zhang, X., Wang, L., Han, C., Que, S., Lou, M., and Lan, J. (2017) PomGnT1 enhances temozolomide resistance by activating epithelial-mesenchymal transition signaling in glioblastoma. Oncol Rep 38, 2911–2918

18. Larsen, I. S. B., Narimatsu, Y., Joshi, H. J., Siukstaite, L., Harrison, O. J., Brasch, J., Goodman, K. M., Hansen, L., Shapiro, L., Honig, B., Vakhrushev, S. Y., Clausen, H., and Halim, A. (2017) Discovery of an *O*-mannosylation pathway selectively serving cadherins and protocadherins. Proc Natl Acad Sci U S A 114, 11163–11168

19. Narimatsu, Y., Joshi, H. J., Schjoldager, K. T., Hintze, J., Halim, A., Steentoft, C., Nason, R., Mandel, U., Bennett, E. P., Clausen, H., and Vakhrushev, S. Y. (2019) Exploring regulation of protein *O*-glycosylation in isogenic human HEK293 cells by differential *O*-glycoproteomics. Mol Cell Proteomics 18, 1396–1409

20. Wheelock, M. J., and Johnson, K. R. (2003) Cadherins as modulators of cellular phenotype. Annu Rev Cell Dev Biol 19, 207–235

21. Guo, H. B., Johnson, H., Randolph, M., and Pierce, M. (2009) Regulation of homotypic cell-cell adhesion by branched N-glycosylation of N-cadherin extracellular EC2 and EC3 domains. J Biol Chem 284, 34986–34997

22. Langer, M. D., Guo, H., Shashikanth, N., Pierce, J. M., and Leckband, D. E. (2012) *N*-glycosylation alters cadherin-mediated intercellular binding kinetics. J Cell Sci 125, 2478–2485

23. Geiss, G. K., Bumgarner, R. E., Birditt, B., Dahl, T., Dowidar, N., Dunaway, D. L., Fell, H. P., Ferree, S., George, R. D., Grogan, T., James, J. J., Maysuria, M., Mitton, J. D., Oliveri, P., Osborn, J. L., Peng, T., Ratcliffe, A. L., Webster, P. J., Davidson, E. H., Hood, L., and Dimitrov, K. (2008) Direct multiplexed measurement of gene expression with color-coded probe pairs. Nat Biotechnol 26, 317–325

24. Winterhalter, P. R., Lommel, M., Ruppert, T., and Strahl, S. (2013) *O*-glycosylation of the non-canonical T-cadherin from rabbit skeletal muscle by single mannose residues. FEBS Letters 587, 3715–3721

25. Vester-Christensen, M. B., Halim, A., Joshi, H. J., Steentoft, C., Bennett, E. P., Levery, S. B., Vakhrushev, S. Y., and Clausen, H. (2013) Mining the *O*-mannose glycoproteome reveals cadherins as major *O*-mannosylated glycoproteins. Proc Natl Acad Sci U S A 110, 21018–21023

26. Emond, M. R., and Jontes, J. D. (2014) Bead aggregation assays for the characterization of putative cell adhesion molecules. J Vis Exp 92, e51762

27. Spence, H. J., Dhillon, A. S., James, M., and Winder, S. J. (2004) Dystroglycan, a scaffold for the ERK-MAP kinase cascade. EMBO Rep 5, 484–489

28. Bao, X., Kobayashi, M., Hatakeyama, S., Angata, K., Gullberg, D., Nakayama, J., Fukuda, M. N., and Fukuda, M. (2009) Tumor suppressor function of laminin-binding alpha-dystroglycan requires a distinct beta3-N-acetylglucosaminyltransferase. Proc Natl Acad Sci U S A 106, 12109–12114

29. Lavoie, H., Gagnon, J., and Therrien, M. (2020) ERK signalling: a master regulator of cell behaviour, life and fate. Nat Rev Mol Cell Biol. 10.1038/s41580-020-0255-7

30. Wheelock, M. J., Shintani, Y., Maeda, M., Fukumoto, Y., and Johnson, K. R. (2008) Cadherin switching. Journal of Cell Science 121, 727

31. Li, X., Wang, X., Tan, Z., Chen, S., and Guan, F. (2016) Role of glycans in cancer cells undergoing epithelial-mesenchymal transition. Front Oncol 6, 33

32. Olea-Flores, M., Zuñiga-Eulogio, M. D., Mendoza-Catalán, M. A., Rodríguez-Ruiz, H. A., Castañeda-Saucedo, E., Ortuño-Pineda, C., Padilla-Benavides, T., and Navarro-Tito, N. (2019) Extracellular-signal regulated kinase: A central molecule driving epithelial-mesenchymal transition in cancer. Int J Mol Sci 20, 2885

33. Lamouille, S., Xu, J., and Derynck, R. (2014) Molecular mechanisms of epithelial–mesenchymal transition. Nature Reviews Molecular Cell Biology 15, 178–196

34. Shay, G., Lynch, C. C., and Fingleton, B. (2015) Moving targets: Emerging roles for MMPs in cancer progression and metastasis. Matrix Biol 44-46, 200–206

35. Arpino, V., Brock, M., and Gill, S. (2015) The role of TIMPs in regulation of extracellular matrix proteolysis. Matrix Biol. 44-46, 247–254

36. Lommel, M., Winterhalter, P. R., Willer, T., Dahlhoff, M., Schneider, M. R., Bartels, M. F., Renner-Müller, I., Ruppert, T., Wolf, E., and Strahl, S. (2013) Protein *O*-mannosylation is crucial for E-cadherin–mediated cell adhesion. Proceedings of the National Academy of Sciences 110, 21024–21029

37. Carvalho, S., Oliveira, T., Bartels, M. F., Miyoshi, E., Pierce, M., Taniguchi, N., Carneiro, F., Seruca, R., Reis, C. A., Strahl, S., and Pinho, S. S. (2016) *O*-mannosylation and N-glycosylation: two coordinated mechanisms regulating the tumour suppressor functions of E-cadherin in cancer. Oncotarget 7, 65231–65246

38. Abbott, K. L., Troupe, K., Lee, I., and Pierce, M. (2006) Integrin-dependent neuroblastoma cell adhesion and migration on laminin is regulated by expression levels of two enzymes in the O-mannosyl-linked glycosylation pathway, PomGnT1 and GnT-Vb. Exp Cell Res 312, 2837–2850

39. Khoder-Agha, F., Harrus, D., Brysbaert, G., Lensink, M. F., Harduin-Lepers, A., Glumoff, T., and Kellokumpu, S. (2019) Assembly of B4GALT1/ST6GAL1 heteromers in the Golgi membranes involves lateral interactions via highly charged surface domains. Journal of Biological Chemistry 294, 14383–14393

40. Graham, J. B., Sunryd, J. C., Mathavan, K., Weir, E., Larsen, I. S. B., Halim, A., Clausen, H., Cousin, H., Alfandari, D., and Hebert, D. N. (2020) Endoplasmic reticulum transmembrane protein TMTC3 contributes to O-mannosylation of E-cadherin, cellular adherence, and embryonic gastrulation. Mol Biol Cell 31, 167–183

41. Acloque, H., Adams, M. S., Fishwick, K., Bronner-Fraser, M., and Nieto, M. A. (2009) Epithelial-mesenchymal transitions: the importance of changing cell state in development and disease. J Clin Invest 119, 1438–1449

42. Thiery, J. P. (2002) Epithelial–mesenchymal transitions in tumour progression. Nature Reviews Cancer 2, 442–454

43. Zeisberg, E. M., Tarnavski, O., Zeisberg, M., Dorfman, A. L., McMullen, J. R., Gustafsson, E., Chandraker, A., Yuan, X., Pu, W. T., Roberts, A. B., Neilson, E. G., Sayegh, M. H., Izumo, S., and Kalluri, R. (2007) Endothelial-to-mesenchymal transition contributes to cardiac fibrosis. Nat Med 13, 952–961

44. Mani, S. A., Guo, W., Liao, M. J., Eaton, E. N., Ayyanan, A., Zhou, A. Y., Brooks, M., Reinhard, F., Zhang, C. C., Shipitsin, M., Campbell, L. L., Polyak, K., Brisken, C., Yang, J., and Weinberg, R. A. (2008) The epithelial-mesenchymal transition generates cells with properties of stem cells. Cell 133, 704–715

45. Haensel, D., and Dai, X. (2018) Epithelial-to-mesenchymal transition in cutaneous wound healing: Where we are and where we are heading. Dev Dyn 247, 473–480

46. Nieto, M. A., Huang, R. Y., Jackson, R. A., and Thiery, J. P. (2016) EMT: 2016. Cell 166, 21–45

47. Lou, Y., Preobrazhenska, O., auf dem Keller, U., Sutcliffe, M., Barclay, L., McDonald, P. C., Roskelley, C., Overall, C. M., and Dedhar, S. (2008) Epithelial-mesenchymal transition (EMT) is not sufficient for spontaneous murine breast cancer metastasis. Dev Dyn 237, 2755–2768

48. Schaeffer, D., Somarelli, J. A., Hanna, G., Palmer, G. M., and Garcia-Blanco, M. A. (2014) Cellular migration and invasion uncoupled: increased migration is not an inexorable consequence of epithelial-to-mesenchymal transition. Mol Cell Biol 34, 3486–3499

49. Van Hoof, J., and Harrisson, F. (1986) Interaction between epithelial basement membrane and migrating mesoblast cells in the avian blastoderm. Differentiation 32, 120–124

50. Nakaya, Y., Sukowati, E. W., Alev, C., Nakazawa, F., and Sheng, G. (2011) Involvement of dystroglycan in epithelial-mesenchymal transition during chick gastrulation. Cells Tissues Organs 193, 64–73

51. LeBert, D. C., Squirrell, J. M., Rindy, J., Broadbridge, E., Lui, Y., Zakrzewska, A., Eliceiri, K. W., Meijer, A. H., and Huttenlocher, A. (2015) Matrix metalloproteinase 9 modulates collagen matrices and wound repair. Development 142, 2136–2146

52. Bozzi, M., Sciandra, F., and Brancaccio, A. (2015) Role of gelatinases in pathological and physiological processes involving the dystrophin-glycoprotein complex. Matrix Biol 44-46, 130–137

53. Gui, T., Sun, Y., Shimokado, A., and Muragaki, Y. (2012) The roles of mitogen-activated protein kinase pathways in TGF-β-induced epithelial-mesenchymal transition. J Signal Transduct 2012, 289243

54. Huang, Q., Miller, M. R., Schappet, J., and Henry, M. D. (2015) The glycosyltransferase LARGE2 is repressed by Snail and ZEB1 in prostate cancer. Cancer Biol Ther 16, 125–136

55. Day, B. W., Lathia, J. D., Bruce, Z. C., D’Souza, R. C. J., Baumgartner, U., Ensbey, K. S., Lim, Y. C., Stringer, B. W., Akgül, S., Offenhäuser, C., Li, Y., Jamieson, P. R., Smith, F. M., Jurd, C. L. R., Robertson, T., Inglis, P. L., Lwin, Z., Jeffree, R. L., Johns, T. G., Bhat, K. P. L., Rich, J. N., Campbell, K. P., and Boyd, A. W. (2019) The dystroglycan receptor maintains glioma stem cells in the vascular niche. Acta Neuropathol 138, 1033–1052

56. Doyle, E. L., Booher, N. J., Standage, D. S., Voytas, D. F., Brendel, V. P., Vandyk, J. K., and Bogdanove, A. J. (2012) TAL Effector-Nucleotide Targeter (TALE-NT) 2.0: tools for TAL effector design and target prediction. Nucleic Acids Res 40, W117–W122

57. Cermak, T., Doyle, E. L., Christian, M., Wang, L., Zhang, Y., Schmidt, C., Baller, J. A., Somia, N. V., Bogdanove, A. J., and Voytas, D. F. (2011) Efficient design and assembly of custom TALEN and other TAL effector-based constructs for DNA targeting. Nucleic Acids Res 39, e82

58. Michele, D. E., Barresi, R., Kanagawa, M., Saito, F., Cohn, R. D., Satz, J. S., Dollar, J., Nishino, I., Kelley, R. I., Somer, H., Straub, V., Mathews, K. D., Moore, S. A., and Campbell, K. P. (2002) Post-translational disruption of dystroglycan–ligand interactions in congenital muscular dystrophies. Nature 418, 417–421

59. Kwon, M., and Firestein, B. L. (2013) DNA transfection: calcium phosphate method. Methods Mol Biol 1018, 107–110

60. Cox, J., and Mann, M. (2008) MaxQuant enables high peptide identification rates, individualized p.p.b.-range mass accuracies and proteome-wide protein quantification. Nature biotechnology 26, 1367–1372

61. Cox, J., Neuhauser, N., Michalski, A., Scheltema, R. A., Olsen, J. V., and Mann, M. (2011) Andromeda: a peptide search engine integrated into the MaxQuant environment. J Proteome Res 10, 1794–1805

62. Vizcaíno, J. A., Côté, R. G., Csordas, A., Dianes, J. A., Fabregat, A., Foster, J. M., Griss, J., Alpi, E., Birim, M., Contell, J., O’Kelly, G., Schoenegger, A., Ovelleiro, D., Pérez-Riverol, Y., Reisinger, F., Ríos, D., Wang, R., and Hermjakob, H. (2013) The PRoteomics IDEntifications (PRIDE) database and associated tools: status in 2013. Nucleic Acids Res 41, D1063–1069

63. Hoffmann, M., Pioch, M., Pralow, A., Hennig, R., Kottler, R., Reichl, U., and Rapp, E. (2018) The fine art of destruction: A guide to in-depth glycoproteomic analyses-exploiting the diagnostic potential of fragment ions. Proteomics 18, e1800282

64. Pioch, M., Hoffmann, M., Pralow, A., Reichl, U., and Rapp, E. (2018) glyXtoolMS: An open-source pipeline for semiautomated analysis of glycopeptide mass spectrometry data. Analytical Chemistry 90, 11908–11916

65. Ruhaak, L. R., Hennig, R., Huhn, C., Borowiak, M., Dolhain, R. J., Deelder, A. M., Rapp, E., and Wuhrer, M. (2010) Optimized workflow for preparation of APTS-labeled *N*-glycans allowing high-throughput analysis of human plasma glycomes using 48-channel multiplexed CGE-LIF. J Proteome Res 9, 6655–6664

66. Hennig, R., Cajic, S., Borowiak, M., Hoffmann, M., Kottler, R., Reichl, U., and Rapp, E. (2016) Towards personalized diagnostics via longitudinal study of the human plasma *N*-glycome. Biochim Biophys Acta 1860, 1728–1738

67. Konze, S. A., Cajic, S., Oberbeck, A., Hennig, R., Pich, A., Rapp, E., and Buettner, F. F. R. (2017) Quantitative assessment of sialo-glycoproteins and *N*-glycans during cardiomyogenic differentiation of human induced pluripotent stem cells. Chembiochem 18, 1317–1331

68. Thoeni, C., Waldherr, R., Scheuerer, J., Schmitteckert, S., Roeth, R., Niesler, B., Cutz, E., Flechtenmacher, C., Goeppert, B., Schirmacher, P., and Lasitschka, F. (2019) Expression analysis of ATP-binding cassette transporters ABCB11 and ABCB4 in primary sclerosing cholangitis and variety of pediatric and adult cholestatic and noncholestatic liver diseases. Canadian Journal of Gastroenterology and Hepatology, 10.1155/2019/1085717

