## Supporting Information for "Glycosyltransferase POMGNT1 deficiency affects N-cadherin-mediated cell-cell adhesion"

From the [1]Centre for Organismal Studies (COS), Glycobiology, Heidelberg University, Heidelberg, Germany, the [2]Max Planck Institute for Dynamics of Complex Technical Systems, Bioprocess Engineering, Magdeburg, Germany, the [3]glyXera GmbH, Magdeburg, Germany, the [4]Center for Child and Adolescent Medicine, Department Pediatrics I, University of Heidelberg, Heidelberg, Germany, the [5]Center for Molecular Biology of Heidelberg University (ZMBH), DKFZ-ZMBH Alliance, Heidelberg, Germany

[#] Present address: Clinic for Heart Surgery, Martin-Luther-University Halle-Wittenberg, Halle/Saale, Germany

*Corresponding author: Sabine Strahl

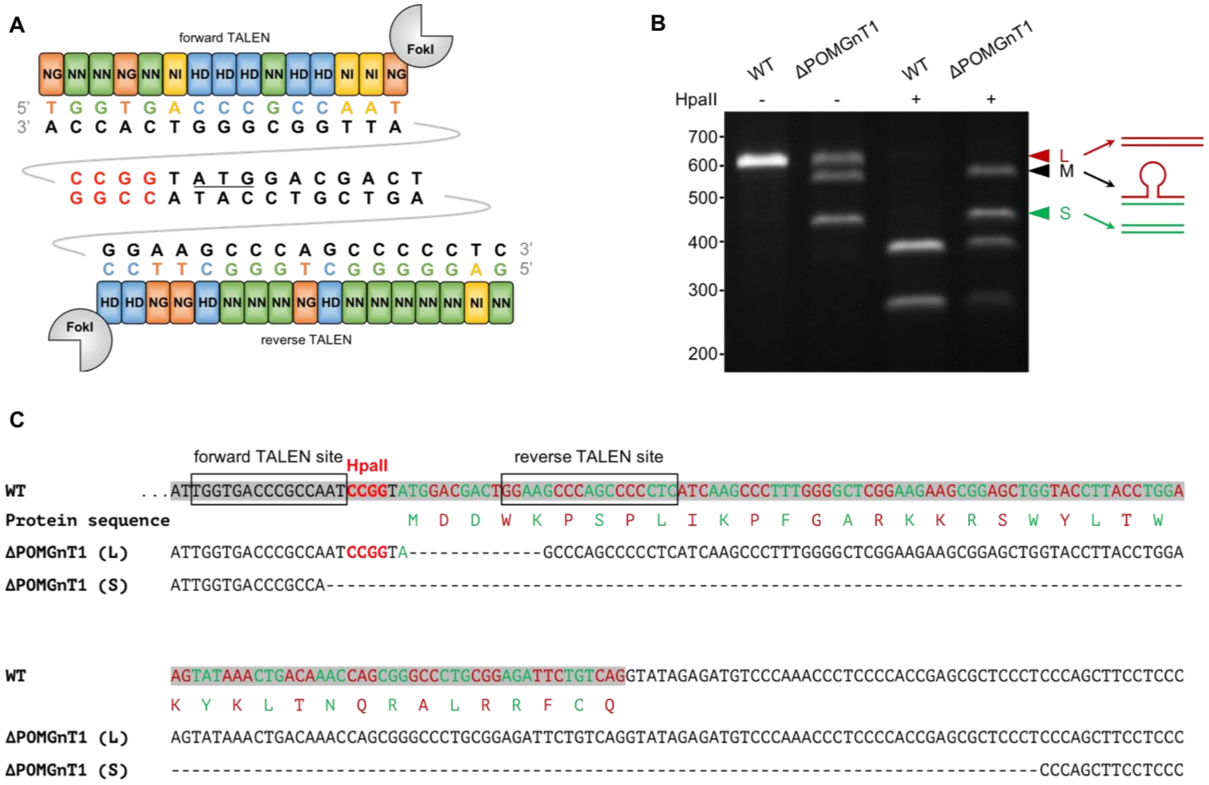

**Figure S1. Genetic verification of TALEN-mediated POMGNT1 disruption.** (A) Scheme depicting the repeat variable di-residue composition of the TALEN pair used for *POMGNT1* disruption and its target sequences surrounding the ATG start codon (underlined) of the *POMGNT1* gene. The unique *HpaII* restriction site within the TALEN-targeted region (in red) was used to screen the targeted locus for the induction of mutations by RFLP analysis. Upon assembly and sequence verification (see Experimental procedures for details) pC-Goldy plasmids encoding forward and reverse TALEN were cotransfected into wild type HEK293T cells. Upon genomic DNA extraction and PCR, *HpaII* resistant PCR products indicated efficient TALEN-mediated cleavage. Single cell knock-outs were obtained upon dilutional cloning and analyzed by PCR and RFLP analysis (shown in B). (B) Screening PCR and RFLP analysis performed on DNA extracted from wild type and ΔPOMGnT1 cells. PCR on genomic DNA from wild type cells reveals a single *HpaII* sensitive PCR product whereas PCR on genomic DNA from ΔPOMGnT1 cells results in three distinct PCR products. Restriction with *HpaII* reveals PCR products M and S (for middle and small size, respectively) to be *HpaII* insensitive, whereas PCR product L (for large) is sensitive to *HpaII*. (C) Sequence of PCR product obtained in wild type HEK293T cells (accompanied by its protein sequence) and PCR products L and S detected in ΔPOMGnT1 cells. TALEN target sequences are depicted in boxes. Sequencing results for fragment L (ΔPOMGnT1 L) reveal a deletion of 13 bp immediately downstream of the *HpaII* site (in red) explaining its *HpaII* insensitivity. The sequence of fragment S (ΔPOMGnT1 S) shows a deletion of 167 bp disturbing the *HpaII* restriction site. In both cases deletions target the POMGNT1 start codon localized within exon 3 of the *POMGNT1* gene (highlighted in grey). For fragment M that shows an intermediate size and is *HpaII* sensitive sequencing results reveal that it is a hybrid of the other two fragments that occurs during PCR analysis due to high sequence overlap. Disruption of the *POMGNT1* start codon accompanied by the absence of an alternative start codon in close proximity preclude *POMGNT1* gene expression. WT: wild type.

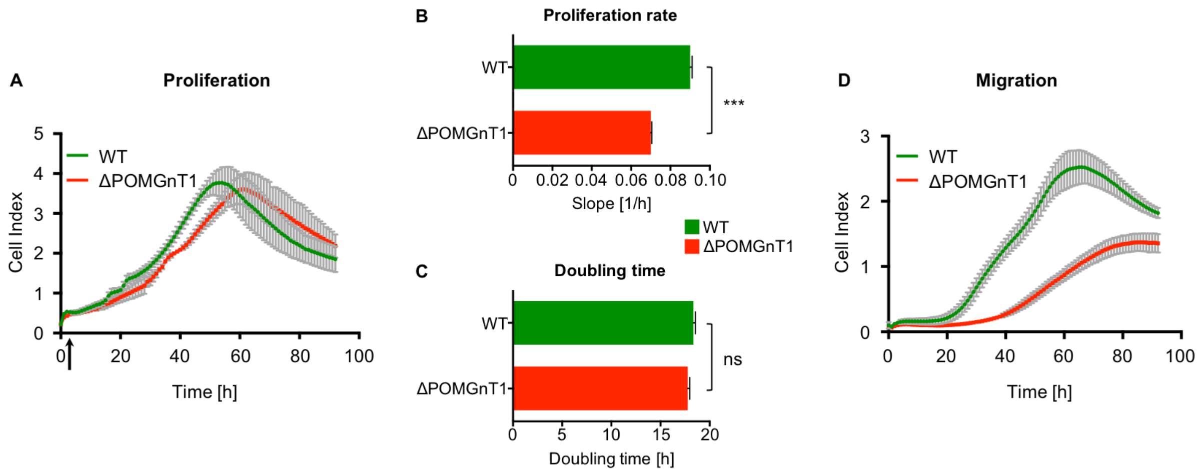

**Figure S2. POMGNT1 deficiency affects cell physiology and migration of HEK293T cells.** (A) XCELLigence-mediated real-time monitoring of differences in cell physiology between WT (green line) and ΔPOMGnT1 cells (red line). The unitless Cell Index calculated as the change of impedance between time point n and in the absence of cells monitors cell physiology in terms of cell number, proliferation rate, cell size/shape and cell-substrate attachment. 2.0 x $10^4$ cells/well were seeded and resulted in an initial peak of the Cell Index (depicted by the arrow) due to cells attaching to the well surface followed by an exponential phase (25-56 h). An exemplary run is shown. Proliferation rates (using the slope as a unit) and doubling time of WT and ΔPOMGnT1 cells were calculated during exponential growth and are depicted in B and C respectively. (B) Relative comparison of the slope (as a unit of the proliferation rate) during exponential growth between WT and ΔPOMGnT1 cells showing decreased proliferation for ΔPOMGnT1 cells. (C) Relative comparison of the doubling time showing no significant difference between WT and ΔPOMGnT1 cells during exponential growth. The experiment was performed 3 times with at least 3 technical replicates per run yielding similar results. Data are presented as means ± SD. Asterisks denote statistical significance in comparison to WT cells: *** $p \leq 0.001$, *n. s*: not significant. (D) XCELLigence-mediated monitoring of differences in migration between WT (green line) and ΔPOMGnT1 cells (red line). 7.5 x $10^3$ cells were seeded in FBS-free medium in the upper compartment of a trans-well compartment (see Experimental procedures for details). The lower compartment contained serum (10% (v/v)). Cells migrating through the porous membrane settled down on the surface facing the FBS-containing medium and started exponential growth after 19 h. Results show that ΔPOMGnT1 cells migrate significantly slower compared to WT cells within the exponential phase (19-64 h). An exemplary run is shown. The experiment was repeated 2 times resulting in a similar outcome.

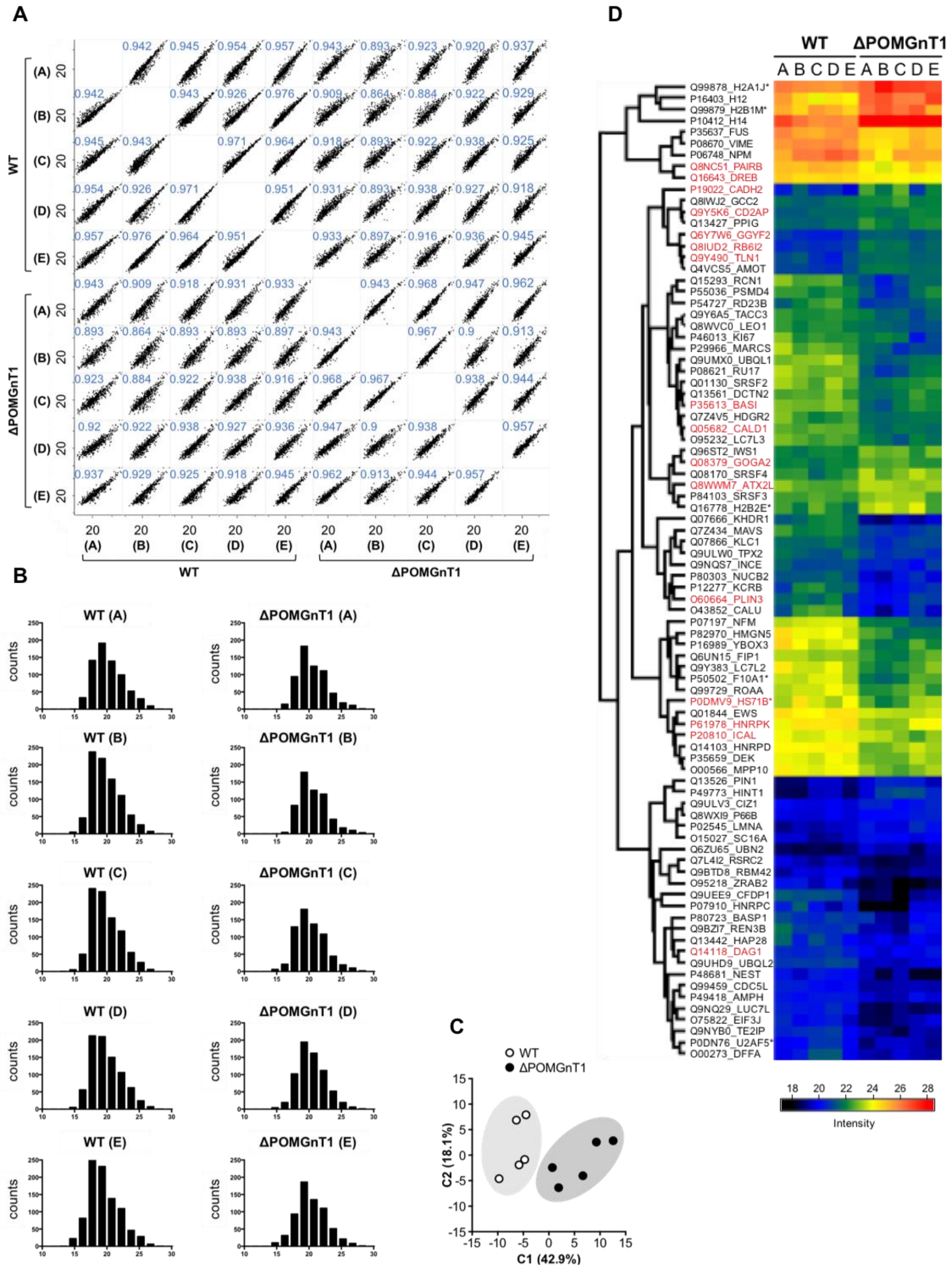
A
WT
ΔPOMGnT1
(A) (B) (C) (D) (E)
20
0.942 0.945 0.954 0.957 0.943 0.893 0.923 0.920 0.937
0.942 0.943 0.926 0.976 0.909 0.864 0.884 0.922 0.929
0.945 0.943 0.971 0.964 0.918 0.893 0.922 0.938 0.925
0.954 0.926 0.971 0.951 0.931 0.893 0.938 0.927 0.918
0.957 0.976 0.964 0.951 0.933 0.897 0.916 0.936 0.945
0.943 0.909 0.918 0.931 0.933 0.943 0.968 0.947 0.962
0.893 0.864 0.893 0.893 0.897 0.943 0.967 0.9 0.913
0.923 0.884 0.922 0.938 0.916 0.968 0.967 0.938 0.944
0.92 0.922 0.938 0.927 0.936 0.947 0.9 0.938 0.957
0.937 0.929 0.925 0.918 0.945 0.962 0.913 0.944 0.957
B
WT (A)
ΔPOMGnT1 (A)
WT (B)
ΔPOMGnT1 (B)
WT (C)
ΔPOMGnT1 (C)
WT (D)
ΔPOMGnT1 (D)
WT (E)
ΔPOMGnT1 (E)
counts
C
○ WT
● ΔPOMGnT1
C2 (18.1%)
C1 (42.9%)
D
WT
ΔPOMGnT1
A B C D E A B C D E
Q99878_H2A1J*
P16403_H12
Q99879_H2B1M*
P10412_H14
P35637_FUS
P08670_VIME
P06748_NPM
Q8NC51_PAIRB
Q16643_DREB
P19022_CADH2
Q8IWJ2_GCC2
Q9Y5K6_CD2AP
Q13427_PPIG
Q6Y7W6_GGYF2
Q8IUD2_RB6I2
Q9Y490_TLN1
Q4VCS5_AMOT
Q15293_RCN1
P55036_PSMD4
P54727_RD23B
Q9Y6A5_TACC3
Q8WVC0_LEO1
P46013_KI67
P29966_MARCS
Q9UMX0_UBQL1
P08621_RU17
Q01130_SRSF2
Q13561_DCTN2
P35613_BASI
Q7Z4V5_HDGR2
Q05682_CALD1
O95232_LC7L3
Q96ST2_IWS1
Q08379_GOGA2
Q08170_SRSF4
Q8WWM7_ATX2L
P84103_SRSF3
Q16778_H2B2E*
Q07666_KHDR1
Q7Z434_MAVS
Q07866_KLC1
Q9ULW0_TPX2
Q9NQS7_INCE
P80303_NUCB2
P12277_KCRB
O60664_PLIN3
O43852_CALU
P07197_NFM
P82970_HMGN5
P16989_YBOX3
Q6UN15_FIP1
Q9Y383_LC7L2
P50502_F10A1*
Q99729_ROAA
P0DMV9_HS71B*
Q01844_EWS
P61978_HNRPK
P20810_ICAL
Q14103_HNRPD
P35659_DEK
O00566_MPP10
Q13526_PIN1
P49773_HINT1
Q9ULV3_CIZ1
Q8WXI9_P66B
P02545_LMNA
O15027_SC16A
Q6ZU65_UBN2
Q7L4I2_RSRC2
Q9BTD8_RBM42
O95218_ZRAB2
Q9UEE9_CFDP1
P07910_HNRPC
P80723_BASP1
Q9BZI7_REN3B
Q13442_HAP28
Q14118_DAG1
Q9UHD9_UBQL2
P48681_NEST
Q99459_CDC5L
P49418_AMPH
Q9NQ29_LUC7L
O75822_EIF3J
Q9NYB0_TE2IP
P0DN76_U2AF5*
O00273_DFFA
18 20 22 24 26 28
Intensity

**Figure S3. Quality assessment of quantitative proteomics performed on wild type and POMGNT1-deficient HEK293T cells.** (A) Scatter plots of log2 transformed LFQ-protein abundances of five replicates from wild type and ΔPOMGnT1 cells plotted against each other on the x- and y-axis, respectively (raw data has been deposited to the ProteomeXchange Consortium via the PRIDE partner repository with the dataset identifier PXD021321). For comparison of linear correlation between the variables x and y, measurements should yield identical protein intensity, exemplified by their Pearson correlation coefficient which ideally is at 1. Based on the multi scatter plot and Pearson correlation, homoscedasticity of the dataset could be assumed. (B) Histograms showing on the x-axis the log2 transformed LFQ-protein intensities divided in several windows (+/- 0.75) starting with an intensity of 8.75 up to 31.25, whereas the y-axis shows the number of proteins in a given window of the x-axis (Table S2). Therefore, an approximate normal distribution of the five replicates from wild type and ΔPOMGnT1 cells was shown. With homoscedasticity and normal distribution, the basic assumptions for the applied Student's t-test were given. (C) The principle component analysis (PCA) of five replicates from wild type and ΔPOMGnT1 cells proteomics data (log2 transformed LFQ-protein intensities). The PCA plot represents 437 proteins that indicated proteomics profile similarity within wild type or ΔPOMGnT1 but also clear differences between the groups. The x- and y-axis plot the two major components (C1 and C2) and the percentage of variances explained by these components (Table S3). (D) Heat map showing the 86 proteins from Fig. 2A with differential expression levels between wild type and ΔPOMGnT1 cells for each individual replicate in rainbow colors. Genes in red cluster under the molecular function term of "cadherin binding involved in cell-cell adhesion" upon gene ontology term functional annotation. Stars depict cases in which several proteins were detected. Only one protein is shown here (Table S4). LFQ-intensities of the proteins associated with the depicted color scale are summarized in Table S5. WT: wild type.

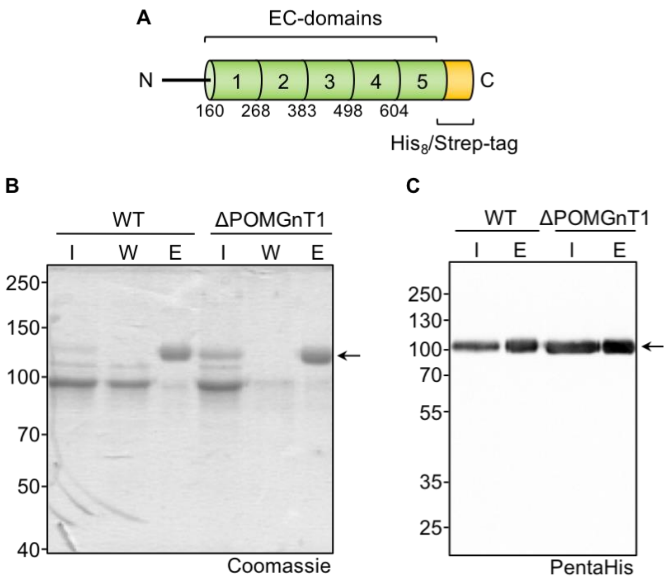

**Figure S4. Purification of the extracellular portion of N-Cdh from HEK293T cells.** (A) Scheme depicting the expressed N-Cdh fusion protein that consists of its five ectodomains C-terminally fused to a $His_8$/Strep-tag. The fusion protein was expressed from pMLHD7, stably transfected into HEK293T cells und purified upon secretion into the media using Ni-NTA IMAC (see Experimental procedures for details). (B, C) Analysis of the fusion protein after Ni-NTA purification by SDS-PAGE and Coomassie staining (B) or Western Blot (C). (B) Coomassie stained gel showing the protein content of the input (I), wash (W) and eluate (E) fractions of the purification from 60 ml culture supernatant. 1.4 µg protein from the eluate fraction were loaded, equaling to a 120-fold concentration from input to eluate. (C) Western Blot showing the N-Cdh fusion protein within the input (I) and eluate (E) fractions detected with an anti-PentaHis antibody. 300 ng protein from the eluate fraction was loaded. Arrows point to recombinant N-Cdh fusion. Data are representative of several independent purifications. WT: wild type.

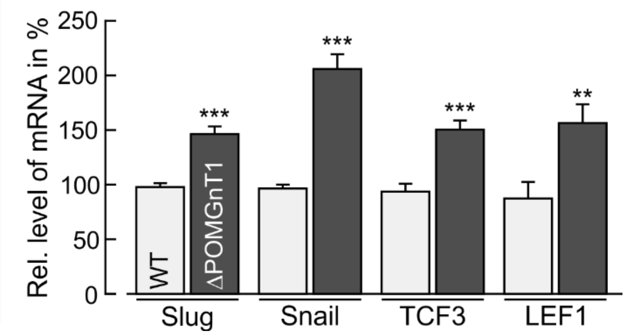

**Figure S5. POMGNT1 deficiency induces changes in the gene expression of EMT-inducing transcription factors in HEK293T cells.** Comparative gene expression analysis by qRT-PCR of the indicated EMT-inducing transcription factors between WT and ΔPOMGnT1 cells. Transcript levels are calculated in %, considering transcript levels of respective genes in WT cells as 100%. HPRT was used for normalization. Assays were performed in duplicate with three dilutions from three biological replicates. Data are represented as means ± SD. Asterisks denote statistical significance in comparison to WT cells: ** $p \leq 0.01$, *** $p \leq 0.001$.

**Table S1: Primers used for qRT-PCR**

| Gene | Primer sequence (5′ → 3′) |
|---|---|
| *CDH2* | Forward: GGTGGAGGAGAAGAAGACCAG |
| | Reverse: GGCATCAGGCTCCACAGT |
| *LAMA2* | Forward: GGCAATCTGAATACACTCGTGAC |
| | Reverse: TGTGTTGGTCCTCTCAGCATCC |
| *CLDN6* | Forward: TTCATCGGCAACAGCATCGT |
| | Reverse: GGTTATAGAAGTCCCGGATGA |
| *ACTA2* | Forward: CTATGCCTCTGGACGCACAACT |
| | Reverse: CAGATCCAGACGCATGATGGCA |
| *VIM* | Forward: AGGCAAAGCAGGAGTCCACTGA |
| | Reverse: ATCTGGCGTTCCAGGGACTCAT |
| *FN* | Forward: TACACTGGGAACACTTACCG |
| | Reverse: CCAATCTTGTAGGACTGACC |
| *MMP1* | Forward: ATGCTGAAACCCTGAAGGTG |
| | Reverse: GAGCATCCCCTCCAATACCT |
| *MMP2* | Forward: CCCCAAAACGGACAAAGAG |
| | Reverse: CACGAGCAAAGGCATCATCC |
| *MMP3* | Forward: GTCTCTTTCACTCAGCCAAC |
| | Reverse: ATCAGGATTTCTCCCCTCAG |
| *MMP8* | Forward: TGGACCCAATGGAATCCTTGC |
| | Reverse: ATAGCCACTCAGAGCCCAGTA |
| *MMP17* | Forward: TCCAGATCGACTTCTCCAAG |
| | Reverse: CCACATGGCTTAACCCAATG |
| *MMP28* | Forward: AGGCATTCCTAGAGAAGTACG |
| | Reverse: CTTACGCCTCATTTTGGTCC |
| *TIMP1* | Forward: GACGGCCTTCTGCAATTCC |
| | Reverse: GTATAAGGTGGTCTGGTTGACTTCTG |
| *TIMP2* | Forward: GAGCCTGAACCACAGGTACCA |
| | Reverse: TCTGTGACCCAGTCCATCCA |
| *TIMP3* | Forward: CCAGGACGCCTTCTGCAA |
| | Reverse: CCCCTCCTTTACCAGCTTCTTC |
| *HPRT* | Forward: AGAATGTCTTGATTGTGGAAGA |
| | Reverse: ACCTTGACCATCTTTGGATTA |

Table S2: Protein LFQ intensity (Log 2) for each sample derived from WT and ΔPOMGnT1 cells

| | | Frequency of proteins in a given window X | | | | | | | | | |
|---|---|---|---|---|---|---|---|---|---|---|---|
| | | **A (WT)** | **B (WT)** | **C (WT)** | **D (WT)** | **E (WT)** | **A (ΔPOMGnT1)** | **B (ΔPOMGnT1)** | **C (ΔPOMGnT1)** | **D (ΔPOMGnT1)** | **E (ΔPOMGnT1)** |
| **Log2 protein intensity value X with a window of +/- 0.75** | **8.75** | 0 | 0 | 0 | 0 | 0 | 0 | 0 | 0 | 0 | 0 |
| | **10.25** | 0 | 0 | 0 | 0 | 0 | 0 | 0 | 0 | 0 | 0 |
| | **11.75** | 0 | 0 | 0 | 0 | 0 | 0 | 0 | 0 | 0 | 0 |
| | **13.25** | 0 | 0 | 1 | 0 | 1 | 0 | 0 | 0 | 0 | 0 |
| | **14.75** | 1 | 6 | 6 | 6 | 23 | 0 | 0 | 7 | 5 | 11 |
| | **16.25** | 34 | 47 | 51 | 57 | 82 | 15 | 5 | 22 | 20 | 32 |
| | **17.75** | 142 | 238 | 241 | 213 | 260 | 95 | 83 | 130 | 103 | 94 |
| | **19.25** | 192 | 219 | 232 | 211 | 232 | 183 | 179 | 181 | 195 | 187 |
| | **20.75** | 140 | 159 | 156 | 151 | 139 | 125 | 127 | 138 | 163 | 136 |
| | **22.25** | 100 | 112 | 118 | 107 | 110 | 112 | 116 | 109 | 113 | 111 |
| | **23.75** | 53 | 55 | 56 | 63 | 47 | 47 | 43 | 45 | 53 | 52 |
| | **25.25** | 31 | 27 | 27 | 25 | 29 | 19 | 18 | 20 | 20 | 22 |
| | **26.75** | 10 | 8 | 7 | 8 | 7 | 12 | 11 | 11 | 7 | 9 |
| | **28.25** | 1 | 1 | 0 | 1 | 1 | 2 | 4 | 3 | 2 | 2 |
| | **29.75** | 0 | 0 | 0 | 0 | 0 | 0 | 0 | 0 | 0 | 0 |
| | **31.25** | 0 | 0 | 0 | 0 | 0 | 0 | 0 | 0 | 0 | 0 |

**Table S3: Principle component analysis (PCA) of WT and ΔPOMGnT1 cells-derived proteomics data.**

| Samples | Component 1 (42.9%) | Component 2 (18.1%) | C: Group1 | T: Item |
|---|---|---|---|---|
| CTL (A) | -4.73211 | -0.84734 | 1 | CTL (A) |
| CTL (B) | -9.72909 | -4.61911 | 1 | CTL (B) |
| CTL (C) | -6.34981 | 6.87674 | 1 | CTL (C) |
| CTL (D) | -4.47574 | 7.91855 | 1 | CTL (D) |
| CTL (E) | -5.83496 | -1.91159 | 1 | CTL (E) |
| POMGNT1 -/- (A) | 6.65168 | -4.02594 | 2 | POMGNT1 -/- (A) |
| POMGNT1 -/- (B) | 12.5791 | 2.83931 | 2 | POMGNT1 -/- (B) |
| POMGNT1 -/- (C) | 9.2966 | 2.57187 | 2 | POMGNT1 -/- (C) |
| POMGNT1 -/- (D) | 0.681938 | -2.431 | 2 | POMGNT1 -/- (D) |
| POMGNT1 -/- (E) | 1.91239 | -6.37149 | 2 | POMGNT1 -/- (E) |

**Table S4: Differentially expressed proteins in *POMGNT1*-deficient cells.** Proteins highlighted in red are under the molecular function term of "cadherin binding involved in cell-cell adhesion" upon GO term functional annotation. Stars (*) depict cases in which not only one but several proteins could belong to the identified peptides. In the heat map in Fig. S3D only one protein name (the first in the row) is shown.

| Significance (-Log10 (P-value)) | Difference (Log2 fold change) | UniProt ID | Protein | |
|---|---|---|---|---|
| 3.021 | -1.199 | Q99878 | H2A1J* | H2A1H, H2A2A, H2A1D, H2A1, H2AJ, H2A2C |
| 3.126 | -1.887 | P16403 | H12 | |
| 2.002 | -1.077 | Q99879 | H2B1M* | H2B1N, H2B1H, H2B2F, , H2B1C, H2B1D, H2BFS, , H2B1K, H2B1L |
| 4.623 | -2.205 | P10412 | H14 | |
| 2.935 | 0.792 | P35637 | FUS | |
| 2.935 | 0.792 | P08670 | VIME | |
| 1.726 | 0.950 | P06748 | NPM | |
| 2.124 | 0.768 | Q8NC51 | PAIRB | |
| 3.222 | 0.508 | Q16643 | DREB | |
| 1.873 | -1.418 | P19022 | CADH2 | |
| 2.289 | -0.488 | Q8IWJ2 | GCC2 | |
| 2.675 | -0.434 | Q9Y5K6 | CD2AP | |
| 2.184 | -0.505 | Q13427 | PPIG | |
| 3.452 | -0.788 | Q6Y7W6 | GGYF2 | |
| 4.030 | -0.958 | Q8IUD2 | RB6I2 | |
| 1.886 | -0.633 | Q9Y490 | TLN1 | |
| 4.189 | -0.839 | Q4VCS5 | AMOT | |
| 2.540 | 1.373 | Q15293 | RCN1 | |
| 1.672 | 0.873 | P55036 | PSMD4 | |
| 1.963 | 0.883 | P54727 | RD23B | |
| 2.625 | 0.722 | Q9Y6A5 | TACC3 | |
| 3.017 | 0.741 | Q8WVC0 | LEO1 | |
| 1.518 | 0.979 | P46013 | KI67 | |
| 3.549 | 1.566 | P29966 | MARCS | |
| 4.853 | 1.444 | Q9UMX0 | UBQL1 | |
| 3.538 | 1.555 | P08621 | RU17 | |
| 2.661 | 0.927 | Q01130 | SRSF2 | |
| 2.219 | 0.716 | Q13561 | DCTN2 | |
| 3.165 | 1.083 | P35613 | BASI | |
| 2.298 | 0.707 | Q7Z4V5 | HDGR2 | |
| 5.608 | 0.948 | Q05682 | CALD1 | |
| 3.130 | 0.878 | O95232 | LC7L3 | |
| 2.908 | -0.769 | Q96ST2 | IWS1 | |

| | | | | |
|---|---|---|---|---|
| 2.511 | -0.525 | Q08379 | GOGA2 | |
| 2.719 | -0.990 | Q08170 | SRSF4 | |
| 2.479 | -0.530 | Q8WWM7 | ATX2L | |
| 2.023 | -0.561 | P84103 | SRSF3 | |
| 2.202 | -1.119 | Q16778 | H2B2E* | H2B1B, H2B1O, H2B1J, H2B |
| 5.456 | 2.497 | Q07666 | KHDR1 | |
| 4.206 | 1.194 | Q7Z434 | MAVS | |
| 3.218 | 0.712 | Q07866 | KLC1 | |
| 2.325 | 0.646 | Q9ULW0 | TPX2 | |
| 2.267 | 0.725 | Q9NQS7 | INCE | |
| 3.153 | 1.404 | P80303 | NUCB2 | |
| 2.674 | 1.401 | P12277 | KCRB | |
| 2.026 | 1.038 | O60664 | PLIN3 | |
| 2.357 | 1.933 | O43852 | CALU | |
| 5.166 | 1.933 | P07197 | NFM | |
| 4.047 | 1.896 | P82970 | HMGN5 | |
| 3.831 | 1.776 | P16989 | YBOX3 | |
| 2.958 | 0.927 | Q6UN15 | FIP1 | |
| 2.853 | 1.330 | Q9Y383 | LC7L2 | |
| 3.234 | 1.422 | P50502 | F10A1* | F10A5 |
| 3.818 | 1.165 | Q99729 | ROAA | |
| 1.613 | 0.975 | P0DMV9 | HS71B* | HS71A |
| 3.713 | 0.792 | Q01844 | EWS | |
| 2.005 | 0.829 | P61978 | HNRPK | |
| 1.813 | 0.631 | P20810 | ICAL | |
| 3.591 | 1.018 | Q14103 | HNRPD | |
| 3.119 | 0.768 | P35659 | DEK | |
| 3.178 | 0.929 | O00566 | MPP10 | |
| 1.997 | -0.983 | Q13526 | PIN1 | |
| 2.912 | -1.737 | P49773 | HINT1 | |
| 2.408 | -0.544 | Q9ULV3 | CIZ1 | |
| 2.718 | -0.484 | Q8WXI9 | P66B | |
| 2.117 | -0.764 | P02545 | LMNA | |
| 2.284 | -0.597 | O15027 | SC16A | |
| 1.859 | -0.600 | Q6ZU65 | UBN2 | |
| 2.357 | 0.651 | Q7L4I2 | RSRC2 | |
| 2.234 | 0.636 | Q9BTD8 | RBM42 | |
| 2.895 | 1.430 | O95218 | ZRAB2 | |
| 3.629 | 2.516 | Q9UEE9 | CFDP1 | |
| 1.737 | 1.754 | P07910 | HNRPC | |

| | | | | |
|---|---|---|---|---|
| 1.552 | 0.969 | P80723 | BASP1 | |
| 2.373 | 1.369 | Q9BZI7 | REN3B | |
| 1.935 | 0.997 | Q13442 | HAP28 | |
| 2.805 | 0.685 | Q9UHD9 | UBQL2 | |
| 6.030 | 1.357 | Q14118 | DAG1 | |
| 2.577 | 1.321 | P48681 | NEST | |
| 4.285 | 0.892 | Q99459 | CDC5L | |
| 2.442 | 0.654 | P49418 | AMPH | |
| 5.834 | 1.584 | Q9NQ29 | LUC7L | |
| 2.053 | 1.016 | O75822 | EIF3J | |
| 3.436 | 1.421 | Q9NYB0 | TE2IP | |
| 2.273 | 0.966 | P0DN76 | U2AF5* | U2AF1, U2AF4 |
| 1.953 | 1.165 | O00273 | DFFA | |

**Table S5: LFQ-intensities of the proteins associated with the depicted color scale.**

| | LFQ-Intensity | | | | | | | |
|---|---|---|---|---|---|---|---|---|
| | | | **colour** | **saturation** | **intensity** | **red** | **green** | **blue** |
| black | 18 | | 160 | 0 | 0 | 0 | 0 | 0 |
| blue | 20 | | 160 | 240 | 120 | 0 | 0 | 255 |
| green | 22 | | 100 | 240 | 60 | 0 | 128 | 64 |
| yellow | 24 | | 40 | 240 | 120 | 255 | 255 | 0 |
| orange | 26 | | 13 | 240 | 150 | 255 | 128 | 64 |
| red | 28 | | 0 | 240 | 120 | 255 | 0 | 0 |
| start | 17.228 | also black | | | | | | |
| end | 28.5364 | also red | | | | | | |

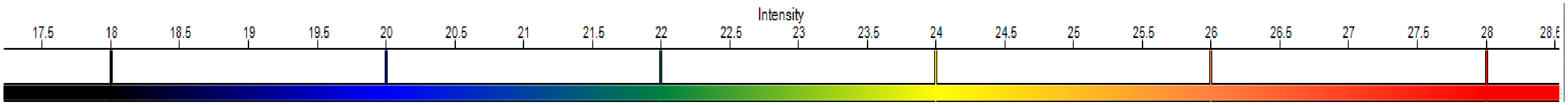

**Table S6: Relative changes in transcript level of several glycosyltransferases in ΔPOMGnT1 cells detected by nCounter analysis.** The transcript level of housekeeping genes ACTB, GAPDH, RAB7A were used for normalization. Assays were performed in duplicate from three biological replicates (n=6) and average ratio (ΔPOMGnT1/WT) of the transcript level of each gene is determined as relative fold change in transcript level in ΔPOMGnT1 cells.

| Gene | Fold change | SD | Log2 fold change |
|---|---|---|---|
| *ALG1* | 0.53 | 0.07 | -0.93 |
| *ALG10* | 1.17 | 0.42 | 0.23 |
| *ALG11* | 0.90 | 0.11 | -0.16 |
| *ALG12* | 0.94 | 0.22 | -0.09 |
| *ALG13* | 0.59 | 0.04 | -0.76 |
| *ALG14* | 0.87 | 0.19 | -0.21 |
| *ALG2* | 0.92 | 0.22 | -0.12 |
| *ALG3* | 0.80 | 0.14 | -0.32 |
| *ALG5* | 0.76 | 0.06 | -0.40 |
| *ALG6* | 1.04 | 0.31 | 0.06 |
| *ALG8* | 0.69 | 0.10 | -0.53 |
| *ALG9* | 0.87 | 0.13 | -0.21 |
| *B3GalNAcT2* | 0.94 | 0.22 | -0.09 |
| *B4GALT1* | 0.73 | 0.11 | -0.45 |
| *DHDDS* | 1.00 | 0.11 | 0.00 |
| *DOLK* | 0.98 | 0.43 | -0.02 |
| *DPAGT1* | 0.86 | 0.09 | -0.22 |
| *DPM1* | 0.96 | 0.10 | -0.06 |
| *DPM2* | 0.85 | 0.11 | -0.24 |
| *DPM3* | 1.23 | 0.45 | 0.29 |
| *DPY19L1* | 0.98 | 0.89 | -0.03 |
| *DPY19L2* | 0.98 | 0.26 | -0.04 |
| *DPY19L3* | 0.67 | 0.22 | -0.58 |
| *DPY19L4* | 0.74 | 0.13 | -0.43 |
| *FUT8* | 0.65 | 0.25 | -0.61 |
| *FUT9* | 1.75 | 1.86 | 0.80 |
| *GALE* | 1.18 | 0.44 | 0.24 |
| *GANAB* | 1.01 | 0.11 | 0.02 |
| *GFPT1* | 1.09 | 0.13 | 0.13 |
| *GMPPA* | 1.05 | 0.12 | 0.07 |
| *GMPPB* | 1.34 | 0.12 | 0.42 |
| *GNE* | 1.38 | 0.94 | 0.46 |
| *MAN1A1* | 0.66 | 0.16 | -0.60 |
| *MAN1B1* | 0.88 | 0.17 | -0.18 |
| *MAN2A1* | 0.63 | 0.11 | -0.68 |
| *MGAT1* | 0.83 | 0.13 | -0.28 |
| *MGAT2* | 0.87 | 0.08 | -0.20 |
| *MGAT3* | 0.65 | 0.61 | -0.63 |
| *MGAT5* | 0.81 | 0.09 | -0.30 |
| *MOGS* | 0.80 | 0.20 | -0.33 |
| *MPDU1* | 1.17 | 0.20 | 0.23 |
| *MPI* | 0.99 | 0.13 | -0.01 |

| | | | |
|---|---|---|---|
| *OGT* | 0.96 | 0.15 | -0.06 |
| *PGM1* | 0.77 | 0.23 | -0.37 |
| *PGM2* | 1.02 | 0.45 | 0.02 |
| *PMM2* | 0.59 | 0.16 | -0.77 |
| *POMGnT1* | 0.41 | 0.34 | -1.28 |
| *POMGnT2* | 0.75 | 0.37 | -0.41 |
| *POMT1* | 1.20 | 0.53 | 0.26 |
| *POMT2* | 1.13 | 0.32 | 0.17 |
| *RFT1* | 0.81 | 0.21 | -0.30 |
| *SLC35A1* | 1.20 | 0.95 | 0.27 |
| *SLC35A2* | 0.92 | 0.17 | -0.12 |
| *SLC35A3* | 0.97 | 0.30 | -0.05 |
| *SLC35C1* | 0.69 | 0.07 | -0.54 |
| *SRD5A3* | 1.06 | 0.09 | 0.09 |
| *ST3GAL3* | 0.31 | 0.13 | -1.68 |
| *ST6GAL1* | 0.17 | 0.20 | -2.54 |
| *UAP1* | 1.10 | 0.07 | 0.13 |

### Supporting Information of N-Cdh *N*- and *O*-man glycoproteomics (SI1- SI4)

#### SI1: Overview of all identified *N*- and *O*-man glycosylation sites of human N-Chd

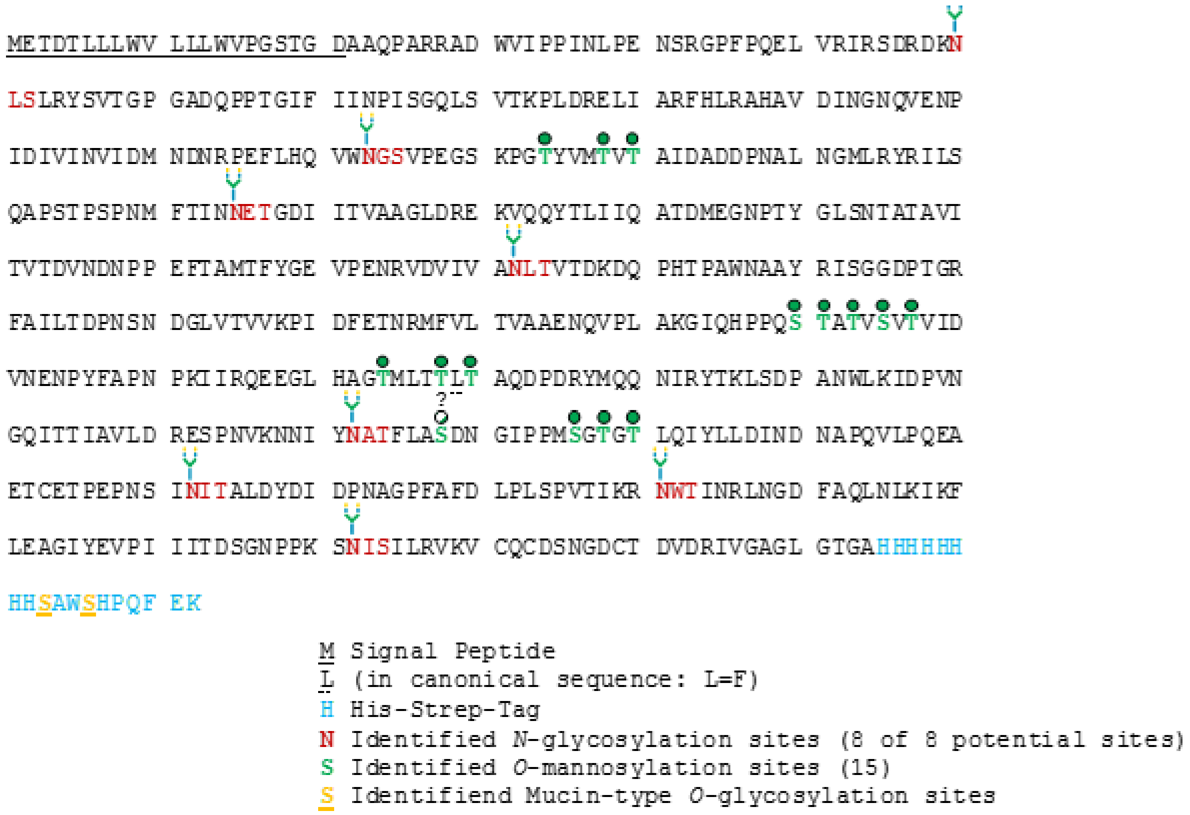

**Figure S6: Summary of all identified *N*-glycan and *O*-mannosylation sites within the protein sequence of human N-Cdh.** Occupied *N*-glycan sites are indicated with a stylized *N*-glycan. Occupied *O*-Man sites are indicated with a green circle (a white circle indicates presumably occupied sites). Please note the protein sequence features a histidine-streptavidin-sequence tag at the C-terminus which was added for the protein purification. Within the strep-tag two mucin-type *O*-glycan sites have been identified.

#### SI2: Site-specific relative changes of ΔPOMTGnT1 N-Cdh *N*-glycan traits in comparison to WT

**Site N190:**

- Site seems to be, if at all, only scarcely occupied
- Only one *N*-Glycopeptide (Pep+HexNAc4Hex5Fuc1NeuAc2) identified (WT, trypsin)

**Site N273:**

**Table S7: Comparative and site-specific *N*-glycoproteomic analysis of quantitative changes in the *N*-glycan microheterogeneity of WT and ΔPOMGnT1-derived N-Cdh for the *N*-glycosylation site N273.** Quantitative changes in the *N*-glycan microheterogeneity are depicted as increase or decrease in ΔPOMGnT1-derived N-Cdh relative to the level in the WT. The quantities represent *N*-glycopeptide area under the curve values of monoisotopic extracted ion chromatograms of the corresponding *N*-glycopeptide precursor ions (EIC MS1) that were summed based on common *N*-glycan traits (*N*-glycopeptides carrying a bisecting GlcNAc, sialylation and/or galactosylation). For each *N*-glycan trait the major *N*-glycan composition with its relative abundance is reported. Abbreviations: Hex (hexose), HexNAc (*N*-acetylhexosamine), Fuc (fucose), NeuAc (*N*-acetylneuraminic acid).

| Site N273 | ΔPOMGnT1 | |
|---|---|---|
| ***N*-glycan traits** | **Sum of relative changes [%]** | **Major *N*-glycan of traits /each trait (± its relative change [%])** |
| Non-Galactosylated | +20.4 | $HexNAc_5Hex_3Fuc_1$ (+10.6 %) |
| Low-Galactosylated | -10.2 | $HexNAc_4Hex_4Fuc_1$ (-11.2 %) |
| Fully Galactosylated | -10.3 | $HexNAc_4Hex_5Fuc_1NeuAc_1$ (-12.2 %) |
| Non-Sialylated | +9.1 | $HexNAc_4Hex_4Fuc_1$ (-11.2 %) |
| Mono-Sialylated | -9.1 | $HexNAc_4Hex_5Fuc_1NeuAc_1$ (-12.2 %) |
| No Bisecting GlcNAc | -15.0 | $HexNAc_4Hex_5Fuc_1NeuAc_1$ (-12.2 %) |
| With Bisecting GlcNAc | +15.0 | $HexNAc_5Hex_3Fuc_1$ (+10.6 %) |

- **Galactosylation:**
  - ΔPOMGnT1: degree of galactosylation goes down significantly, resulting in an increase of non-galactosylated *N*-glycopeptides
- **Sialylation:**
  - ΔPOMGnT1: number of mono-sialylated *N*-glycopeptides goes down significantly, resulting in an increase of non-sialylated *N*-glycopeptides (no di-sialylated *N*-glycopeptides detected)
- **Bisecting GlcNAc:**
  - ΔPOMGnT1: number of *N*-glycopeptides with bisecting GlcNAc increases significantly

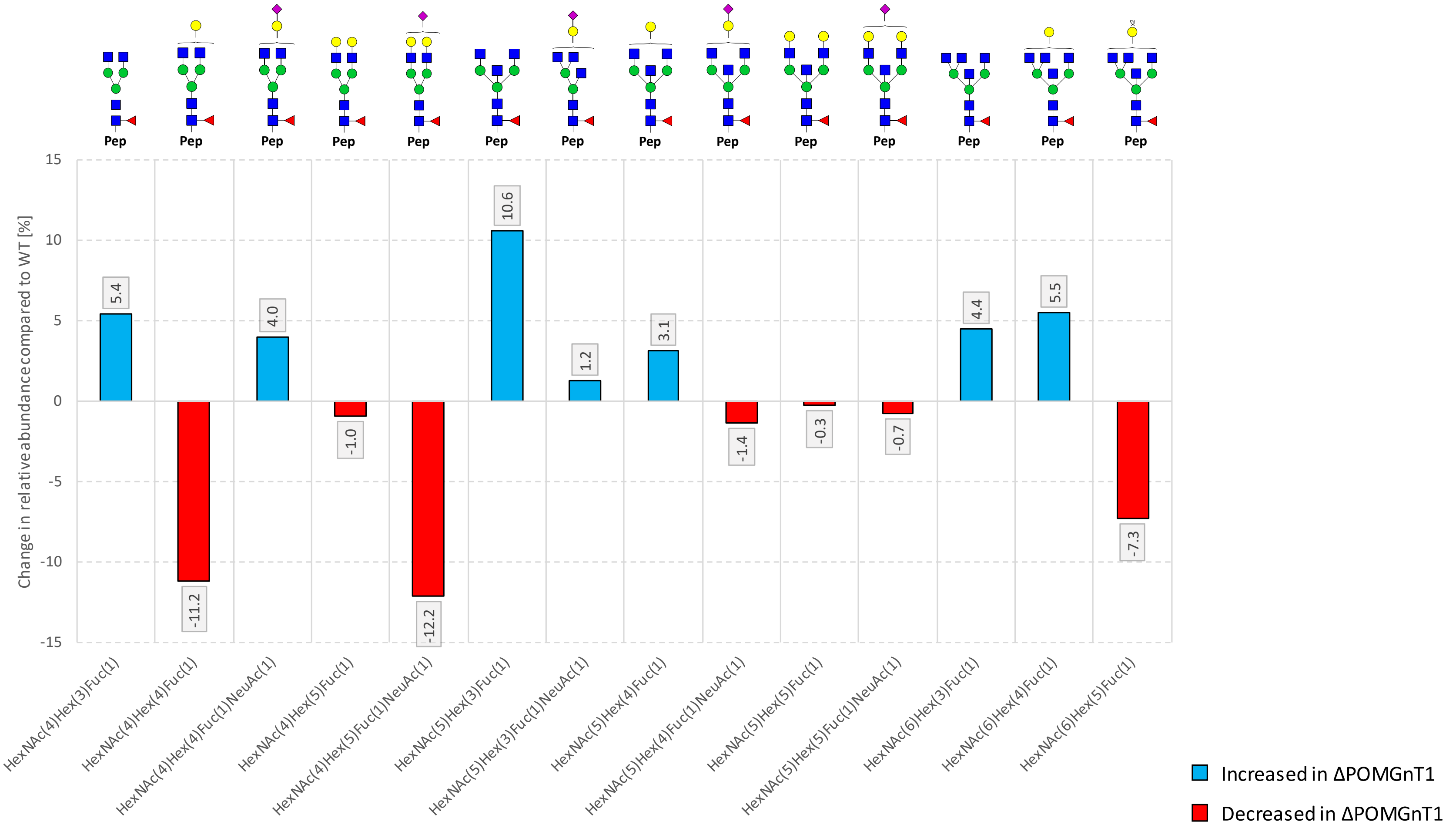

**Figure S7: Site N273-specific relative changes in abundance of *N*-glycan compositions detected in ΔPOMGnT1-derived N-Cdh compared to the WT-derived N-Cdh.** *N*-glycan structures were drawn with GlycoWorkbench Version 1.1, by following the guideline of the Consortium for Functional Glycomics. Abbreviations: Hex (hexose), HexNAc (*N*-acetylhexosamine), Fuc (fucose), NeuAc (*N*-acetylneuraminic acid), blue square (*N*-acetylglucosamine), green circle (mannose), yellow circle (galactose), pink diamond (*N*-acetylneuraminic acid), red triangle (fucose), Pep (peptide).

**Site 325:**

**Table S8: Comparative and site-specific *N*-glycoproteomic analysis of quantitative changes in the *N*-glycan microheterogeneity of WT and ΔPOMGnT1-derived N-Cdh for the *N*-glycosylation site N325.** Quantitative changes in the *N*-glycan microheterogeneity are depicted as increase or decrease in ΔPOMGnT1-derived N-Cdh relative to the level in the WT. The quantities represent *N*-glycopeptide area under the curve values of monoisotopic extracted ion chromatograms of the corresponding *N*-glycopeptide precursor ions (EIC MS1) that were summed based on common *N*-glycan traits (*N*-glycopeptides carrying a bisecting GlcNAc, sialylation and/or galactosylation). For each *N*-glycan trait the major *N*-glycan composition with its relative abundance is reported. Abbreviations: Hex (hexose), HexNAc (*N*-acetylhexosamine), Fuc (fucose), NeuAc (*N*-acetylneuraminic acid).

| Site N325 | ΔPOMGnT1 | |
|---|---|---|
| ***N*-glycan traits** | **Sum of relative changes [%]** | **Major *N*-glycan of traits /each trait (± its relative change [%])** |
| Non-Galactosylated | +28.2 | $HexNAc_5Hex_3Fuc_1$ (+19.8 %) |
| Low-Galactosylated | -6.1 | $HexNAc_5Hex_4Fuc_1NeuAc_1$ (-9.8 %) |
| Fully Galactosylated | -22.2 | $HexNAc_5Hex_5Fuc_1$ (-6.3 %) |
| Non-Sialylated | +22.7 | $HexNAc_5Hex_5Fuc_1$ (+19.8 %) |
| Mono-Sialylated | -17.8 | $HexNAc_5Hex_4Fuc_1NeuAc_1$ (-9.8 %) |
| Di-Sialylated | -4.8 | $HexNAc_4Hex_5Fuc_1NeuAc_2$ (-3.6 %) |
| No Bisecting GlcNAc | +9.1 | $HexNAc_4Hex_5Fuc_1$ (+6.7 %) |
| With Bisecting GlcNAc | -9.1 | $HexNAc_5Hex_3Fuc_1$ (+19.8 %) |

- **Galactosylation:**
  - ΔPOMGnT1: degree of galactosylation goes down significantly, resulting in an increase of non-galactosylated *N*-glycopeptides
- **Sialylation:**
  - ΔPOMGnT1: number of mono- and di-sialylated *N*-glycopeptides goes down significantly, resulting in an increase of non-sialylated *N*-glycopeptides
- **Bisecting GlcNAc:**
  - ΔPOMGnT1: number of *N*-glycopeptides with bisecting GlcNAc decreases significantly

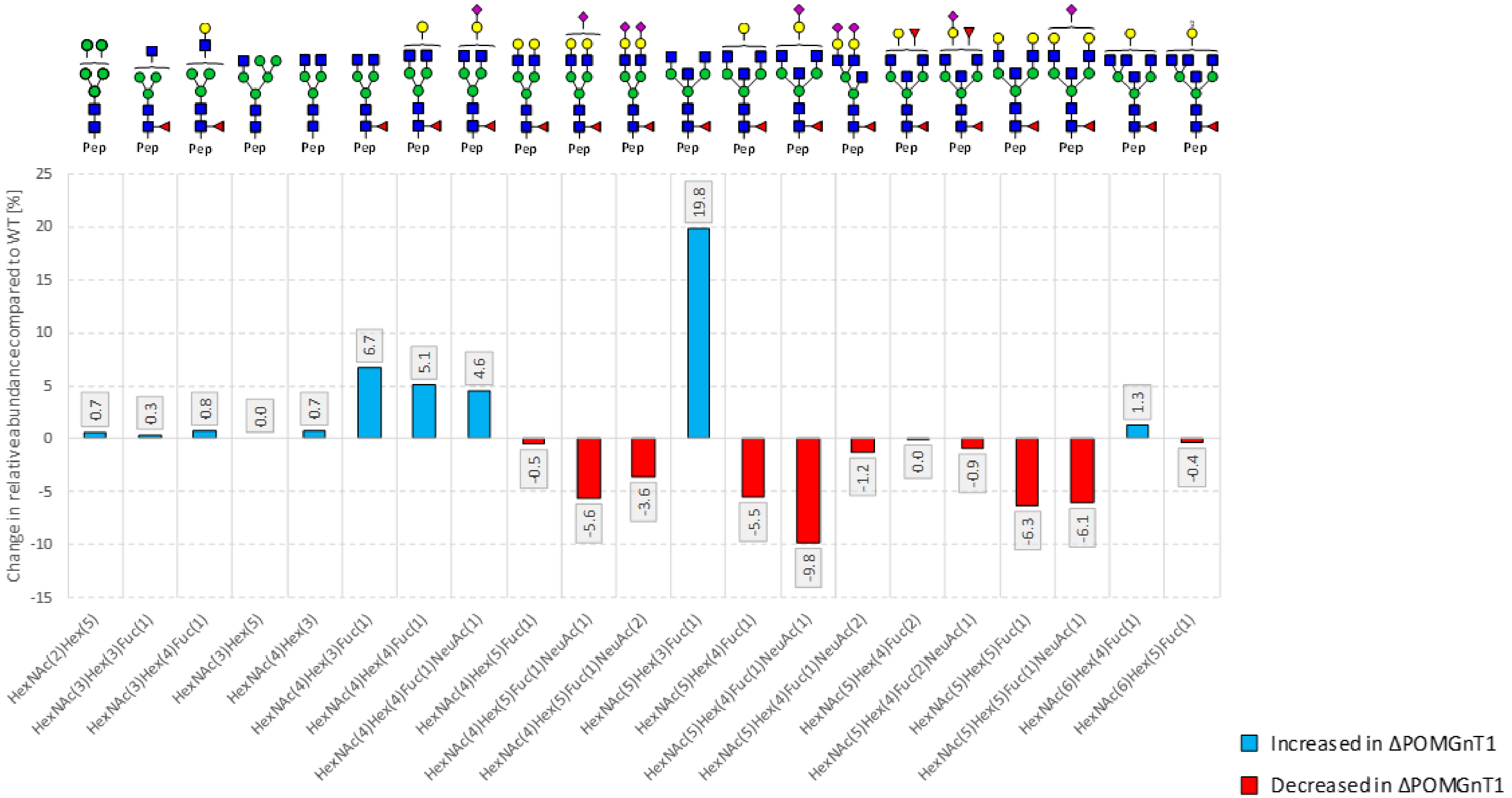

**Figure S8: Site N325-specific relative changes in abundance of *N*-glycan compositions detected in ΔPOMGnT1-derived N-Cdh compared to the WT-derived N-Cdh.** N-glycan structures were drawn with GlycoWorkbench Version 1.1, by following the guideline of the Consortium for Functional Glycomics. Abbreviations: Hex (hexose), HexNAc (*N*-acetylhexosamine), Fuc (fucose), NeuAc (*N*-acetylneuraminic acid), blue square (*N*-acetylglucosamine), green circle (mannose), yellow circle (galactose), pink diamond (*N*-acetylneuraminic acid), red triangle (fucose), Pep (peptide).

**Site N402:**

**Table S9: Comparative and site-specific *N*-glycoproteomic analysis of quantitative changes in the *N*-glycan microheterogeneity of WT and ΔPOMGnT1-derived N-Cdh for the *N*-glycosylation site N402.** Quantitative changes in the *N*-glycan microheterogeneity are depicted as increase or decrease in ΔPOMGnT1-derived N-Cdh relative to the level in the WT. The quantities represent *N*-glycopeptide area under the curve values of monoisotopic extracted ion chromatograms of the corresponding *N*-glycopeptide precursor ions (EIC MS1) that were summed based on common *N*-glycan traits (*N*-glycopeptides carrying a bisecting GlcNAc, sialylation and/or galactosylation). For each *N*-glycan trait the major *N*-glycan composition with its relative abundance is reported. Abbreviations: Hex (hexose), HexNAc (*N*-acetylhexosamine), Fuc (fucose), NeuAc (*N*-acetylneuraminic acid).

| Site N402 | ΔPOMGnT1 | |
|---|---|---|
| ***N*-glycan traits** | **Sum of relative changes [%]** | **Major *N*-glycan of traits /each trait (± its relative change [%])** |
| Non-Galactosylated | +12.6 | $HexNAc_5Hex_3Fuc_1$ (+8.7 %) |
| Low-Galactosylated | +16.4 | $HexNAc_6Hex_4Fuc_1$ (7.1 %) |
| Fully Galactosylated | -29.0 | $HexNAc_5Hex_5Fuc_1NeuAc_2$ (-8.3 %) |
| Non-Sialylated | +15.8 | $HexNAc_5Hex_3Fuc_1$ (+8.7 %) |
| Mono-Sialylated | -10.8 | $HexNAc_5Hex_5Fuc_1NeuAc_1$ (-5.5 %) |
| Di-Sialylated | -5.0 | $HexNAc_4Hex_5Fuc_1NeuAc_2$ (-8.3 %) |
| No Bisecting GlcNAc | -14.0 | $HexNAc_4Hex_5Fuc_1NeuAc_2$ (-8.3 %) |
| With Bisecting GlcNAc | +14.0 | $HexNAc_5Hex_3Fuc_1$ (+8.7 %) |

- **Galactosylation:**
  - ΔPOMGnT1: number of fully galactosylated *N*-glycopeptides goes down significantly, resulting in an increase of low- and non-galactosylated *N*-glycopeptides
- **Sialylation:**
  - ΔPOMGnT1: number of mono- and di-sialylated *N*-glycopeptides goes down, resulting in an increase of non-sialylated *N*-glycopeptides
- **Bisecting GlcNAc:**
  - ΔPOMGnT1: number of *N*-glycopeptides with bisecting GlcNAc increases significantly

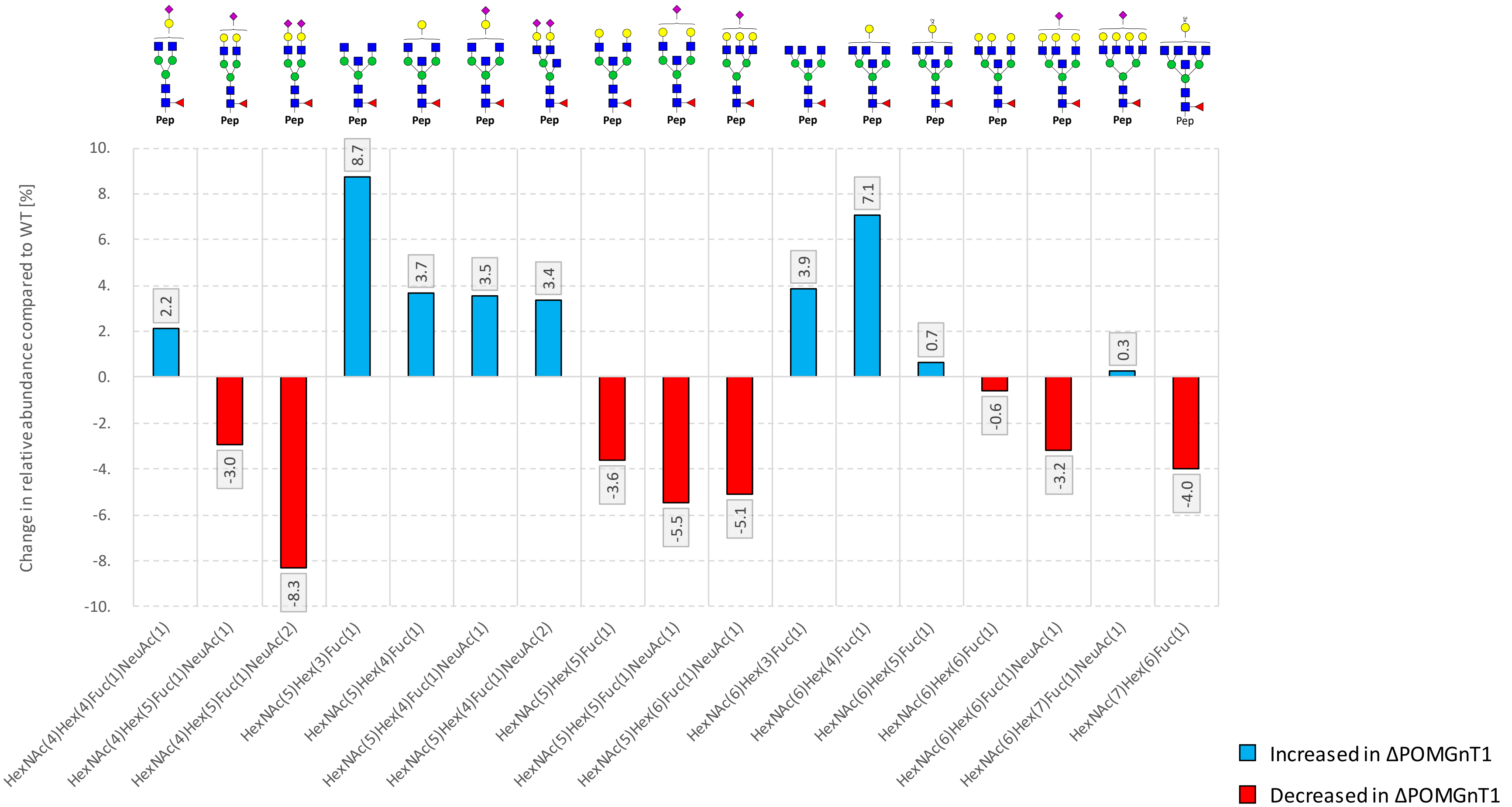

**Figure S9: Site N402-specific relative changes in abundance of *N*-glycan compositions detected in ΔPOMGnT1-derived N-Cdh compared to the WT-derived N-Cdh.** *N*-glycan structures were drawn with GlycoWorkbench Version 1.1, by following the guideline of the Consortium for Functional Glycomics. Abbreviations: Hex (hexose), HexNAc (*N*-acetylhexosamine), Fuc (fucose), NeuAc (*N*-acetylneuraminic acid), blue square (*N*-acetylglucosamine), green circle (mannose), yellow circle (galactose), pink diamond (*N*-acetylneuraminic acid), red triangle (fucose), Pep (peptide).

**Site N572:**

**Table S10: Comparative and site-specific *N*-glycoproteomic analysis of quantitative changes in the *N*-glycan microheterogeneity of WT and ΔPOMGnT1-derived N-Cdh for the *N*-glycosylation site N572**. Quantitative changes in the *N*-glycan microheterogeneity are depicted as increase or decrease in ΔPOMGnT1-derived N-Cdh relative to the level in the WT. The quantities represent *N*-glycopeptide area under the curve values of monoisotopic extracted ion chromatograms of the corresponding *N*-glycopeptide precursor ions (EIC MS1) that were summed based on common *N*-glycan traits (*N*-glycopeptides carrying a bisecting GlcNAc, sialylation and/or galactosylation). For each *N*-glycan trait the major N-glycan composition with its relative abundance is reported. Abbreviations: Hex (hexose), HexNAc (*N*-acetylhexosamine), Fuc (fucose), NeuAc (*N*-acetylneuraminic acid).

| Site N572 | ΔPOMGnT1 | |
|---|---|---|
| ***N*-glycan traits** | **Sum of relative changes [%]** | **Major *N*-glycan of traits /each trait (± its relative change [%])** |
| Non-Galactosylated | +11.8 | $HexNAc_5Hex_3$ (+19.8 %) |
| Low-Galactosylated | -1.7 | $HexNAc_4Hex_4$ (+11.6 %) |
| Fully Galactosylated | -10.1 | $HexNAc_4Hex_5$ (-7.6 %) |
| Non-Sialylated | -0.6 | $HexNAc_5Hex_3$ (+19.8 %) |
| Mono-Sialylated | +0.6 | $HexNAc_4Hex_4NeuAc_1$ (+2.2 %) |
| No Bisecting GlcNAc | -6.9 | $HexNAc_4Hex_4$ (+11.6 %) |
| With Bisecting GlcNAc | +6.9 | $HexNAc_5Hex_3$ (+19.8 %) |

- **Galactosylation:**
  - ΔPOMGnT1: number of fully galactosylated *N*-glycopeptides goes down significantly, resulting in an increase of non-galactosylated *N*-glycopeptides. Number of low-galactosylated *N*-glycopeptides does not change significantly
- **Sialylation:**
  - ΔPOMGnT1: no significant changes with regard to sialylation
- **Bisecting GlcNAc:**
  - ΔPOMGnT1: number of *N*-glycopeptides with bisecting GlcNAc increases significantly

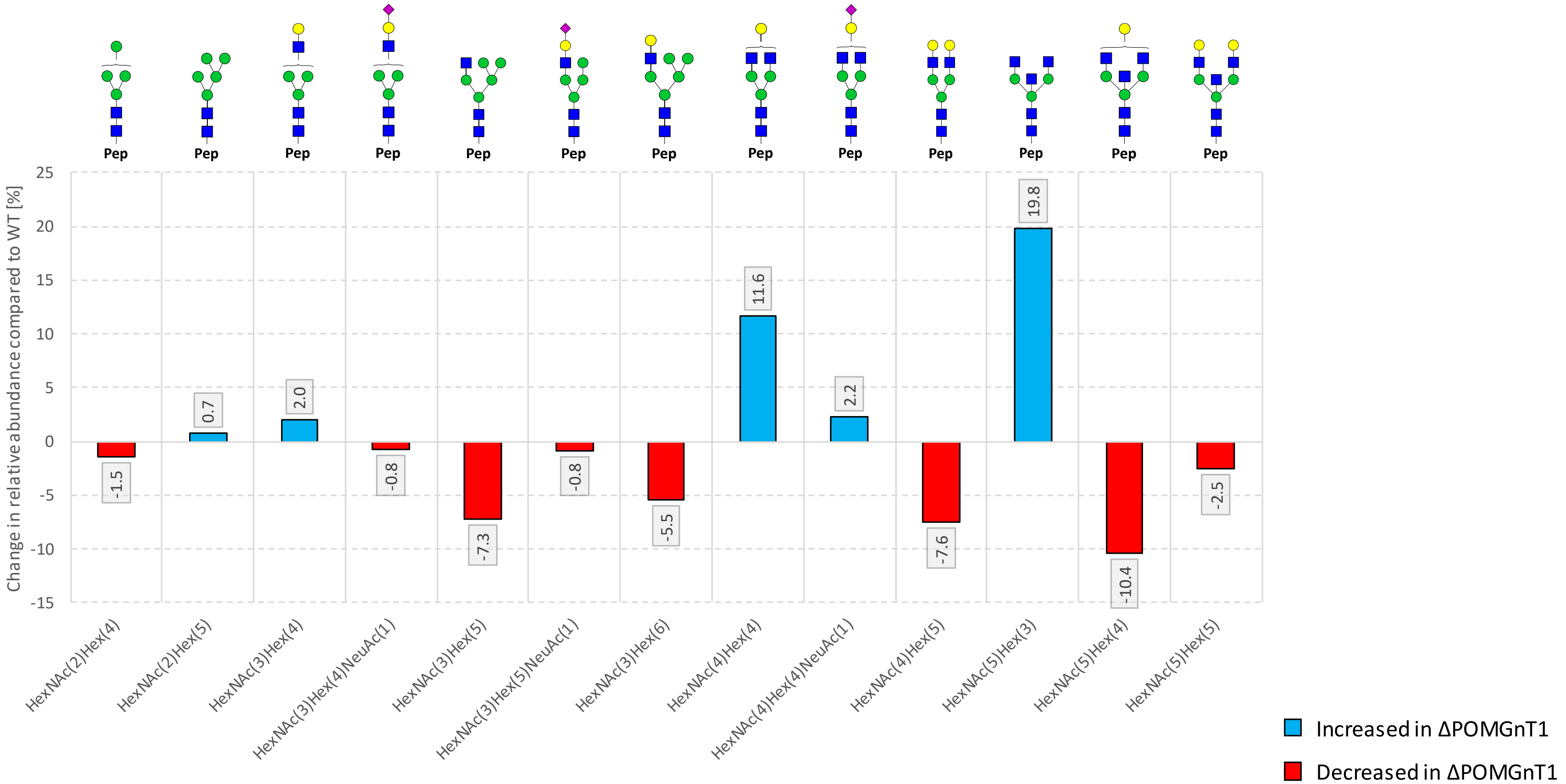

**Figure S10: Site N572-specific relative changes in abundance of *N*-glycan compositions detected in ΔPOMGnT1-derived N-Cdh compared to the WT-derived N-Cdh.** *N*-glycan structures were drawn with GlycoWorkbench Version 1.1, by following the guideline of the Consortium for Functional Glycomics. Abbreviations: Hex (hexose), HexNAc (*N*-acetylhexosamine), Fuc (fucose), NeuAc (*N*-acetylneuraminic acid), blue square (*N*-acetylglucosamine), green circle (mannose), yellow circle (galactose), pink diamond (*N*-acetylneuraminic acid), red triangle (fucose), Pep (peptide).

**Site N622:**

**Table S11: Comparative and site-specific *N*-glycoproteomic analysis of quantitative changes in the *N*-glycan microheterogeneity of WT and ΔPOMGnT1-derived N-Cdh for the *N*-glycosylation site N622.** Quantitative changes in the N-glycan microheterogeneity are depicted as increase or decrease in ΔPOMGnT1-derived N-Cdh relative to the level in the WT. The quantities represent *N*-glycopeptide area under the curve values of monoisotopic extracted ion chromatograms of the corresponding *N*-glycopeptide precursor ions (EIC MS1) that were summed based on common N-glycan traits (*N*-glycopeptides carrying a bisecting GlcNAc, sialylation and/or galactosylation). For each *N*-glycan trait the major N-glycan composition with its relative abundance is reported. Abbreviations: Hex (hexose), HexNAc (*N*-acetylhexosamine), Fuc (fucose), NeuAc (*N*-acetylneuraminic acid).

| N622 | ΔPOMGnT1 | |
|---|---|---|
| ***N*-glycan traits** | **Sum of relative changes [%]** | **Major *N*-glycan of traits /each trait (± its relative change [%])** |
| Non-Galactosylated | +18.6 | $HexNAc_5Hex_3Fuc_1$ (+9.6 %) |
| Low-Galactosylated | +18.2 | $HexNAc_6Hex_5Fuc_1NeuAc_2$ (+6.1 %) |
| Fully Galactosylated | -36.8 | $HexNAc_6Hex_6Fuc_1NeuAc_2$ (-11.8 %) |
| Non-Sialylated | +16.6 | $HexNAc_5Hex_3Fuc_1$ (+9.6 %) |
| Mono-Sialylated | -6.6 | $HexNAc_6Hex_6Fuc_1NeuAc_1$ (-3.7 %) |
| Di-Sialylated | -10.0 | $HexNAc_6Hex_6Fuc_1NeuAc_2$ (-11.8 %) |
| No Bisecting GlcNAc | -2.0 | $HexNAc_4Hex_3Fuc_1$ (+4.4 %) |
| With Bisecting GlcNAc | +2.0 | $HexNAc_6Hex_6Fuc_1NeuAc_2$ (-11.8 %) |

- **Galactosylation:**
    - ΔPOMGnT1: number of fully galactosylated *N*-glycopeptides goes down significantly, resulting in an increase of low and non-galactosylated *N*-glycopeptides.
- **Sialylation:**
    - ΔPOMGnT1: number of non-sialylated *N*-glycopeptides increases. Number of mono- and di-sialylated *N*-glycopeptides decreases.
- **Bisecting GlcNAc:**
    - ΔPOMGnT1: only minor changes with regard to bisecting GlcNAc

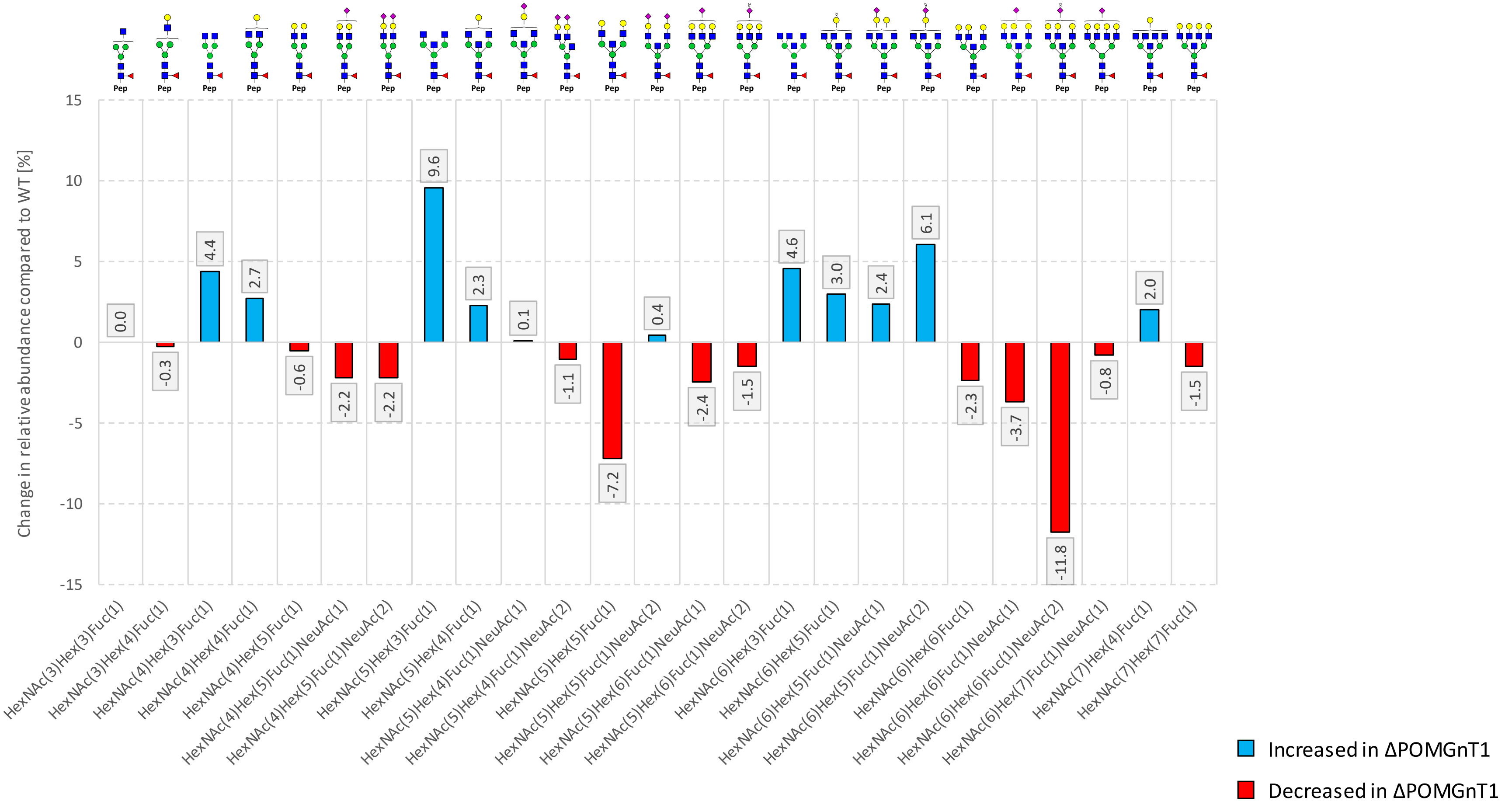

**Figure S11: Site N622-specific relative changes in abundance of *N*-glycan compositions detected in ΔPOMGnT1-derived N-Cdh compared to the WT-derived N-Cdh.** *N*-glycan structures were drawn with GlycoWorkbench Version 1.1, by following the guideline of the Consortium for Functional Glycomics. Abbreviations: Hex (hexose), HexNAc (*N*-acetylhexosamine), Fuc (fucose), NeuAc (*N*-acetylneuraminic acid), blue square (*N*-acetylglucosamine), green circle (mannose), yellow circle (galactose), pink diamond (*N*-acetylneuraminic acid), red triangle (fucose), Pep (peptide).

**Site N651:**

**Table S12: Comparative and site-specific *N*-glycoproteomic analysis of quantitative changes in the *N*-glycan microheterogeneity of WT and ΔPOMGnT1-derived N-Cdh for the *N*-glycosylation site N651.** Quantitative changes in the *N*-glycan microheterogeneity are depicted as increase or decrease in ΔPOMGnT1-derived N-Cdh relative to the level in the WT. The quantities represent *N*-glycopeptide area under the curve values of monoisotopic extracted ion chromatograms of the corresponding *N*-glycopeptide precursor ions (EIC MS1) that were summed based on common *N*-glycan traits (*N*-glycopeptides carrying a bisecting GlcNAc, sialylation and/or galactosylation). For each *N*-glycan trait the major *N*-glycan composition with its relative abundance is reported. Abbreviations: Hex (hexose), HexNAc (*N*-acetylhexosamine), Fuc (fucose), NeuAc (*N*-acetylneuraminic acid).

| Site N651 | ΔPOMGnT1 | |
|---|---|---|
| ***N*-glycan traits** | **Sum of relative changes [%]** | **Major *N*-glycan of traits /each trait (± its relative change [%])** |
| Non-Galactosylated | +21.3 | $HexNAc_2Hex_7$ (+9.4 %) |
| Low-Galactosylated | -19.7 | $HexNAc_3Hex_6NeuAc_1$ (-11.9 %) |
| Fully Galactosylated | -1.6 | $HexNAc_4Hex_5$ (-0.6 %) |
| Non-Sialylated | +19.6 | $HexNAc_2Hex_7$ (+9.4 %) |
| Mono-Sialylated | -19.6 | $HexNAc_3Hex_6NeuAc_1$ (-11.9 %) |
| No Bisecting GlcNAc | +3.0 | $HexNAc_3Hex_6NeuAc_1$ (-11.9 %) |
| With Bisecting GlcNAc | -3.0 | $HexNAc_5Hex_4NeuAc_1$ (-2.2 %) |

- **High-mannose-type *N*-glycans:** site N651 mainly features high-mannose-type *N*-glycans
- **Galactosylation:**
  - ΔPOMGnT1: number of low-galactosylated *N*-glycopeptides goes down significantly, resulting in an increase of non-galactosylated *N*-glycopeptides. The number of fully galactosylated *N*-glycopeptides does not change significantly.
- **Sialylation:**
  - ΔPOMGnT1: number of mono-sialylated *N*-glycopeptides decreases significantly. As a result, the number of non-sialylated *N*-glycopeptides increases accordingly.
- **Bisecting GlcNAc:**
  - ΔPOMGnT1: no significant changes with regard to bisecting GlcNAc.

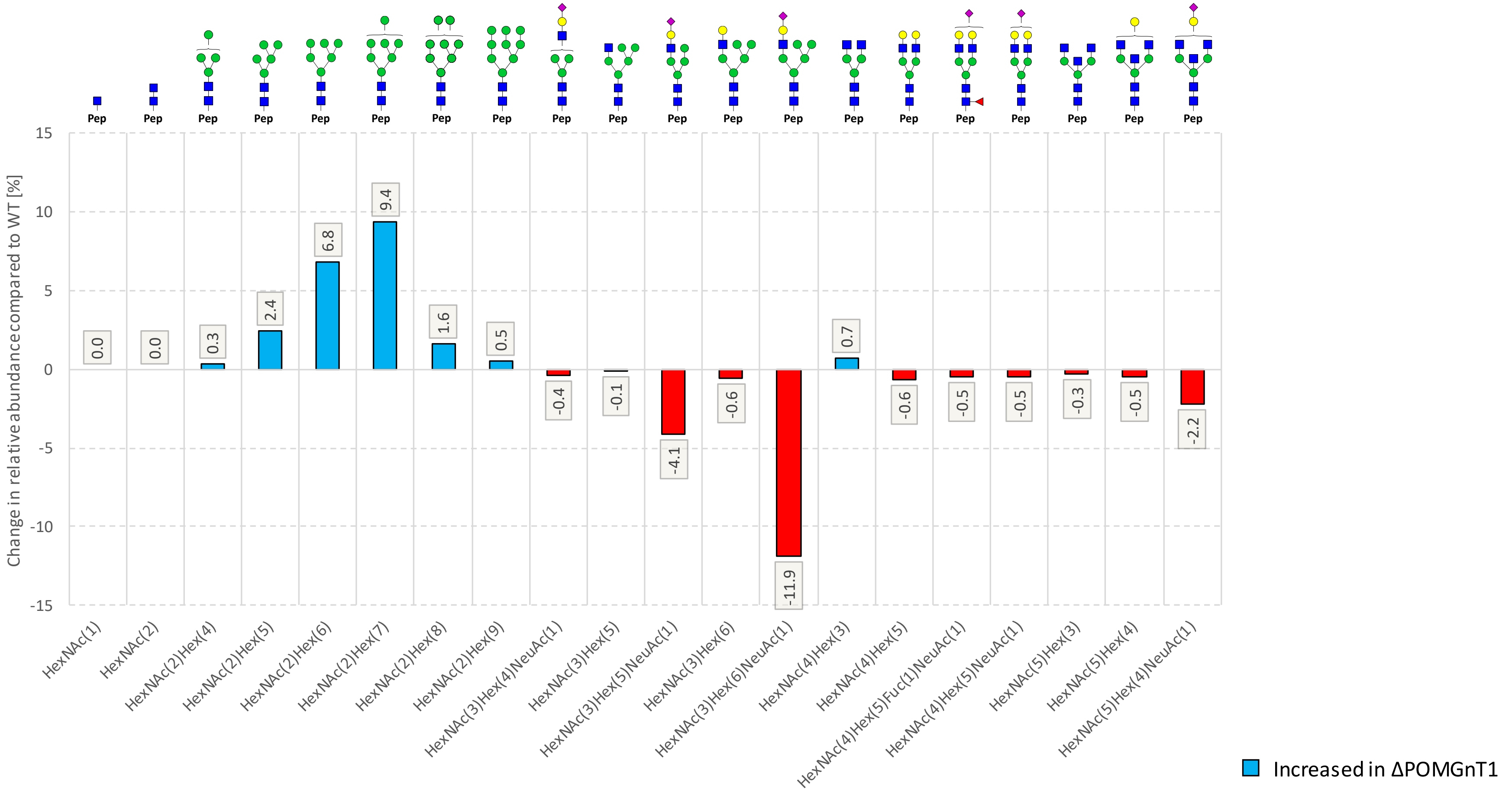

**Figure S12: Site N651-specific relative changes in abundance of *N*-glycan compositions detected in ΔPOMGnT1-derived N-Cdh compared to the WT-derived N-Cdh.** *N*-glycan structures were drawn with GlycoWorkbench Version 1.1, by following the guideline of the Consortium for Functional Glycomics. Abbreviations: Hex (hexose), HexNAc (*N*-acetylhexosamine), Fuc (fucose), NeuAc (*N*-acetylneuraminic acid), blue square (*N*-acetylglucosamine), green circle (mannose), yellow circle (galactose), pink diamond (N-acetylneuraminic acid), red triangle (fucose), Pep (peptide).

**<u>Site N692</u>**

**Table S13: Comparative and site-specific *N*-glycoproteomic analysis of quantitative changes in the *N*-glycan microheterogeneity of WT and ΔPOMGnT1-derived N-Cdh for the *N*-glycosylation site N692.** Quantitative changes in the *N*-glycan microheterogeneity are depicted as increase or decrease in ΔPOMGnT1-derived N-Cdh relative to the level in the WT. The quantities represent *N*-glycopeptide area under the curve values of monoisotopic extracted ion chromatograms of the corresponding N-glycopeptide precursor ions (EIC MS1) that were summed based on common *N*-glycan traits (*N*-glycopeptides carrying a bisecting GlcNAc, sialylation and/or galactosylation). For each *N*-glycan trait the major *N*-glycan composition with its relative abundance is reported. Abbreviations: Hex (hexose), HexNAc (N-acetylhexosamine), Fuc (fucose), NeuAc (N-acetylneuraminic acid).

| Site N692 | ΔPOMGnT1 | |
|---|---|---|
| ***N*-glycan traits** | **Sum of relative changes [%]** | **Major *N*-glycan of traits /each trait (± its relative change [%])** |
| Non-Galactosylated | +22.5 | $HexNAc_2Hex_2$ (+6.3 %) |
| Low-Galactosylated | +24.6 | $HexNAc_5Hex_4Fuc_1$ (+8.8 %) |
| Fully Galactosylated | -47.1 | $HexNAc_4Hex_5Fuc_1NeuAc_1$ (-21.2 %) |
| Non-Sialylated | +29.8 | $HexNAc_5Hex_4Fuc_1$ (+8.8 %) |
| Mono-Sialylated | -25.6 | $HexNAc_4Hex_5Fuc_1NeuAc_1$ (-21.2 %) |
| Di-Sialylated | -4.3 | $HexNAc_4Hex_5Fuc_1NeuAc_2$ (-4.3 %) |
| No Bisecting GlcNAc | -3.0 | $HexNAc_4Hex_5Fuc_1NeuAc_1$ (-21.2 %) |
| With Bisecting GlcNAc | +3.0 | $HexNAc_5Hex_4Fuc_1$ (+8.8 %) |

- **Galactosylation:**
  - ΔPOMGnT1: number of fully galactosylated *N*-glycopeptides goes down significantly, resulting in an increase of low and non-galactosylated *N*-glycopeptides.
  - 
- **Sialylation:**
  - ΔPOMGnT1: number of non-sialylated *N*-glycopeptides increases significantly. In return the number of mono-sialylated *N*-glycopeptides decreases. The number of di-sialylated *N*-glycopeptides does not change significantly.
- **Bisecting GlcNAc:**
  - ΔPOMGnT1: number of *N*-glycopeptides with bisecting GlcNAc does not change significantly.

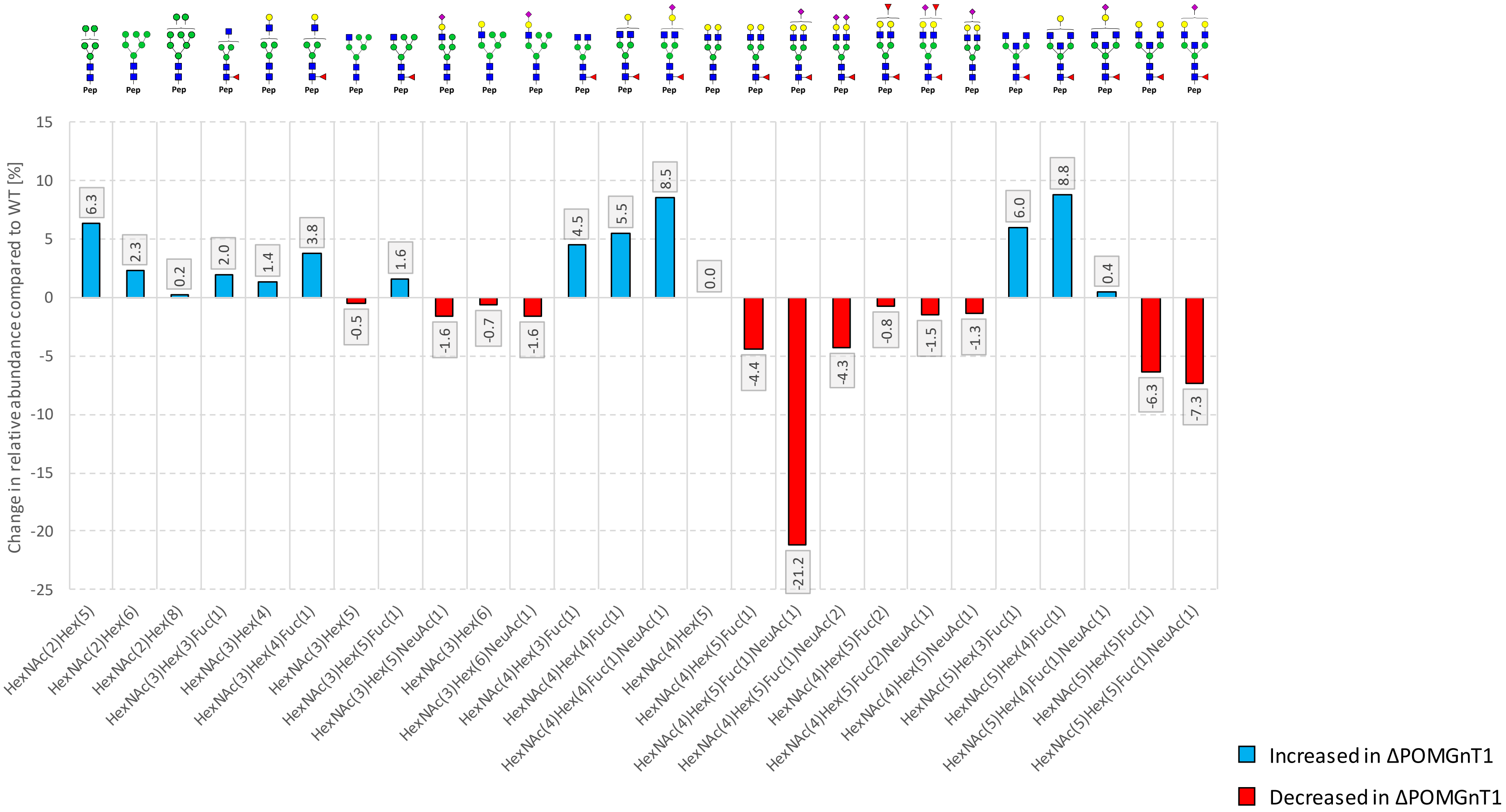

**Figure S13: Site N692-specific relative changes in abundance of *N*-glycan compositions detected in ΔPOMGnT1-derived N-Cdh compared to the WT-derived N-Cdh.** *N*-glycan structures were drawn with GlycoWorkbench Version 1.1, by following the guideline of the Consortium for Functional Glycomics. Abbreviations: Hex (hexose), HexNAc (*N*-acetylhexosamine), Fuc (fucose), NeuAc (*N*-acetylneuraminic acid), blue square (*N*-acetylglucosamine), green circle (mannose), yellow circle (galactose), pink diamond (*N*-acetylneuraminic acid), red triangle (fucose), Pep (peptide).

#### SI3: Number of identified N-Cdh *N*-glycopeptides

**Table S14: Number of identified and proteinase K-generated N-Cdh *N*-glycopeptides per *N*-glycosylation site when using HCD. low fragmentation.**

| Prot K \| HCD.low | | |
|---|---|---|
| ***N*-Glycosylation Site**<br>Peptide Sequence | **ΔPOMGnT1** | **WT** |
| **N273** | **26** | **20** |
| NGSVPEGSKPG | 1 | 1 |
| VWNG | 8 | 7 |
| VWNGSVPE | 3 | 3 |
| VWNGSVPEGSKPG | 14 | 9 |
| **N325** | **72** | **61** |
| INNETGDII | 13 | 9 |
| INNETGDIIT | 13 | 8 |
| NETGDII | 28 | 24 |
| NETGDIIT | 2 | 6 |
| TINNETGDII | 12 | 11 |
| TINNETGDIIT | 4 | 3 |
| **N402** | **17** | **20** |
| ANLT | 10 | 11 |
| NLTVT | 7 | 9 |
| **N572** | **11** | **15** |
| IYNAT | 11 | 15 |
| **N622** | **24** | **28** |
| SINIT | 24 | 28 |
| **N651** | **6** | **12** |
| NWTINR | 1 | 4 |
| RNW | 2 | 2 |
| RNWTIN | 3 | 6 |
| **N692** | **10** | **4** |
| DSGNPPKSNIS | 10 | 4 |

**Table S15: Number of identified and proteinase K-generated N-Cdh *N*-glycopeptides per *N*-glycosylation site when using HCD. step fragmentation.**

| Prot K \| HCD.step | | |
|---|---|---|
| ***N*-Glycosylation Site**<br>Peptide Sequence | **ΔPOMGnT1** | **WT** |
| **N273** | **23** | **15** |
| NGSVPEGSKPG | 1 | 1 |
| NGSVPEGSKPGTY | 3 | 1 |
| VWNG | 7 | 4 |
| VWNGSVPE | - | 1 |
| VWNGSVPEGSKPG | 12 | 7 |
| VWNGSVPEGSKPGTY | | 1 |
| **N325** | **48** | **47** |
| INNETGDII | 7 | 8 |
| INNETGDIIT | 1 | 2 |
| NETGDII | 23 | 18 |
| NETGDIIT | 3 | 6 |
| TINNETGDII | 13 | 11 |
| TINNETGDIIT | 1 | 2 |
| **N402** | **6** | **6** |
| ANLT | 2 | 6 |
| NLTVT | 4 | - |
| **N572** | **5** | **8** |
| IYNAT | 5 | 8 |
| **N622** | **22** | **26** |
| SINIT | 22 | 26 |
| **N651** | **2** | **5** |
| RNWTIN | 2 | 5 |
| **N692** | **12** | **11** |
| DSGNPPKSNIS | 12 | 11 |

**Table S16: Number of identified tryptic N-Cdh *N*-glycopeptides per *N*-glycosylation site when using HCD. low fragmentation.**

| Trypsin \| HCD.low | | |
|---|---|---|
| **N-Glycosylation Site**<br>Peptide Sequence | **ΔPOMGnT1** | **WT** |
| **N190** | **-** | **1** |
| DKNLSLR | - | 1 |
| **N325** | **1** | **-** |
| ILSQAPSTPSPNMFTINNETGDIITVAAGLDR | 1 | - |
| **N651** | **26** | **26** |
| NWTINR | 22 | 21 |
| RNWTINR | 4 | 5 |
| **N692** | **37** | **34** |
| SNISILR | 37 | 34 |

**Table S17: Number of identified tryptic N-Cdh *N*-glycopeptides per *N*-glycosylation site when using HCD. step fragmentation.**

| Trypsin \| HCD.step | | |
|---|---|---|
| ***N*-Glycosylation Site**<br>Peptide Sequence | **ΔPOMGnT1** | **WT** |
| **N190** | **-** | **1** |
| DKNLSLR | - | 1 |
| **N325** | **1** | **-** |
| ILSQAPSTPSPNMFTINNETGDIITVAAGLDR | 1 | - |
| **N402** | **1** | **-** |
| VDVIVANLTVTDKDQPHTPAWNAAYR | 1 | - |
| **N651** | **17** | **22** |
| NWTINR | 13 | 19 |
| RNWTINR | 4 | 3 |
| **N692** | **26** | **22** |
| SNISILR | 26 | 22 |

#### SI4: Overview of *O*-Man Glycoproteomics of N-Cdh:

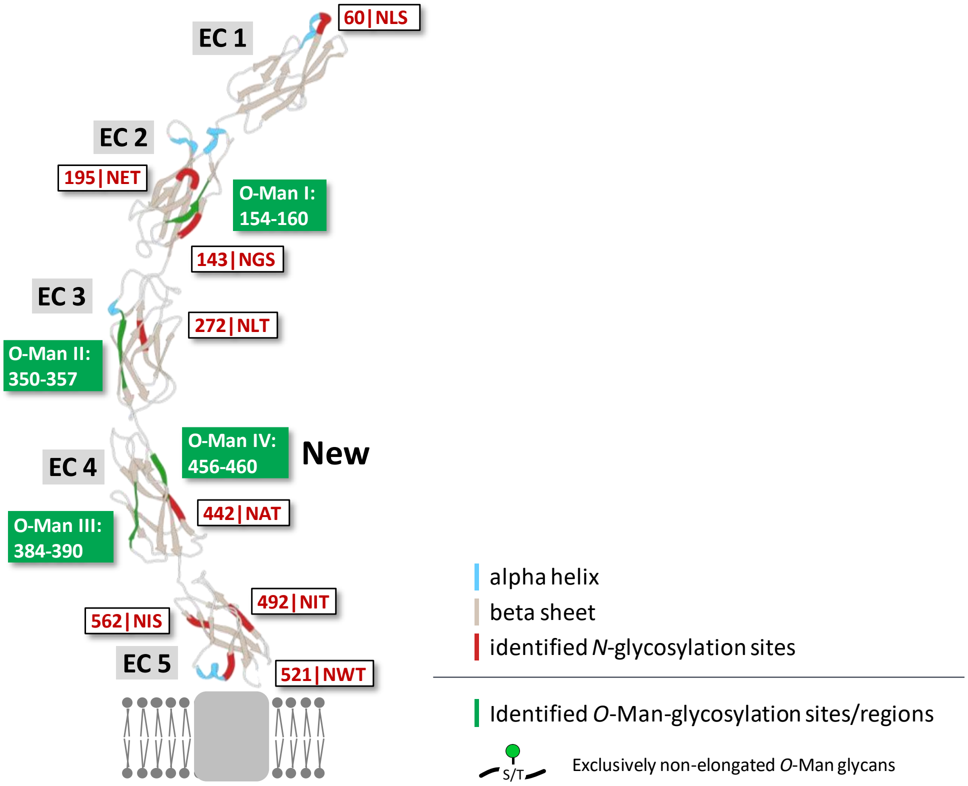

**Figure S14: Site-specific *O*-mannosylation glycoproteomic analysis of extracellular (EC) domains of human N-Cdh, purified from WT and ΔPOMGnT1 HEK293T cells.** A total of 4 N-Cdh *O*-mannosylation regions (I-IV) with their respective O-Man glycoforms (exclusively non-elongated *O*-Man glycans) were identified. To achieve this, hydrophilic interaction liquid chromatography (HILIC)-enriched tryptic and proteinase K-generated N-Cdh *O*-Man glycopeptides were analyzed by nano-reversed-phase liquid chromatography coupled online to an electrospray ionization tandem mass spectrometer (nano-RP-LC-ESI-OT-OT-MS/MS). All *O*-Man regions are located within beta sheets. No significant differences in the *O*-Man microheterogeneity between WT and ΔPOMGnT1 N-Cdh were detected. Please note, *O*-Man region IV has not been reported before.

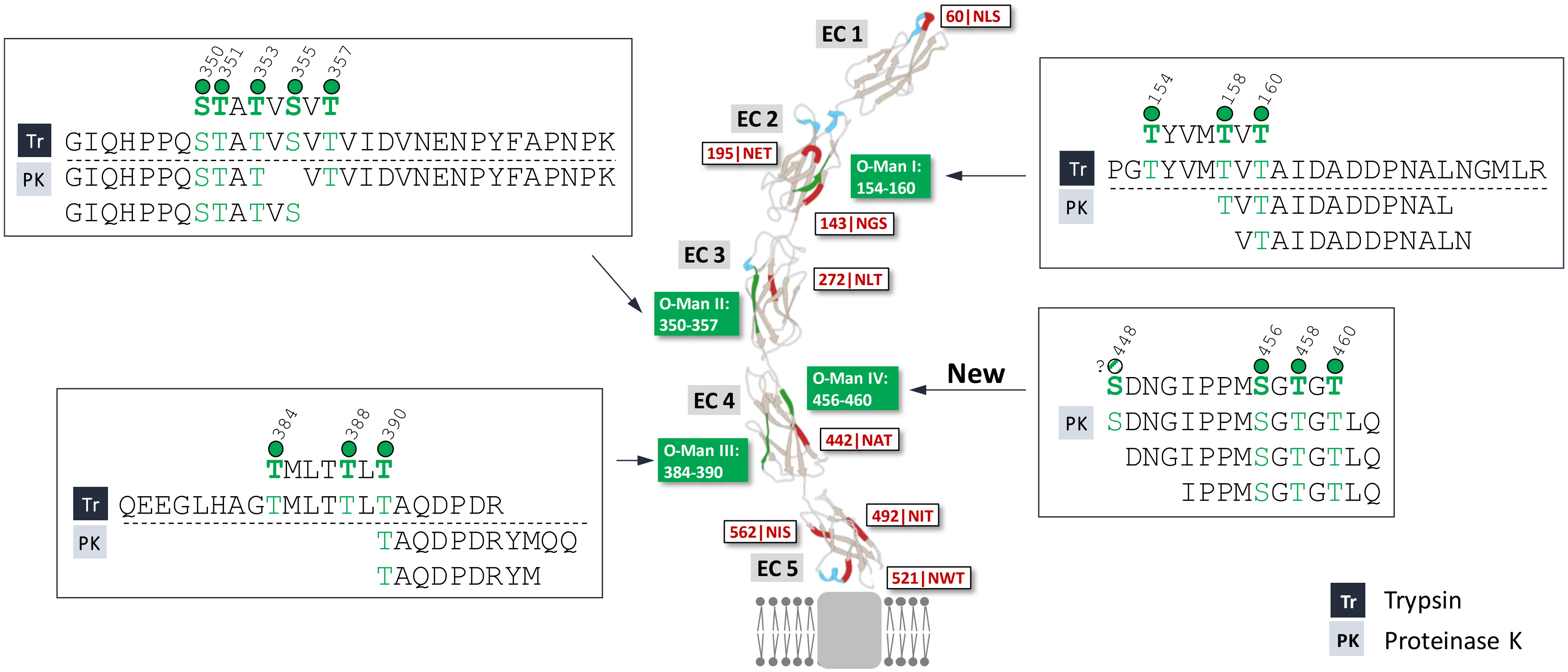

**Figure S15: Identified tryptic and proteinase-K-generated *O*-Man glycopeptides within human N-Cdh**. Occupied *O*-Man sites are indicated with a green circle (a white circle indicates presumably occupied sites).

#### Byonic Settings

| General Search Parameters (tryptic digest) | General Search Parameters (no-specific digest) |
|---|---|
| Proteolytic Cleavage Sites: RK (C-terminal) | Proteolytic Cleavage Sites: - |
| Digestion Specificity: Fully Specific | Digestion Specificity: Non specific |
| Missed Cleavages: 2 | Missed Cleavages: 0 |
| Precursor Mass Tolerance: 10 ppm | Precursor Mass Tolerance: 10 ppm |
| Fragmentation Type: QTOF/HCD | Fragmentation Type: QTOF/HCD |
| Fragment Mass Tolerance: 20 ppm | Fragment Mass Tolerance: 20 ppm |
| Recalibration (Lock Mass): 445.12003 | Recalibration (Lock Mass): 445.12003 |

*N*-Glycopeptide Search

Total Common Max: 1
Total rare Max: 2
Carbamidomethyl / +57.021464 @ C | fixed
Carbamidomethyl / +57.021464 @ NTerm | rare1
Deamidation / +0.984016 @ N, Q | rare1
Oxidation / +15.994915 @ M, W, Y | rare1
Dioxidation / +31.989829 @ W, Y | rare1
Dehydrated / -18.010565 @ S, T | rare1
Trp → Kynurenin / +3.994915 @ W | rare1
Phospho / +79.966331 @ S, T, Y | rare1
N-Glycan Search: N-glycan Database, Human | common1

*O-Man*-Glycopeptide Search

Total Common Max: 5
Total rare Max: 2
Carbamidomethyl / +57.021464 @ C | fixed
Carbamidomethyl / +57.021464 @ NTerm | rare1
Deamidation / +0.984016 @ N, Q | rare1
Oxidation / +15.994915 @ M, W, Y | rare1
Dioxidation / +31.989829 @ W, Y | rare1
Dehydrated / -18.010565 @ S, T | rare1
Trp → Kynurenin / +3.994915 @ W | rare1
Phospho / +79.966331 @ S, T, Y | rare1
O-Man-Glycan Search:
*Scenario I:* Hex(1) @ S, T | common5
*Scenario II:* HexNAc(1)Hex(1) @ S, T | common1; HexNAc(1)Hex(2) @ S, T | common1; HexNAc(2)Hex(1) @ S, T | common1; HexNAc(2)Hex(2) @ S, T | common1; HexNAc(2)Hex(3) @ S, T | common1; HexNAc(1)Hex(2)NeuAc(1) @ S, T | common1; HexNAc(2)Hex(3)NeuAc(1) @ S, T | common1

#### SI 5: Supporting Information of N-Cdh *N*-Glycomics

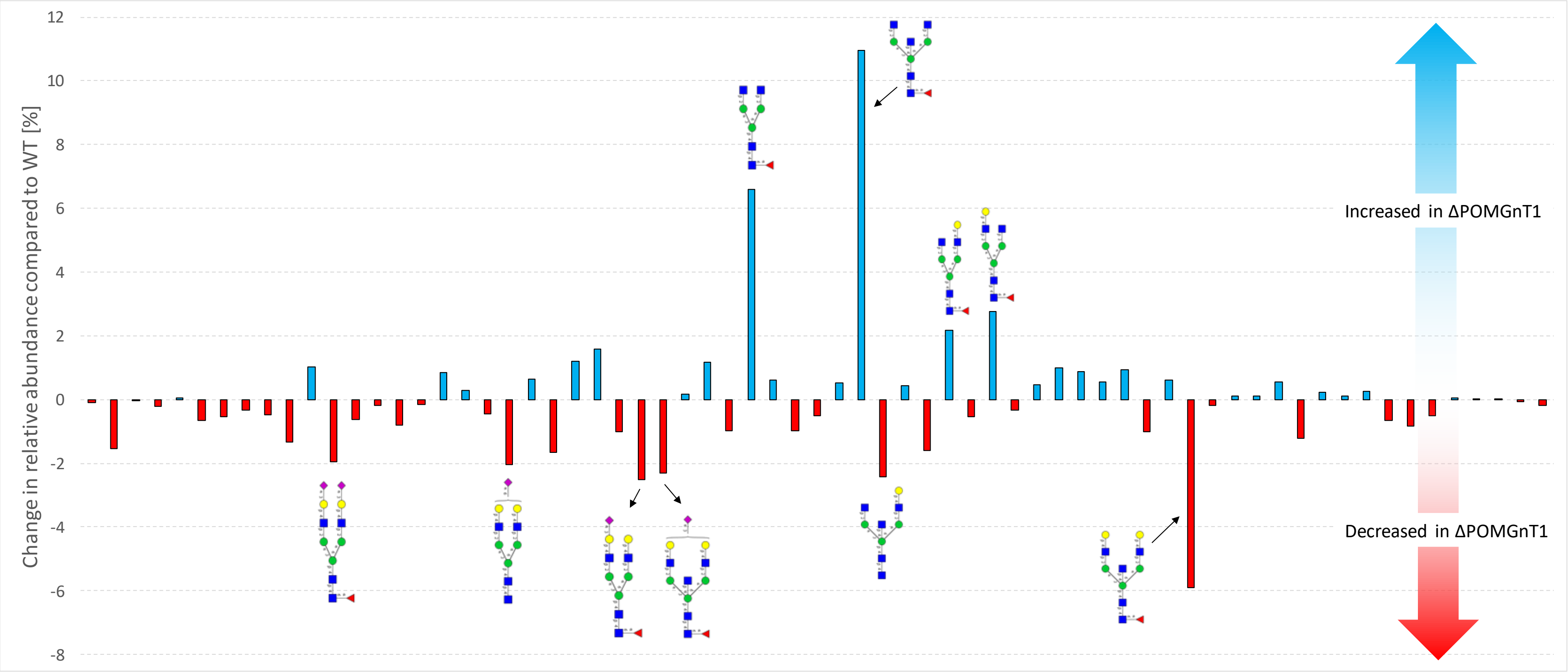

**Figure S16: xCGE-LIF-based extended *N*-glycan profiling of N-Cdh derived from wild-type (WT) and ΔPOMGnT1 HEK293T cells**. The change in relative abundance of N-Cdh *N*-glycan structures derived from ΔPOMGnT1 and WT HEK293T cells is shown as bar graphs. Blue bars indicate ΔPOMGnT1 N-Cdh *N*-glycan structures with an increased abundance compared to WT. Red bars indicate ΔPOMGnT1 N-Cdh *N*-glycan structures with a decreased abundance compared to WT. For *N*-glycan structures with the most significant changes (relative change >2%) is depicted. *N*-glycan structures were drawn with GlycoWorkbench Version 1.1, by following the guideline of the Consortium for Functional Glycomics. Further details on all identified and quantified *N*-glycan structures are provided in Table S18.

**Table S18: List of identified and quantified *N*-glycan structures on N-Cdh derived from wild-type (WT) and ΔPOMGnT1 HEK293T cells.**

| Peak Number | *N*-Glycan Name | N-Cdh from | | Delta (ΔPOMGnT1-WT) [%] |
|---|---|---|---|---|
| | | WT | ΔPOMGnT1 | |
| | | Rel. Sig. Intensity in [%] | | |
| 1 | Not Identified | 0.30 | 0.21 | -0.08 |
| 2 | FA2BG2SuS1(2,3)? | 3.14 | 1.61 | -1.53 |
| 3 | A4G4S4(2,3) | 0.19 | 0.18 | -0.01 |
| 4 | Not Identified | 0.38 | 0.15 | -0.22 |
| 5 | A2G2S2(2,6) | 0.11 | 0.15 | 0.04 |
| 6 | FA4G4S4(2,3) | 0.76 | 0.10 | -0.66 |
| 7 | Not Identified | 0.66 | 0.11 | -0.54 |
| 8 | FA2G2S2(2,6) | 0.34 | 0.00 | -0.34 |
| 9 | FA2G2S1(2,3)S1(2,6) | 0.98 | 0.51 | -0.47 |
| 10 | FA2G2S1(2,3)S1(2,6) | 1.74 | 0.39 | -1.35 |
| 11 | FA2BG2S1(2,3)S1(2,6) | 1.46 | 2.48 | 1.02 |
| 12 | FA2G2S2(2,3) | 2.43 | 0.48 | -1.96 |
| 13 | A2BG2Su? | 0.81 | 0.18 | -0.62 |
| 14 | A4G4S3(2,3) | 0.47 | 0.30 | -0.17 |
| 15 | FA4G4S3(2,3) | 0.79 | 0.00 | -0.79 |
| 16 | FA2BG2Su? | 4.06 | 3.91 | -0.15 |
| 17 | A3G3S2(2,3) | 0.71 | 1.55 | 0.84 |
| 18 | FA2G1S1(2,3)[6] | 0.34 | 0.62 | 0.27 |
| 19 | FA3G3S2(3) | 0.46 | 0.00 | -0.46 |
| 20 | A2G2S1(2,6) | 2.88 | 0.83 | -2.05 |
| 21 | FA2G1S1(2,3)[3] | 1.15 | 1.77 | 0.63 |
| 22 | Not Identified | 1.84 | 0.17 | -1.67 |
| 23 | Man5 & FA2G2S1(2,6) | 5.76 | 6.97 | 1.21 |
| 24 | A2G0 | 0.00 | 1.59 | 1.59 |
| 25 | FA2G2S1(2,3)[6] | 1.01 | 0.00 | -1.01 |
| 26 | FA2G2S1(2,3)[3] | 3.53 | 1.01 | -2.52 |
| 27 | FA2BG2S1(2,3) | 3.06 | 0.75 | -2.31 |
| 28 | FA4G4S2(2,3) | 0.00 | 0.15 | 0.15 |
| 29 | A2BG2 | 2.12 | 3.29 | 1.16 |
| 30 | Man6 | 9.08 | 8.10 | -0.98 |
| 31 | FA2G0 | 2.66 | 9.27 | 6.61 |
| 32 | Not Identified | 0.00 | 0.61 | 0.61 |
| 33 | Not Identified | 0.99 | 0.00 | -0.99 |
| 34 | Not Identified | 0.76 | 0.24 | -0.52 |
| 35 | Not Identified | 0.00 | 0.52 | 0.52 |
| 36 | FA2BG0 | 5.78 | 16.73 | 10.96 |
| 37 | A2BG1[6] | 2.95 | 0.51 | -2.44 |

| | | | | |
|---|---|---|---|---|
| 38 | Man7 | 1.80 | 2.24 | 0.44 |
| 39 | A2BG1[3] & FA3G3S1(2,3) | 1.73 | 0.12 | -1.61 |
| 40 | FA2G1[6] | 0.56 | 2.75 | 2.19 |
| 41 | Man5-A1G1 | 0.54 | 0.00 | -0.54 |
| 42 | FA2G1[3] | 0.82 | 3.59 | 2.77 |
| 43 | FA2BG1[6] & A2G2 | 9.28 | 8.94 | -0.33 |
| 44 | Not Identified | 0.10 | 0.56 | 0.46 |
| 45 | FA2BG1[3] | 1.86 | 2.86 | 1.00 |
| 46 | FA3[2,6]G1 | 0.18 | 1.06 | 0.88 |
| 47 | A2F1[6]G2 | 0.38 | 0.95 | 0.56 |
| 48 | FA3[2,6]G1 | 0.00 | 0.95 | 0.95 |
| 49 | FA2G2 | 3.03 | 2.01 | -1.02 |
| 50 | A3[2,6]G2 | 0.25 | 0.85 | 0.60 |
| 51 | FA2BG2 | 8.35 | 2.45 | -5.90 |
| 52 | FA2F1[6]G2 | 0.19 | 0.00 | -0.19 |
| 53 | FA2F1[3]G2 | 0.00 | 0.10 | 0.10 |
| 54 | FA3[2,6]G2 | 0.23 | 0.33 | 0.10 |
| 55 | FA3[2,6]G2 & FA4G1 | 0.30 | 0.85 | 0.55 |
| 56 | FA3[2,6]G2 | 1.21 | 0.00 | -1.21 |
| 57 | A3[2,6]G3 & A4G2 | 0.16 | 0.39 | 0.24 |
| 58 | FA3[2,4]G2 & FA3B[2,6]G2 | 0.49 | 0.59 | 0.10 |
| 59 | FA3B[2,6]G2 | 0.33 | 0.60 | 0.27 |
| 60 | FA3[2,6]G3 & FA4G2 | 1.22 | 0.57 | -0.65 |
| 61 | FA3B[2,6]G3 & FA3[2,4]G3 | 1.21 | 0.39 | -0.83 |
| 62 | FA3B[2,4]G3 | 1.07 | 0.55 | -0.52 |
| 63 | FA4G3 | 0.20 | 0.27 | 0.06 |
| 64 | FA4G3 | 0.10 | 0.10 | 0.00 |
| 65 | FA4BG3 | 0.18 | 0.21 | 0.04 |
| 66 | FA4G4 | 0.20 | 0.14 | -0.07 |
| 67 | FA4BG4 | 0.30 | 0.11 | -0.19 |
